# Consumption of processed foods impairs memory function through dietary advanced glycation end-products

**DOI:** 10.64898/2026.01.07.698065

**Authors:** Anna M. R. Hayes, Molly E. Klug, Madhav Sharma, Alicia E. Kao, Shan Sun, Edwin D. J. Lopez Gonzalez, Houming Zhu, Joshua C. Dent, Richard J. Clark, David R. Sell, Deepitha Nelson, Vincent M. Monnier, Linda Tsan, Jessica J. Rea, Arun Ahuja, Natalie Tanios, Isabella Gianatiempo, Mugil V. Shanmugam, Yedam Park, Kristie B. Yu, Elaine Y. Hsiao, Lindsey A. Schier, Anthony A. Fodor, Trent M. Woodruff, Sarah C. Shuck, Cornelius Gati, Bruce E. Herring, Melinda T. Coughlan, Scott E. Kanoski

**Author notes:** Corresponding author: Scott E. Kanoski, Current Address: 3616 Trousdale Parkway, AHF-252, Los Angeles, CA 90089-0372.

## Abstract

Consumption of processed foods is associated with dementia, obesity, and other negative health outcomes. Sustained heat treatment, a common food processing approach to enhance flavor, induces the chemical Maillard reaction that promotes the formation of dietary advanced glycation end-products (AGEs). The neurocognitive impacts of consuming dietary AGEs are poorly understood. Here we modeled an AGE-rich diet through heat treatment fed to rats during adolescence, a critical period of neural development, to mechanistically evaluate the long-term impact of early life dietary AGEs on behavioral and neural processes. Consuming the AGE-rich diet impaired hippocampal-dependent memory function and altered the gut microbiome without inducing obesity or nonspecific behavioral deficits. AGE-induced memory deficits were coupled with impaired hippocampal glutamatergic synaptic neurotransmission and altered expression in the synapse-pruning complement system. Hippocampal synaptic deficits likely result from direct AGE-complement interactions, as our extended studies reveal competitive antagonist action of AGEs on complement receptors. Memory impairments were prevented by administration of the AGE-inhibitor, alagebrium, and by supplementation with an AGE-inhibiting bacterial taxon, *Lactococcus lactis*, which was depleted in the heat-treated diet. These findings reveal a functional connection between early life dietary AGEs, the microbiome, and memory impairments, thus illuminating mechanisms through which food processing negatively impacts neurocognition.

## Main text

The dietary landscape has changed drastically throughout history, with an exponential increase in food processing since the industrial revolution^1^. Consumption of highly-processed foods (a.k.a., ‘ultra-processed foods’) is linked with metabolic, cardiovascular, and neurodegenerative disorders^2–6^. To date, research on the health implications of processed food consumption has predominantly relied on classification systems that categorize foods by different levels of ‘processing’, including the NOVA system that groups foods into four categories^7–13^. A major limitation of such classification systems is that they do not isolate the specific food processing characteristics that underlie negative health impacts, thereby limiting insights into underlying biological mechanisms. To bridge this gap, we implemented a reductionist approach by utilizing a diet model that incorporates a singular common food processing approach – heat treatment^14^. Sustained thermal treatment of food, particularly under conditions of high heat (150 °C or higher), low moisture, and high pH, induces a series of chemical reactions in foods referred to as the ‘Maillard reaction’^15^. In addition to enhancing the flavor of food, the Maillard reaction promotes the formation of pro-inflammatory compounds known as advanced glycation end-products (AGEs)^16,17^. Dietary AGEs have been linked with dementia^18^, metabolic syndrome^19–21^, and compromised gut barrier integrity^22–24^, yet the underlying drivers of these impacts remain poorly understood. Here, we utilized a rodent model of early life AGE-rich diet exposure to uncover the effects of sustained heat treatment of food on neurocognition. Our results reveal novel gut bacterial, neurobiological, and molecular mechanisms through which a common food processing approach disrupts memory function independent of obesity or disordered metabolism.

### Early life consumption of a heat-treated diet high in advanced glycation end-products (AGEs) increases physiological AGE levels

Male Sprague Dawley rats were maintained during the juvenile-adolescent period (from postnatal days [PN] 26-60) on either a control (non-heat-treated) standard purified rodent chow diet, or the same diet, but heat-treated to increase levels of AGEs (diet heated for 1 h at 160 °C in a forced-draft oven; Extended Data Fig. 1a-c) (Fig. 1a). The heat-treated AGE-rich diet resulted in elevated levels of free AGEs (i.e., AGEs that are not bound to protein), including methylglyoxal-derived hydroimidazolone (MG-H1), carboxymethyllysine (CML), and carboxyethyllysine (CEL), in serum (Fig. 1b-e). These elevated serum AGEs were accompanied by higher AGE levels in urine (Fig. 1f-i; including increased carboxyethylguanosine [CEG]). Such findings corroborate previous research indicating that ingestion of exogenous AGEs is associated with increased levels of free AGEs in the body in both adult rats and humans^25,26^.

**Figure 1.**
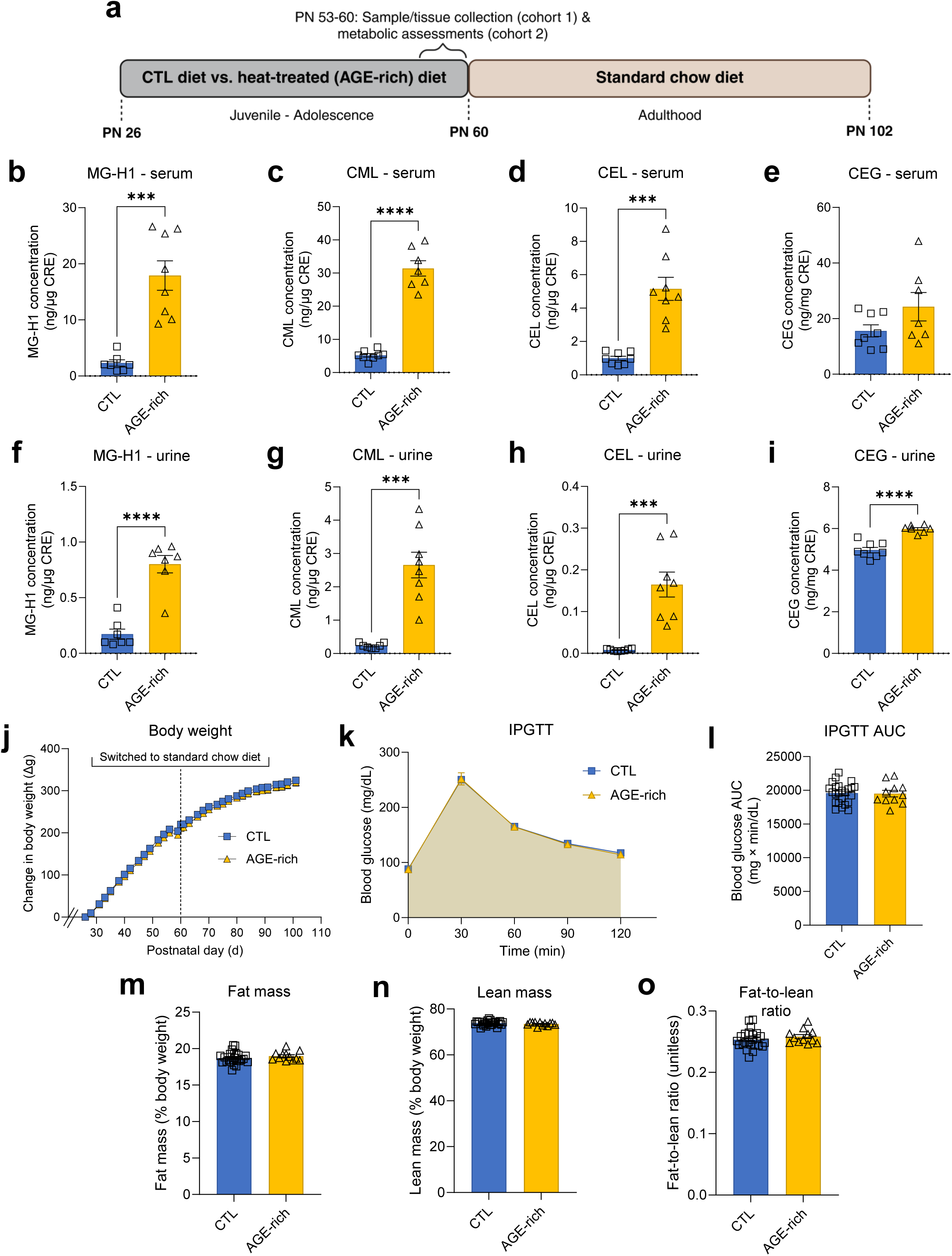
Consumption of a heat-treated diet during early life leads to elevated levels of circulating and excreted advanced glycation end-products (AGEs) without altering body weight trajectory, glucose tolerance, or body composition. **a,** Experimental timeline illustrating early life diet exposure of either a control diet (CTL, AIN-93G) or heat-treated AGE-rich diet (AIN-93G diet, heated for 1 h at 160 °C in a forced-draft oven) from postnatal days (PN) 26-60. One cohort of rats was euthanized on PN 56 for sample/tissue collection. For a second cohort, metabolic assessments (glucose tolerance, body composition) were performed from PN 53-60. On PN 60, both groups in the second cohort were switched to standard chow until the end of the experiment at PN 102. **b,** Concentration of MG-H1 in serum in the postprandial state on PN 56 in rats that received the CTL vs. AGE-rich diet during early life (n=7 CTL, n=8 AGE-rich; Welch’s t-test, P=0.0005). **c,** Concentration of CML in serum in the postprandial state on PN 56 in rats receiving the CTL vs. AGE-rich diet during early life (n=8 CTL, n=7 AGE-rich; Welch’s t-test, P<0.0001). **d,** Concentration of CEL in serum in the postprandial state on PN 60 in rats receiving the CTL vs. AGE-rich diet during early life (n=8 CTL, n=8 AGE-rich; two-sample Kolmogorov-Smirnov test, P=0.0002). **e,** Concentration of CEG in serum in the postprandial state on PN 56 in rats receiving the CTL vs. AGE-rich diet during early life (n=8 CTL, n=7 AGE-rich; unpaired two-tailed t-test, P=0.13). The elevated levels of MG-H1, CML, and CEL in rats receiving the AGE-rich diet indicates that they are absorbing AGEs from the diet. **f,** Concentration of MG-H1 in urine collected during the dark cycle on PN 53 in rats that received the CTL vs. AGE-rich diet during early life (n=7 CTL, n=7 AGE-rich; unpaired two-tailed t-test, P<0.0001). **g,** Concentration of CML in urine collected during the dark cycle on PN 53 in rats that received the CTL vs. AGE-rich diet during early life (n=7 CTL, n=8 AGE-rich; Welch’s t-test, P=0.0004). **h,** Concentration of CEL in urine collected during the dark cycle on PN 53 in rats that received the CTL vs. AGE-rich diet during early life (n=8 CTL, n=8 AGE-rich; two-sample Kolmogorov-Smirnov test, P=0.0002). **i,** Concentration of CEG in urine collected during the dark cycle on PN 53 in rats that received the CTL vs. AGE-rich diet during early life (n=8 CTL, n=7 AGE-rich; unpaired two-tailed t-test, P<0.0001). The elevated levels of MG-H1, CML, CEL, and CEG in rats receiving the AGE-rich diet indicates that they are both absorbing and excreting higher amounts of AGEs – which are likely from the diet. **j,** Change in body weight over time in rats receiving the CTL vs. AGE-rich diets from PN 26-60 followed by standard chow (n=24 CTL, n=12 AGE-rich; 2-way ANOVA, diet group [P=0.5159], time [P<0.0001], diet group × time [P>0.9999]). **k,** Blood glucose values over time from IPGTT performed on PN 59 (n=24 CTL, n=11 AGE rich; 2-way ANOVA, diet group [P=0.8294], time [P<0.0001], diet group × time [P=0.9912]). **l,** Blood glucose AUC from IPGTT performed on PN 59 (n=24 CTL, n=11 AGE-rich; unpaired two-tailed t-test, P=0.89). **m,** Body fat mass on PN 60 (n=24 CTL, n=12 AGE-rich; unpaired two-tailed t-test, P=0.31). **n,** Body lean mass on PN 60 (n=24 CTL, n=12 AGE-rich; unpaired two-tailed t-test, P=0.34). **o,** Fat-to-lean ratio on PN 60 (n=24 CTL, n=12 AGE-rich; unpaired two-tailed t-test, P=0.33). The lack of differences in body weight, glucose tolerance, and body composition suggest that the AGE-rich diet is not altering body weight and glucose regulation. Error bars represent standard error of the mean (SEM). *P<0.05, **P<0.01, ***P<0.001, ****P<0.0001. All n’s indicate number of rats per group. Additional details about the statistical analyses for each subpanel can be found in Supplementary Table S1. AGE, advanced glycation end-product; ANOVA, analysis of variance; AUC, area under the curve; CEG, carboxyethylguanosine; CEL, carboxyethyllysine; CML, carboxymethyllysine; CTL, control [diet]; IPGTT, intraperitoneal glucose tolerance test; MG-H1, methylglyoxal-derived hydroimidazolone; PN, postnatal day.

Despite having increased levels of circulating AGEs, rats consuming the AGE-rich diet in early life did not exhibit differences in body weight trajectory, glucose tolerance, or body composition compared to controls (Fig. 1j-o). The pelletization process used to produce the rodent feed pellets, another form of food processing, also did not alter any of these outcomes, as indicated by a complete lack of differences between groups receiving non-heat-treated diets that were either pelleted or powdered (Extended Data Fig. 1d, f-k). While the group consuming the AGE-rich diet consumed fewer calories during the experimental diet period than controls (Extended Data Fig. 1e), the lack of differences in body weight and body composition between groups suggest that this reduction in energy intake was compensated for by reductions in energy expenditure in the AGE-rich group.

### AGE-rich diet consumption during early life impairs memory function

Previous research identified the adolescent developmental stage as an influential period of neurocognitive development, which is emphasized by early life dietary factors having long-lasting impacts on memory function^27–30^. Here we evaluated the effects of early life exposure to the AGE-rich diet on cognitive function during adulthood, following a switch of both groups to standard chow at PN 60 to confine the experimental diet exposure period to early life (Fig. 2a). Rats that consumed the heat-treated diet in early life showed impaired memory function in both the novel object in context task (Fig. 2b-d and Extended Data Fig. 2a-d) and novel location recognition task (Fig. 2 e-g and Extended Data Fig. 2e-g), which assess contextual episodic memory and spatial recognition memory, respectively. Rats consuming the heat-treated diet during early life did not exhibit deficits in the novel object recognition task (Fig. 2h-j and Extended Data Fig. 2h-j). Early life heat-treated AGE-rich diet consumption also did not alter anxiety-like behavior or locomotor activity, as indicated through the zero maze and open field tests (Fig. 2k-p and Extended Data Fig. 2k-n).

**Figure 2.**
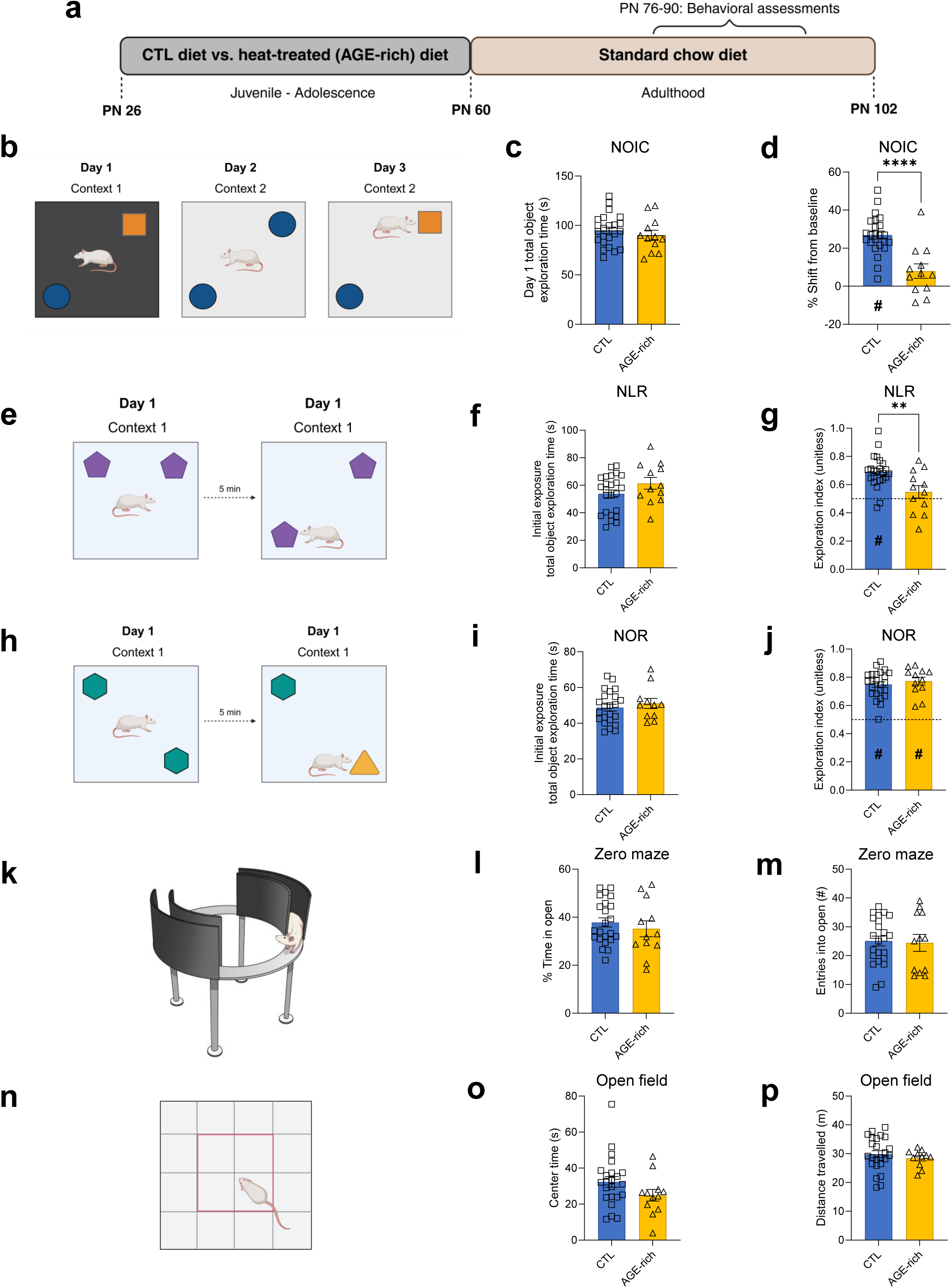
Consumption of a heat-treated AGE-rich diet during early life leads to hippocampal-dependent memory impairments in adulthood, without impacting perirhinal cortex-dependent memory, anxiety-like behavior, or locomotor activity. **a**, Experimental timeline illustrating early life diet exposure of either a control diet (CTL, AIN-93G) or heat-treated AGE-rich diet from postnatal days (PN) 26-60. On PN 60, both groups in the second cohort were switched to standard chow until the end of the experiment. Behavioral assessments were performed from PN 76-90. **b**, Illustration of the NOIC procedure. **c**, Total object exploration time during NOIC day 1 (n=23 CTL, n=12 AGE-rich; unpaired two-tailed t-test, P=0.41). A difference in time spent exploring the objects during this initial exposure would indicate that the interpretation of object exploration behavior during later stages could be confounded by initial exposure encoding; since there are no differences here we can rule out this potential confound. **d**, NOIC percent shift from baseline object exploration (n=24 CTL, n=12 AGE-rich; unpaired two-tailed t-test, P<0.0001; one-sample t-test for difference from 0 [chance], P<0.0001 CTL, P=0.06 AGE-rich). A positive percent shift from baseline indicates that the rats preferentially explored the novel object in context on NOIC day 3 and had intact hippocampal-dependent memory function. **e**, Illustration of the NLR procedure. **f**, Total object exploration time during the initial exposure period for NLR (n=24 CTL, n=12 AGE-rich; unpaired two-tailed t-test, P=0.14). **g**, NLR object exploration index during the test exposure period (n=23 CTL, n=12 AGE-rich; unpaired two-tailed t-test, P=0.004; one-sample t-test for difference from 0.5 [chance], P<0.0001 CTL, P=0.28 AGE-rich). An exploration index higher than 0.5 (chance) indicates that the rats recognized the object moved to a novel location and thus have intact hippocampal-dependent spatial recognition memory. **h**, Illustration of the NOR procedure. **i**, Total object exploration time during the initial exposure period for NOR (n=23 CTL, n=11 AGE-rich; unpaired two-tailed t-test, P=0.50). **j**, NOR object exploration index during the test exposure period (n=24 CTL, n=12 AGE-rich; unpaired two-tailed t-test, P=0.49; one-sample t-test for difference from 0.5 [chance], P<0.0001 CTL, P<0.0001 AGE-rich). An exploration index higher than 0.5 (chance) indicates that the rats recognized the novel object and thus have intact perirhinal cortex-dependent object novelty recognition. **k**, Illustration of the zero maze test of anxiety-like behavior. **l**, Percent time in the open arm zones during the zero maze test (n=23 CTL, n=12 AGE-rich; unpaired two-tailed t-test, P=0.47). **m**, Number of entries into the open arm zones during the open maze test (n=23 CTL, n=12 AGE-rich; unpaired two-tailed t-test, P=0.83). **n**, Illustration of the open field test of anxiety-like behavior and locomotor activity. **o**, Time spent in the center zone during the open field test (n=23 CTL, n=12 AGE-rich; unpaired two-tailed t-test, P=0.12). More time spent in the center is indicative of decreased anxiety-like behavior. **p**, Distance travelled during the open field test, which is indicative of locomotor activity (n=24 CTL, n=11 AGE-rich; Welch’s t-test, P=0.35). Error bars represent standard error of the mean (SEM). *P<0.05, **P<0.01, ***P<0.001, ****P<0.0001. # indicates statistically significant difference from chance (P<0.05), which was 0.0 for NOIC and 0.5 for NLR and NOR. All n’s indicate number of rats per group. Additional details about the statistical analyses for each subpanel can be found in Supplementary Table S1. AGE, advanced glycation end-product; ANOVA, analysis of variance; CTL, control [diet]; NLR, novel location recognition; NOIC, novel object in context; NOR, novel object recognition; PN, postnatal day.

### AGE-rich diet impairs hippocampus glutamatergic neurotransmission and alters complement system targets

The AGE-rich diet was associated with memory impairments in the novel object in context and novel location procedures, both of which are dependent on an intact hippocampus^31–34^. However, memory performance was not affected in the novel object recognition task using procedures that are not hippocampal-dependent^35^. These findings suggest that hippocampal-dependent memory processes are particularly vulnerable to the effects of an AGE-rich diet during early life. To understand how consumption of a heat-treated diet high in AGEs is ultimately impairing hippocampal-dependent memory function, we performed field recordings in dorsal hippocampus slices from rats that received either the AGE-rich diet or control diet during early life. Results showed that rats receiving the AGE-rich diet had decreased field-to-volley ratio across all fiber volley amplitudes tested (Fig. 3a-b), indicating a robust impairment in glutamatergic transmission, and by extension, in glutamatergic synapse development/plasticity.

**Figure 3.**
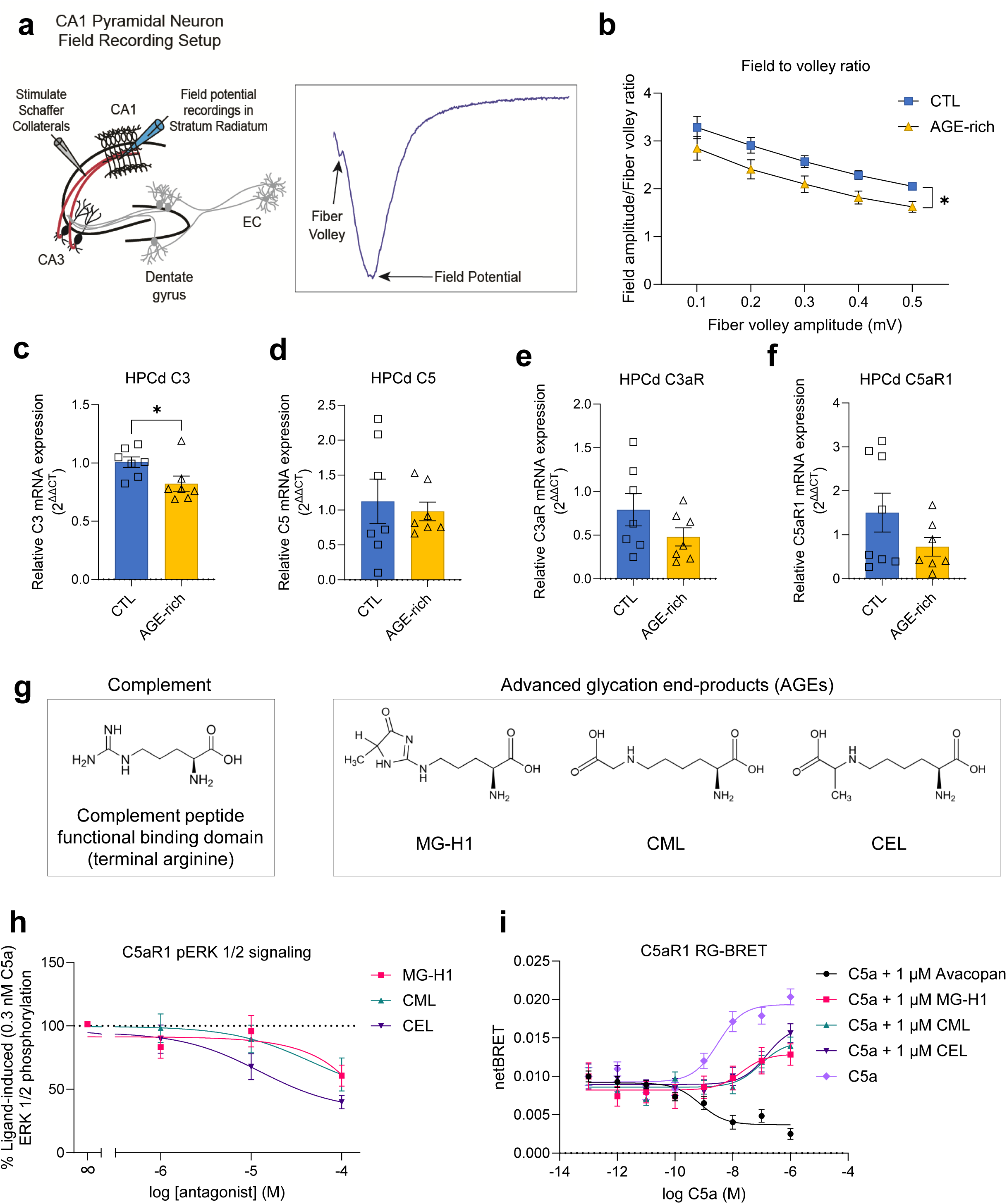
Early life consumption of an AGE-rich diet disrupts glutamatergic neurotransmission and dorsal hippocampus complement system activation. **a**, Illustration of the electrophysiology field potential recording setup. **b**, Ratio of field to volley from electrophysiology field potential recordings of dorsal hippocampus (HPCd) brain tissue slices in rats after 30-d early life AGE-rich diet (n=14 CTL tissue slices, n=15 AGE-rich tissue slices; 2-way repeated measures ANOVA with factors of diet group, fiber volley amplitude [repeated measure], and diet group × fiber volley interaction; P=0.04 diet group, P<0.0001 fiber volley amplitude, P=0.99 diet group × fiber volley amplitude). **c**, mRNA expression of complement component 3 (C3) in the HPCd (n=7 CTL, n=7 AGE-rich; unpaired two-tailed t-test, P=0.04). **d**, mRNA expression of complement component 5 (C5) in the HPCd (n=7 CTL, n=7 AGE-rich; unpaired two-tailed t-test, P=0.68). **e**, mRNA expression of C3aR in the HPCd (n=7 CTL, n=7 AGE-rich; unpaired two-tailed t-test, P=0.17). **f**, mRNA expression of C5aR1 in the HPCd (n=8 CTL, n=7 AGE-rich; unpaired two-tailed t-test, P=0.16). **g**, Structures of the peptide required for complement system receptor activation and of AGEs elevated in the heat-treated diet. **h**, C5aR1 pERK 1/2 signaling in response to MG-H1, CML, and CEL (smoothed lines indicate nonlinear least squares fit for each treatment). **i**, C5aR1 Receptor-G protein-Bioluminescence Resonance Energy Transfer (RG-BRET) assay for binding affinities between C5a and MG-H1, CML, CEL, or the C5aR1-selective antagonist avacopan (smoothed lines indicate nonlinear least squares fit for each treatment). Error bars represent standard error of the mean (SEM). *P<0.05. All n’s indicate number of rats or slices per group. Additional details about the statistical analyses for each subpanel can be found in Supplementary Table S1. AGE, advanced glycation end-product; ANOVA, analysis of variance; C3, complement component 3; C3aR, receptor for complement component 3a; C5, complement component 5; C5aR1, receptor for complement component 5a; CA1, Cornu Ammonis subfield 1; CA3, Cornu Ammonis subfield 3; EC, entorhinal cortex; CEL, carboxyethyllysine; CML, carboxymethyllysine; CTL, control [diet]; HPCd, dorsal hippocampus; MG-H1, methylglyoxal-derived hydroimidazolone; pERK, phospho-extracellular signal-regulated kinase; PN, postnatal day; RG-BRET, Receptor-G protein-Bioluminescence Resonance Energy Transfer.

To further probe how glutamatergic synaptic transmission was being compromised, we examined hippocampal expression of various targets that are associated with hippocampal-dependent memory, including the endogenous AGE receptor (RAGE), nuclear factor kappa-light-chain-enhancer of activated B cells (NFĸB), and postsynaptic density protein 95 (PSD-95) in rats given a control or AGE-rich diet^36–41^. We also examined expression of the complement system components, which have emerged as key players for synaptic pruning in the central nervous system (particularly the hippocampus), with implications in brain development as well as neurodegenerative diseases such as Alzheimer’s^42,43^. Previous research has indicated that consumption of a heat-treated AGE-rich diet disrupts complement system function and accompanying levels of complement components 3 and 5 (C3 and C5) in the periphery in rodents^22^. Our present results revealed that the AGE-rich diet did not impact hippocampal expression of RAGE or NFĸB, with a trend observed for a reduction in PSD95 (Extended Data Fig. 3a-c). For the complement system components, the AGE-rich diet yielded a significant reduction in expression of C3 in the hippocampus, whereas, while not statistically significant, the AGE-rich diet also produced trending reductions in C5, C3aR, and C5aR1 (Fig. 3c-f). These results highlight interactions with the complement system as a new mechanism via which AGEs could be altering synaptic signaling in the brain.

### AGEs are competitive antagonists of complement system receptors

Given our results indicating that dietary AGEs negatively impact complement signaling in the dorsal hippocampus, and that key structural characteristics of AGEs are strikingly similar to the complement peptide functional binding domain (i.e., involving a terminal arginine; Fig. 3g)^44^, we tested whether AGEs that were elevated in the heat-treated diet (MG-H1, CML, and CEL) affect phosphorylation of ERK 1/2 induced by complement 5 (C5a) in cell culture. Results revealed that all three AGEs reduced C5a ligand-induced phosphorylation of ERK 1/2 (Fig. 3h). Next, to evaluate whether these functional interactions are mediated directly by complement receptor binding, we measured the binding affinities of MG-H1, CML, and CEL for the complement system receptor C5aR1 using a Receptor-G protein-Bioluminescence Resonance Energy Transfer (RG-BRET) cell-based signaling assay^45^. Consistent with the ERK cell culture data, results revealed that all three AGEs exhibited antagonism function for C5aR1 relative to C5a alone in the RG-BRET assay (Fig. 3i).

Our results reveal that key AGEs molecularly interact with complement system receptors. We hypothesize that free AGEs with low molecular weights (<400 Da) – which includes MG-H1, CML, and CEL – are able to pass through the blood-brain-barrier due to their relatively small size^46,47^, a hypothesis supported by findings that isotopically-labelled dietary CML accumulated in the brain in mice^48^. This permeability would enable AGEs to act within the hippocampus to suppress complement signaling, and as proxy, to impair synaptic plasticity and associated memory. Since complement receptors are expressed throughout the body, the connection between AGEs and the complement system could also have implications on numerous physiological and neurobiological processes, such as immune function, synaptic pruning, and metabolism^42,43,49^.

Overall, our findings reveal that free AGEs, including those elevated in a heat-treated diet, can directly interact with complement system receptors, as confirmed through two distinct and complementary assays probing functional AGE-complement system interactions. We hypothesize that dietary AGEs engage hippocampal complement receptors to disrupt synaptic development/plasticity, and by extension, memory function.

### Dietary AGE-associated memory impairments are reversed by an AGE-inhibiting drug

Because sustained heat treatment can alter various physicochemical properties in foods (e.g., moisture content, levels of heat-labile vitamins), our initial experiments were unable to directly establish whether dietary AGEs themselves are causally related to the hippocampal-dependent memory impairments observed. Thus, in another set of experiments, we subjected rats receiving the heat-treated AGE-rich diet or control diet to either the AGE-inhibitor drug alagebrium (ALA)^22,50^ in drinking water (1 mg/kg/day) or to drinking water alone during the same PN 26-60 exposure period (Fig. 4a and Extended Data Fig. 4a-b). Results showed that ALA prevented hippocampal-dependent memory impairments in the novel object in context task in rats receiving the heat-treated diet but had no effect in control rats (Fig. 4b-c and Extended Data Fig. 4c-d). ALA had no impacts on perirhinal cortex-dependent memory performance in the novel recognition test (Fig. 4d-e and Extended Data Fig. 4e). Overall, these findings indicate that the hippocampal-dependent memory impairments and associated changes in hippocampal glutamatergic and complement signaling resulting from heat-treated diet consumption are due to the presence of AGEs in this diet.

**Figure 4.**
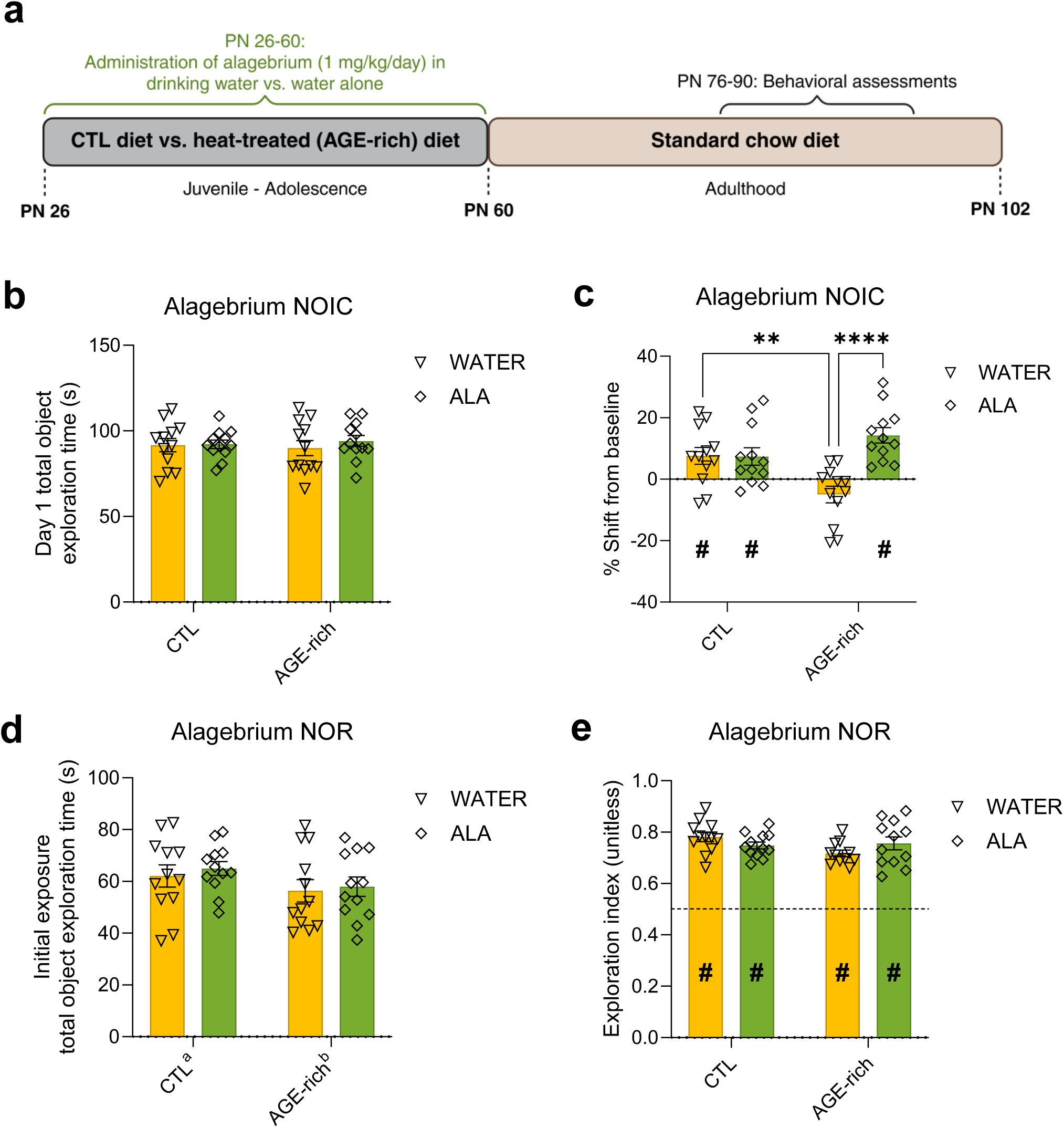
Administration of the AGE-inhibitor drug alagebrium (ALA) in early life prevents dietary AGE-induced memory impairments. **a**, Experimental timeline for ALA experiments. **b**, Total object exploration time during NOIC day 1 (n=12 CTL w/water, n=12 CTL w/ALA, n=12 AGE-rich w/water, n=12 AGE-rich w/ALA; 2-way ANOVA with factors of diet group, drug [water vs. ALA], and diet group × drug interaction; P=0.57 diet group, P=0.98 drug, P=0.53 diet group × drug). **c**, NOIC percent shift from baseline object exploration (n=12 CTL w/water, n=12 CTL w/ALA, n=12 AGE-rich w/water, n=12 AGE-rich w/ALA; 2-way ANOVA with factors of diet group, drug [water vs. ALA], and diet group × drug interaction; P=0.0015 diet group, P=0.32 drug, P=0.002 diet group × drug; post-hoc comparisons with Fisher’s LSD – P=0.97 CTL water vs. ALA, P<0.0001 AGE-rich water vs. ALA, P=0.004 Water CTL vs. AGE-rich, P=0.09 ALA CTL vs. AGE-rich; one-sample t-test for difference from 0 [chance], P=0.02 CTL w/water, P=0.02 CTL w/ALA, P=0.09 AGE-rich w/water, P<0.0001 AGE-rich w/ALA). **d**, Total object exploration time during the initial exposure period for NOR (n=12 CTL w/water, n=12 CTL w/ALA, n=12 AGE-rich w/water, n=12 AGE-rich w/ALA; 2-way ANOVA with factors of diet group, drug [water vs. ALA], and diet group × drug interaction; P=0.04 diet group, P=0.63 drug, P=0.83 diet group × drug). **e**, NOR object exploration index during the test exposure period (n=12 CTL w/water, n=12 CTL w/ALA, n=11 AGE-rich w/water, n=12 AGE-rich w/ALA; mixed effects analysis with factors of diet group, drug [water vs. ALA], and diet group × drug interaction; P=0.87 diet group, P=0.14 drug, P=0.06 diet group × drug; one-sample t-test for difference from 0.5 [chance], P<0.0001 CTL w/water, P<0.0001 CTL w/ALA, P<0.0001 AGE-rich w/water, P<0.0001 AGE-rich w/ALA). Error bars represent standard error of the mean (SEM).**P<0.01, ****P<0.0001. # indicates statistically significant difference from chance (P<0.05), which was 0.0 for NOIC and 0.5 for NOR. All n’s indicate number of rats per group. Additional details about the statistical analyses for each subpanel can be found in Supplementary Table S1. AGE, advanced glycation end-product; ALA, alagebrium; ANOVA, analysis of variance; CTL, control [diet]; NOIC, novel object in context; NOR, novel object recognition; PN, postnatal day.

### A heat-treated AGE-rich diet suppresses *Lactococcus* abundance in the food, as well as in the gut

Given evidence linking dietary AGEs with the gut microbiome^22–24,51^, we examined gut microbiota composition in rats receiving the heat-treated AGE-rich diet vs. the control diet. Results identified significant differences in microbial composition, revealed by PCoA in fecal samples collected at PN 56 (during the experimental diet period; Fig. 5a and Extended Data Fig. 5.1a-f). The sole most striking difference in microbial taxa was observed for the relative abundance of *Lactococcus* (genus), such that it was exponentially higher in control rats (Fig. 5b; relative abundances for other taxa shown in Extended Data Fig. 5.1g-l). Furthermore, the relative abundance of *Lactococcus* was directly correlated with the rats’ memory performance in the novel object in context task (Fig. 5c; additional correlations shown in Extended Data Fig. 5.1m-p). No differences in Shannon index, an indicator of alpha-diversity, were observed (Extended Data Fig. 5.1q-y).

**Figure 5.**
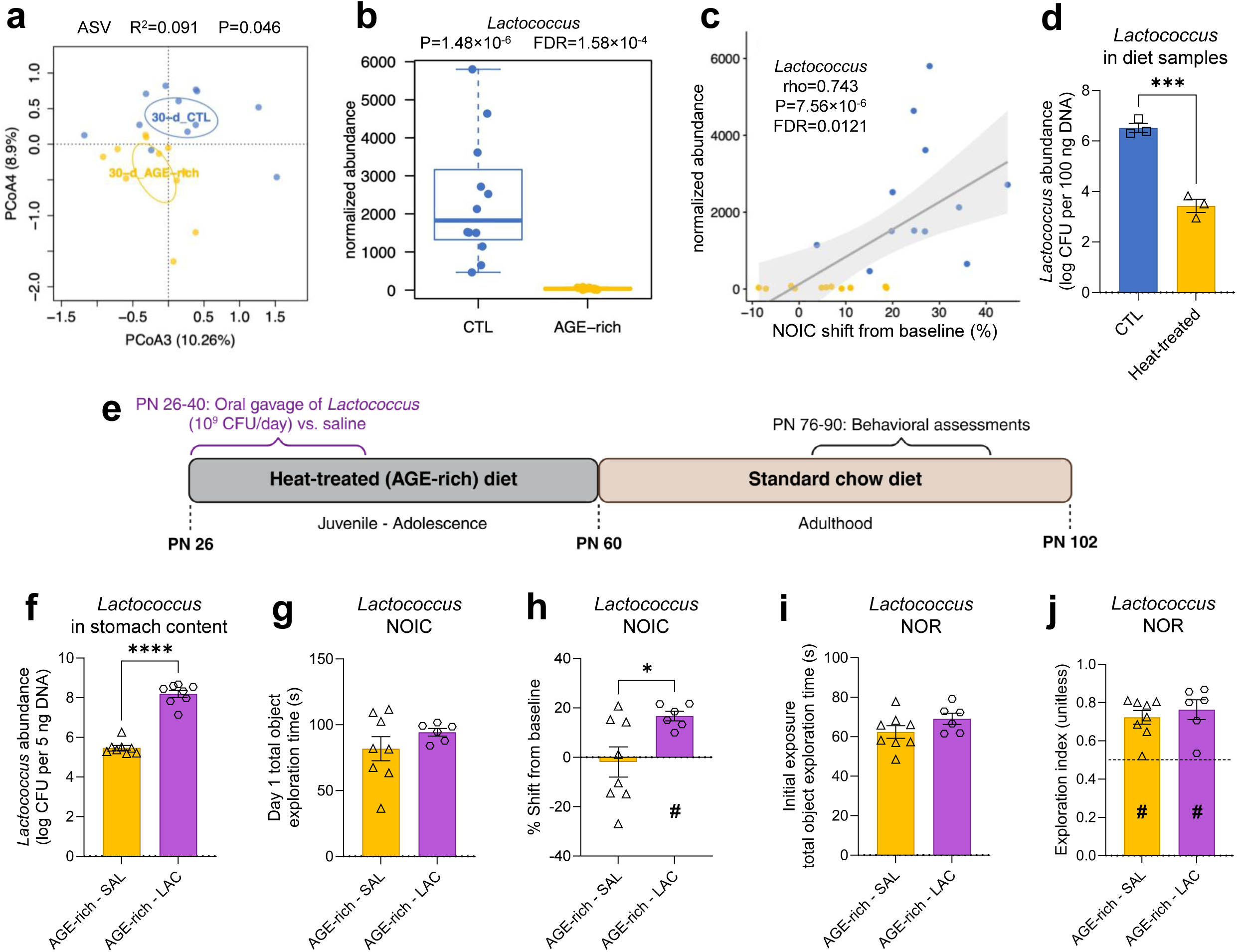
Consumption of a heat-treated AGE-rich diet in early life drives changes in the gut microbial taxon *Lactococcus*, and *Lactococcus* supplementation with an AGE-rich diet prevents the development of memory impairments. **a,** Principal coordinate analysis plot of amplicon sequence variant abundances in fecal samples from rats after 30-day consumption of the AGE-rich diet (n=12 CTL, n=12 AGE-rich; Bray-Curtis dissimilarity, coordinates 3 and 4; R^2^=0.09, P=0.046). **b**, Fecal *Lactococcus* abundance (n=12 CTL, n=12 AGE-rich; non-parametric Wilcoxon rank sum test; P=1.48×10^-6^, FDR=1.58×10^-4^). **c**, Correlation between NOIC performance (percent shift from baseline) and *Lactococcus* abundance (Spearman’s correlation, rho=0.743, P=7.56×10^-6^, FDR=0.0121). **d**, *Lactococcus lactis* subspecies *lactis* abundance in diet samples (n=3 CTL diet, n=3 AGE-rich diet; unpaired two-tailed t-test, P=0.0006). **e**, Experimental timeline for *Lactococcus* supplementation experiments. **f**, *Lactococcus lactis* subspecies *lactis* abundance in stomach content 2 h post-meal (n=8 AGE-rich - SAL, n=8 AGE-rich - LAC; unpaired two-tailed t-test, P<0.0001). **g**, Total object exploration time during NOIC day 1 (n=8 AGE-rich - SAL, n=6 AGE-rich - LAC; unpaired two-tailed t-test, P=0.28). **h**, NOIC percent shift from baseline object exploration (n=8 AGE-rich - SAL, n=6 AGE-rich - LAC; Welch’s t-test, P=0.02; one-sample t-test for difference from 0 [chance], P=0.77 AGE-rich - SAL, P=0.0003 AGE-rich - LAC). **i**, Total object exploration time during the initial exposure period for NOR (n=8 AGE-rich - SAL, n=6 AGE-rich - LAC; unpaired two-tailed t-test, P=0.16). **j**, NOR object exploration index during the test exposure period (n=8 AGE-rich - SAL, n=6 AGE-rich - LAC; unpaired two-tailed t-test, P=0.52; one-sample t-test for difference from 0.5 [chance], P=0.0004 AGE-rich - SAL, P=0.004 AGE-rich - LAC). Error bars represent standard error of the mean (SEM).*P<0.05, ***P<0.001, ****P<0.0001. # indicates statistically significant difference from chance (P<0.05), which was 0.0 for NOIC and 0.5 for NOR. All n’s indicate number of rats or samples per group. Additional details about the statistical analyses for each subpanel can be found in Supplementary Table S1. AGE, advanced glycation end-product; ANOVA, analysis of variance; ASV, amplicon sequence variant; CFU, colony forming unit; CTL, control [diet]; FDR, false discovery rate; LAC, *Lactococcus*; NOIC, novel object in context; NOR, novel object recognition; PN, postnatal day; SAL, saline [vehicle].

Typically used as a starter culture in dairy products, *Lactococcus* is a lactic acid bacterium that has not yet been explicitly identified as a long-term colonizing resident of the gut microbiota in rodents or humans, yet is strikingly similar in function to commonly-consumed probiotics in humans, such as *Lactobacillus*^52–55^. Our results confirmed that the heat-treated diet had significantly (and exponentially) lower abundance of *Lactococcus* (more specifically, *Lactococcus lactis* subspecies *lactis*) compared to the control diet (Fig. 5d). *Lactococcus lactis* is likely presented to the gut directly from the control diet due to the large proportion of the dairy protein casein in this diet. Heating this diet at the Maillard reaction-inducing conditions we used, however, yielded a form of sterilization that reduced *Lactococcus lactis* microbial abundance.

### *Lactococcus* supplementation offsets dietary AGE-induced memory deficits

To test whether *Lactococcus lactis* directly interacts with AGEs to influence cognition, we provided *Lactococcus lactis* subspecies *lactis* as a supplement via oral gavage (1 × 10^9^ CFU/day suspended in 0.9% w/v saline) from PN 26-40 to rats receiving the heat-treated AGE-rich diet, with control rats receiving the AGE-rich diet coupled with gavages of saline (Fig. 5e and Extended Data Fig. 5.2a-b). While both groups of rats remained on the AGE-rich diet until PN 60, the gavages were only performed from PN 26-40 as previous research has identified this timeframe as a sensitive window of hippocampal vulnerability to dietary influences^56^. To verify that our supplementation altered *Lactococcus* levels within the body, we confirmed that increased *Lactococcus lactis* abundance was observed in the stomach 1 h post-gavage (Fig. 5f). Although no differences were found in colonic levels of *Lactococcus lactis* 1 h post-gavage (Extended Data Fig. 5.2c), it is highly probable that the bacteria had not yet transited to this distal region of the gastrointestinal tract within a 1-h timeframe^57,58^.

Following the diet exposure and supplementation periods, rats were switched to standard chow diet at PN 60, as with previous cohorts, and behavioral assessments were performed in adulthood (Fig. 5e). Rats that received the AGE-rich diet supplemented with *Lactococcus* in early life did not exhibit impairments in the novel object in context memory task and performed significantly better than the group that had received the AGE-rich diet and saline as a control treatment (Fig. 5g-h and Extended Data Fig. 5.2d-e). There were no differences in hippocampal-independent novel object recognition performance, or performance in either the zero maze or open field tests for anxiety-like behavior (Fig. 5i-j and Extended Data Fig. 5.2f-i), although there was a trending difference in distance travelled in the open field test (Extended Data Fig. 5.2j).

Given that the microbiome impacts circulating levels of short-chain fatty acids (SCFAs), branched-chain fatty acids (BCFAs), and lactate^59^, and that these metabolites can influence memory function^60^, we measured levels of SCFAs (acetate, propionate, butyrate), isobutyrate (a BFCA), and lactate in the serum, cecal, and fecal contents of rats fed the AGE-rich or control diet. Results revealed minimal changes in these metabolites as a function of diet (Extended Data Fig. 5.3a-o), with one significant group difference found for fecal isobutyrate, which was elevated in the AGE-rich vs. control group. These findings suggest that AGE-rich diet-associated reductions in *Lactococcus lactis* did not yield substantial changes in microbiome-associated metabolites, and by extension, that the memory deficits observed in rats fed the AGE-rich diet are unlikely due to altered levels of microbiome-associated metabolites.

Collectively, results indicate that heat treatment of an otherwise healthy rodent diet created a situation in which dietary AGEs negatively impacted neurocognitive function by two means: 1) by increasing levels of AGEs within the diet itself via the Maillard reaction, and, therefore, within the body; and 2) by diminishing the levels of *Lactococcus* (specifically *Lactococcus lactis* subspecies *lactis*), which have AGE-degrading enzymes^61^. Indeed, *Lactococcus lactis* not only degrades AGEs *in vitro*, but also *in vivo* in both humans and rats^61^. These findings imply a protective effect of this bacteria taxon on reducing the negative consequences of dietary AGE consumption, thus identifying a novel purpose for consuming increasingly popular probiotic supplements in humans.

## Discussion

Our findings reveal how the consumption of dietary advanced glycation end-products (AGEs) from an otherwise healthy diet during early life periods of development leads to long-lasting hippocampal-dependent memory impairments in rats. These neurocognitive effects are coupled with alterations in hippocampal glutamatergic neurotransmission and complement system activation, yet are observed independent of obesity, metabolic dysfunction, or nonspecific behavioral abnormalities. We further revealed that AGEs elevated in heat-treated foods act as competitive inhibitors of complement system receptors, which is likely driving the impaired synaptic plasticity observed from this dietary model. Furthermore, our results show a role for the gut microbiota – and even microorganisms within certain foods – in mediating these dietary AGE-induced effects.

While the levels of AGEs in ‘ultra-processed foods’ have not been comprehensively quantified, our diet contained approximately 2.3 mg CML per 100 g diet, which is within range of the amount of CML in candy bars (2.0 mg CML per 42-g portion of a Kit Kat bar and 2.9 mg CML per 51-g portion of a Milky Way bar)^62^. From a food processing and preparation perspective, findings from a recent crossover human study involving 2-week diet interventions in healthy adults indicated that serum AGE levels (CML, MG-H1, pyrraline) are modulated by altering cooking methods (e.g., steaming vs. grilling)^63^. While our results do not allow inferences to be made about ultra-processed food consumption specifically, it is possible that AGEs within some ultra-processed foods could play a role in their negative impacts on health. Indeed, recent findings based on the UK Biobank study showed that higher intake of the AGEs CML, CEL, and MG-H1 was associated with increased risk of dementia, independent of genetic risk^18^. These results are further corroborated by evidence linking elevated dietary AGEs with accelerated memory decline in elderly individuals^64,65^. While these studies examining AGEs in humans did not incorporate evaluation of systematic measures of food processing, another report exploring ultra-processed food consumption in a cohort from the UK Biobank study (participants 55+ years of age at baseline, followed for a median of 10 years) also found a relationship between higher intake of ultra-processed foods and increased risk of dementia^66^. Collectively, this evidence strongly suggests that dietary AGEs should be considered in future research investigating the neurocognitive impacts of processed food consumption.

Our experiments specifically focused on early life diet exposure, whereas previous research in rodents examining the impacts of dietary AGE consumption during adulthood on cognitive function has indicated similar deleterious effects^67,68^. Namely, in adult male and female C57BL/6 mice, Akhter et al.^67^ found that 17-month AGE-rich diet consumption (formulated as a modified diet containing 1000 ppm MG-BSA) resulted in memory deficits as evidenced through the Morris Water Maze test. Additionally, 7-month consumption of a high-AGE diet (generated through irradiation) accelerated memory impairments in the Morris Water Maze test in Tg2576 Alzheimer’s disease model male mice^68^. Given previous research indicating that the juvenile-adolescent stage (∼PN 26-60 in rats) is a particularly vulnerable period of cognitive development, especially for processes engaging the hippocampus^29,69–74^, we specifically targeted this early life period for the current experiments. Notably, we have previously shown that early-adolescence (PN 26-41) is an even smaller window of long-lasting hippocampal vulnerability to dietary insults in male rats^56^, which may contribute to how *Lactococcus* supplementation during this time in the present set of experiments was sufficient to prevent dietary AGE-induced memory deficits. Based on previous evidence in rodent models and correlations in aging humans^18,64,67,68^, we hypothesize that a longer duration of diet exposure may be required in order for behavioral phenotypes to develop when AGE-rich diet exposures take place later in life.

It is important to note that there are numerous AGE compounds beyond those that we quantified for the present work. CML, CEL, and MG-H1 are low molecular-weight AGEs that have been widely studied for years^75^, but there are hundreds more AGEs that can be found in heat-treated foods. As noted earlier, AGEs can also be bound vs. unbound to proteins^76–79^. It is thus possible that not all AGEs exert the same effects on the body. Furthermore, AGEs are not only generated in (and consumed through) foods, but they can also form endogenously within the body^80,81^. While our results have demonstrated that dietary AGEs can increase the levels of AGEs within circulation (in agreement with previous evidence^25,67,80^), whether high consumption of dietary AGEs also influences the formation of endogenous AGEs is poorly understood.

Our results establish evidence for a direct interaction between AGEs and the complement system, which has been implicated in synaptic development/plasticity^42,43^ and is likely a mechanism mediating the early life dietary AGE-induced memory deficits we observed. An AGE-rich diet has previously been linked with altered complement system signaling in the periphery, as well as impaired gut barrier integrity and kidney function in rats^22^. Our research builds upon these findings, as elevated AGEs in the periphery could also be acting on complement system receptors in the brain to disrupt their normal activation. An outstanding question is how precisely the dietary AGEs in our model transit to the brain to act on complement system receptors in the hippocampus. We reason that MG-H1, CML, and CEL are sufficiently small to pass through the blood-brain-barrier^46,47^, thus enabling them to act directly within the hippocampus as complement receptor antagonists. Consistent with this hypothesis, previous research revealed that mice exposed to isotopically-labelled dietary AGEs (CML) exhibited substantial AGE accumulation in the brain via RAGE-independent mechanisms^48^.

The current findings highlight the role that other dietary factors can have in influencing the impact of dietary AGEs on cognitive function. Indeed, previous evidence has indicated that *Lactococcus lactis* isolated from kimchi can degrade CML *in vitro* and prevent elevated levels of dietary CML within circulation in rats and humans^61^. In the context of the present work, we showed that heat treatment of the standard diet substantially decreased its inherent levels of *Lactococcus lactis*, which likely created an ideal environment for AGEs to accumulate and impart more severe effects. While certain types of thermal treatments are often used in food processing to eliminate or drastically reduce the levels of harmful microorganisms from a food safety standpoint, it is possible that these treatments also reduce the levels of beneficial bacteria (while also promoting the formation of AGEs, depending on the conditions). Beyond food-originating microorganisms, studies have also shown that polyphenols within or derived from foods (especially plant foods) can inhibit the formation of AGEs and possibly mitigate their harmful effects on the body^82–86^. Such dietary factors need to be taken into consideration when studying AGEs, and they also afford opportunities for the development of diet-based strategies to prevent and/or offset AGE-induced effects on the body.

Collectively, this work provides evidence for a functional connection between dietary AGE exposure during early life and impaired hippocampal-dependent memory function, while identifying the complement system as a potential mediator. Furthermore, our findings implicate the gut microbiota and food-originating microorganisms in heat-treated AGE-rich diet-induced memory impairments, signifying both the complexities and opportunities for understanding – and potentially offsetting – the effects of these food processing-induced compounds.

## Methods

### Subjects

Male Sprague Dawley rats (Envigo, Indianapolis, IN, USA) were used for all experiments. All rodent research experiments were performed in the animal vivarium at the University of Southern California. Accordingly, all animal procedures were approved by the University of Southern California Institutional Animal Care and Use Committee (IACUC) in accordance with the National Research Council Guide for the Care and Use of Laboratory Animals (protocol # 21096).

Rats arrived at the animal facility on postnatal day (PN) 25 and received their assigned diet starting on PN 26. They were housed under a reverse 12:12 h light:dark cycle, with lights turned off at 11:00 and on at 23:00, and with a temperature of 22-24 °C and humidity of 40-50%. All rats were singly housed in hanging wire cages throughout the experiments to facilitate food spillage collection for accurate measurement of individual food intake and prevention of coprophagy, with the exception of the rats for the electrophysiology experiments, which were housed in shoebox cages with elevated wire racks to prevent coprophagy. Rats were weighed three times per week between 8:30-10:30 (shortly prior to the onset of the dark cycle). The number of animals used for each cohort are provided in the general experimental overview below, and the specific number used in each analysis is included in Supplementary Table S1.

### Dietary model

To model a “processed” diet, we thermally treated an otherwise healthy pelleted, non-irradiated and non-autoclaved, AIN-93G Purified diet (TD.94045; Envigo, Indianapolis, IN, USA; 19% kcal from protein, 64% kcal from carbohydrate, 17% kcal from fat; 3.8 kcal/g) at 160 °C for 1 h in a forced-draft oven (ST-8; Across International, Livingston, NJ, USA). Pelleted AIN-93G Purified diet from the same batches was used as the control for all experiments. For the metabolism, microbiome, and behavior cohort, powdered AIN-93G Purified diet was used for an additional control group of rats (CTL-PWD) to determine if the pelletization process itself could impart changes on physiological or behavioral outcomes. Due to the change in hardness that occurred with heat treatment, both pelleted AIN-93G diets were marginally crushed with a dedicated hammer and accompanying setup to reduce their particle size and make them easier for the rats to consume. Diets were served in either small glass jars fastened to the sides of the cages or ceramic small animal pet food bowls. All animals received *ad libitum* access to water. If a given cohort had not been used for tissue collection, at PN 60 all groups of rats were switched to receive standard grain-based chow until the end of experimentation (LabDiet 5001; PMI Nutrition International, Brentwood, MO, USA; 28.5% kcal from protein, 13.5% kcal from fat, 58.0% kcal from carbohydrate; 3.36 kcal/g).

### General experimental overview per cohort of rats

Animal experiments were performed in a series of cohorts. In order to prevent initial differences in body weights across groups within a given cohort, rats were pseudo-randomly assigned to groups based on their body weights on PN 26. Experimental conditions differed per cohort, as detailed below. Experimental dietary exposures were implemented from PN 26-56 (for terminal tissue collection) or PN 26-60, which approximately represents the juvenile and adolescent periods in rats^87,88^. Terminal tissue collection occurred at either PN 56 or PN 102 depending on the cohort. For cohorts that underwent behavioral assays, all groups were switched to standard chow at PN 60 until the end of experimentation at PN 102 (at which point tissues were collected). Descriptions of the cohorts are as follows:

#### Tissue cohorts

Two cohorts of rats were dedicated solely to tissue collection. In each cohort, two groups of rats received either the heat-treated AGE-rich diet or control diet from PN 26-56, and terminal tissue collection occurred at PN 56 (N=16 total rats per cohort; n=8, CTL and n=8, AGE-rich). For one of these cohorts, tissues were collected when rats were in the fasted state (food removed at 9:00). For the other cohort, tissues were collected when rats were in the fed state. To facilitate this, rats were fasted for ∼16-17 h, fed a large meal in a staggered fashion, and euthanized 2 h post-meal. The euthanasia procedures used for these cohorts are described in the Supplementary Methods.

#### Metabolism, microbiome, and behavior cohort

One cohort of rats consisted of three groups that received either the heat-treated AGE-rich diet (n=12, AGE-rich), the pelleted control diet (n=12, CTL-PLT), or a powdered version of the control diet (n=12, CTL-PWD) from PN 26-60, at which point all groups were switched to standard chow (N=36 rats total). For assessments of metabolic outcomes, an intraperitoneal glucose tolerance test (IPGTT) was performed for this cohort at PN 59, and body composition was measured at PN 60 (prior to the diet switch that day). For characterization of the gut microbiome, fecal samples were collected at PN 56 and 90, and cecal samples were collected during tissue collection on PN 102. For behavioral assessments, novel object recognition (NOR) and novel location recognition (NLR) were performed from PN 76-79; novel object in context (NOIC) was performed from PN 80-84, zero maze was conducted on PN 87, and open field was conducted on PN 88-89. Procedures for these metabolic, microbiome, and behavioral assessments are described in the Supplementary Methods. All groups were sacrificed for tissue collection on PN 102 in the fasted state. Due to the lack of differences between groups receiving the non-heat-treated diets that were either pelleted (CTL-PLT) or powdered (CTL-PWD), these two groups have been combined as a shared control (CTL) in Fig. 1j-o and Fig. 2. Results for the distinct groups are shown in Extended Data Fig. 1d-k and Extended Data Fig. 2.

#### Electrophysiology cohort

To assess the effects of early life dietary AGE exposure on synaptic transmission, one cohort of rats was dedicated to electrophysiology experimentation. This cohort consisted of two groups that received either the heat-treated AGE-rich diet or control diet from PN 26-60+ (N=6 total rats; n=3, CTL and n=3, AGE-rich). Field potential recordings were performed with acutely collected hippocampus slices on separate days for each rat between PN 60-70 (i.e., rats were not switched to standard chow at PN 60) as described in the Supplementary Methods.

#### Alagebrium cohort

In order to determine whether the behavior effects observed were associated with AGEs themselves, one cohort of rats received the drug alagebrium (ALA; 1 mg/kg body weight/day given in drinking water; catalog # 28758, AstaTech, Bristol, PA, USA) or vehicle (drinking water alone) simultaneously with either the AGE-rich or control diet from PN 26-60. As such, there were four groups (N=48 rats total): AGE-rich with water (vehicle; n=12, AGE-rich), AGE-rich with ALA (alagebrium, drug; n=12, AGE-rich + ALA), CTL with water (n=12, CTL), and CTL with ALA (n=12, CTL + ALA). Fluid intake and body weights were measured daily – and fresh water and ALA solutions were provided for each rat – between 8:00-10:00 from PN 26-60 to facilitate the administration of ALA. At PN 60, all groups were switched to standard chow and no longer received ALA (as applicable), at which point fluid intake was no longer measured. NOR was performed from PN 75-78, NOIC was conducted from PN 80-84, and tissues were collected on PN 102 in the fasted state.

Regarding ALA administration, solutions were made fresh daily from PN 26-60 using reverse-osmosis water (the rodent drinking water for the vivarium), with ALA concentrations tailored based on the average amount of water that animals consumed the previous day. Concentrations of ALA ranged from 0.00067% to 0.0012%. The presence of ALA in drinking water at these concentrations did not affect the overall amount of fluid consumed (Extended Data Fig. 4b, Supplementary Table S1). Because ALA is light-sensitive, care was taken to ensure that solutions were not exposed to light by using opaque water bottles – and solutions were replaced daily.

Due to the labor-intensive nature of the ALA administration, the alagebrium cohort was split into two squads, with the two AGE-rich diet groups run as one squad and the two CTL diet groups run as the other squad. Rats in the AGE-rich diet squad had significantly higher body weights upon arrival (at PN 26) compared to rats in the CTL squad, and they exhibited different body weight growth trajectories (Extended Data Fig. 4a) and water intakes relative to their body weights (Extended Data Fig. 4b). These differences in body weight did not affect object exploration at critical stages of behavioral tasks or behavioral outcomes (Fig. 4b-e, Extended Data Fig. 4d-e).

#### *Lactococcus* cohorts (tissue and behavior)

To gain insights into the potential role of gut microbial taxa / dietary probiotics in AGE-induced effects on memory function, two cohort of rats received daily supplementation of *Lactococcus lactis* subspecies *lactis* (200 µL of 10^9^ CFU/mL; AGE-rich – LAC) or vehicle (0.9% saline; AGE-rich – SAL) via oral gavage between 12:00-13:00 (in the fed state, given that lights went off in the housing room at 11:00) from PN 26-40. This exposure timing was based on our previous research indicating that early adolescence (∼PN 26-40) is an especially vulnerable period for dietary influences on the hippocampus in male rats^56^. One cohort was used for tissue collection to verify whether the supplement resulted in increased levels of *Lactococcus lactis* subspecies *lactis* within the gastrointestinal tract. Behavioral assessments were performed in the other cohort. All rats in both cohorts received the heat-treated AGE-rich diet.

For the tissue cohort, rats received daily gavages from PN 26-40 (N=16 rats total; n=8, AGE-rich – LAC and n=8, AGE-rich – SAL). On the last day (PN 40), gavages were staggered so that each rat was euthanized ∼1 h thereafter, and all gavages were performed in the fed state (after lights out at 11:00). Contents from the stomach and cecum were collected in cryotubes, flash-frozen in dry ice, and transferred to a −80 °C freezer until further quantification of *Lactococcus lactis* subspecies *lactis* via qPCR as described in the Supplementary Methods.

For the behavior cohort, rats received daily gavages from PN 26-40, at which point gavages were ceased (N=14 rats total; n=6, AGE-rich – LAC and n=8, AGE-rich – SAL). Rats remained on either the heat-treated AGE-rich diet or control diet until PN 60 (as in other cohorts), at which point they were switched to standard chow until the end of the experiment. For this cohort, NOR was performed from PN 74-46, NOIC was conducted from PN 82-86, zero maze was performed on PN 87, and open field took place on PN 89. Tissues were collected on PN 102 in the fasted state.

### Additional methods

Methods for metabolic assessments (i.e., body weight, food intake, glucose tolerance, body composition), behavior assessments (i.e., novel object in context, novel location recognition, novel object recognition, zero maze, open field), euthanasia and tissue collection, quantification of AGEs in diets and biological samples, electrophysiology experimentation, AGE-complement system binding affinity assays, qPCR, gut microbiome analyses, gut metabolite analyses, and *Lactococcus* growth and quantification are all described in the Supplementary Methods file.

### Statistical analyses

Data are presented as means ± standard errors (SEM) for error bars where applicable in all figures. Each experimental group was analyzed with its respective control group(s) per experimental cohort. Statistical analyses were performed using Prism software (GraphPad, Inc., version 8.4.2, San Diego, CA, USA) or R (4.2.0, R Core Team 2022). Significance was considered at *p* < 0.05. Detailed descriptions of the specific statistical tests per figure panel can be found in Table S1. In all cases, model assumptions were checked (normality by Shapiro-Wilk test and visual inspection of a qq plot for residuals per analysis, equal variance/ homoscedasticity by an F test to compare variances and visual inspection of a residual plot per analysis). Data for quantification of MG-H1 and CML in serum, CML in urine, distance travelled during the open field test, performance in NOIC for the *Lactococcus* cohort (% shift from baseline), *Lactococcus lactis* abundance in cecal samples in the *Lactococcus* cohort, and percent time in the open arm from the zero maze test for the *Lactococcus* cohort (7 analyses) violated the assumption of homoscedasticity and thus Welch’s t-tests were employed for these analyses to account for unequal variances. Data for quantification of CEL in serum as well as CEL in urine violated the assumption of normality, so a two-sample Kolmogorov-Smirnov test was implemented for these analyses (2 analyses). Brown-Forsythe ANOVAs were performed for total object exploration during the final exposure of NLR and distance travelled during the open field test (2 analyses) due to unequal standard deviations between groups (CTL-PWD, CTL-PLT, AGE-rich). Non-parametric Wilcoxon rank sum tests were performed for comparing the abundance of individual taxa for the gut microbiome analyses (6 analyses). Group sample sizes were based on prior knowledge gained from extensive experience with rodent dietary studies involving behavior testing.

## Data availability

All data are available upon reasonable request from the corresponding author.

## Code availability

All codes are available upon reasonable request from the corresponding author.

## Acknowledgements

We thank all the Kanoski Lab undergraduate researchers for their support with experimentation. Figures 1a, 2a, 2b, 2e, 2h, 2k, 2n, 4a, and 5e were made with assistance from BioRender. This work was supported by the National Institute of Diabetes and Digestive and Kidney Diseases under grant DK123423 (awarded to SEK and AAF); the National Institute on Aging through a Postdoctoral Ruth L. Kirschstein National Research Service Award under grant F32AG077932 (awarded to AMRH); and the National Institute of Diabetes and Digestive and Kidney Diseases through Predoctoral Ruth L. Kirschstein National Research Service Awards under grants F31DK137484 (awarded to JJR) and F31DK138777 (awarded to MEK).

## Contributions

Conceptualization: A.M.R.H., S.E.K., M.T.C., L.A.S., B.E.H., C.G. Data curation: A.M.R.H., M.E.K., M.Sharma., S.S., E.L.G., D.N. Formal analysis: A.M.R.H., M.E.K., S.S., E.L.G., H.Z., J.C.D., R.J.C., D.R.S., D.N. Funding acquisition: S.E.K. Investigation: A.M.R.H., M.E.K., M.Sharma., A.E.K., S.S., E.L.G., H.Z., J.C.D., R.J.C., D.R.S., D.N., L.T., J.J.R., A.A., N.T., I.G., M.Shanmugam., Y.P. Methodology: A.M.R.H., M.Sharma., S.S., E.L.G., H.Z., J.C.D., R.J.C., D.R.S., D.N., V.M.M, L.T., K.B.Y., E.Y.H., A.A.F., T.M.W., S.C.S., C.G., B.E.H., M.T.C. Project administration and supervision: S.E.K., M.T.C., B.E.H., C.G. Validation: A.M.R.H., M.E.K., L.T., H.Z., J.C.D., R.J.C. Visualization: A.M.R.H., M.Sharma., H.Z., J.C.D., R.J.C., S.S., B.E.H., S.E.K. Writing: A.M.R.H., S.E.K. Writing – review and editing: all authors.

## Ethical statement

All experiments were approved by the Institutional Animal Care and Use Committee at the University of Southern California (protocol #21096) and performed in accordance with the National Research Council Guide for the Care and Use of Laboratory Animals, which is in compliance with the National Institutes of Health Guide for the Care and Use of Laboratory animals (NIH Publications No. 8023, revised 1978).

## Supplementary Methods

### Metabolic assessments

#### Body weight and food intake measurements

Unless otherwise indicated, body weights and food intakes – including spillage collected on cardboard placed under the hanging wire cage in which each rat was housed – were measured for all cohorts at least three times per week between 8:30-10:30 (shortly before the onset of the dark cycle). Food intake was calculated as the change in weight of the food receptacle and its contents across a given set of days, minus the weight of food spillage over the same time.

The number of total kilocalories each rat consumed for their respective diet was calculated by multiplying the amount of the diet consumed by its respective energy density. Because of the moisture loss that occurred with heat treatment, an energy density of 4.0 kcal/g was used for the heat-treated AIN-93G diet (determined by moisture loss experiments during which the weight of the pan containing a batch of pellets was weighed before and after baking [performed in triplicate]; data not shown; the difference indicated the amount of moisture loss). For the pelleted and powdered Purified AIN-93G diets an energy density of 3.8 kcal/g was used, and an energy density of 3.36 kcal/g was used for the standard chow diet.

#### Glucose tolerance

An intraperitoneal (IP) glucose tolerance test (IPGTT) was used to assess glucose tolerance on PN 59 in the cohort of rats that underwent metabolism, microbiome, and behavior assessments. Rats were food restricted for 22 h prior to the test, as done previously by our group^1–4^. Blood was sampled from the tip of the tail for baseline blood glucose readings (performed in the fasted state) and all subsequent blood glucose readings. A glucometer was used to measure glucose levels (One Touch Ultra2, LifeScan, Inc., Milpitas, CA, USA). After collecting the baseline reading, each rat received an IP injection of a 50% glucose solution (1 g/kg body weight; Dextrose anhydrous, VWR Chemicals, LLC, Solon, OH, USA).

Aside from the baseline reading, blood glucose measurements were subsequently collected at 30, 60, 90, and 120 min following injection. Area under the curve for each rat was calculated using the trapezoidal rule with 0 mg/dL set as the mathematical “baseline”.

#### Body composition

Nuclear magnetic resonance (NMR) was used to determine body composition on PN 60 in the cohort of rats that underwent metabolism, microbiome, and behavior assessments according to previous procedures^1,4–6^. Rats were lightly food restricted for 1 h, weighed, and scanned using a Bruker NMR Minispec LF 90II (Bruker Daltonics, Inc., Billerica, MA, USA) interfaced to a computer equipped with Minispec Plus 4.1.5 software. Percent fat mass, percent lean mass, and fat-to-lean ratio were calculated as [fat mass (g)/body weight (g)] × 100, [lean mass (g)/body weight (g)] × 100, and [fat mass (g)/lean mass (g)], respectively.

### Behavior assessments

#### Novel object in context (NOIC)

Hippocampal-dependent contextual episodic memory function was assessed using NOIC^7,8^. The 5-day procedure was adapted from previous methods^8^, conducted in accordance with previous studies by our group^1,2,4,5,9–12^.

Each day consisted of one 5-min session per rat, with cleaning of the apparatus and objects using 10% ethanol between rats. The two initial days were habituation to the two contexts used, such that rats were exposed to one context per day in a counterbalanced order per diet group. Accordingly, rats were placed in Context 1, a semitransparent box (41.9 cm L × 41.9 cm W × 38.1 cm H) with yellow stripes, or Context 2, a black opaque box (38.1 cm L × 63.5 cm W × 35.6 cm H) in a counterbalanced order. Contexts were presented in distinct rooms, both with similar dim ambient lighting yet distinct extra-box contextual features. On the subsequent day after the two habituation days, each animal was placed in Context 1 containing single copies of Object A and Object B situated on diagonal equidistant markings with sufficient space for the rat to circle the objects (NOIC Day 1; Fig. 2b). Objects were an assortment of hard plastic containers and tin canisters (with covers). Objects were distinct from what animals were exposed to for NOR and NLR, which are described below. The sides where the objects were placed were counterbalanced per rat by diet group. On the following day (NOIC Day 2), rats were placed in Context 2 with duplicate copies of Object A. The next day was the test day (NOIC Day 3), during which rats were placed again in Context 2, but with single copies of Object A and Object B; Object B was therefore not a novel object, but its placement in Context 2 was novel to the rat (‘novel object in context’). Each time the rats were situated in the contexts, care was taken so that they were consistently placed with their head facing away from both of the objects. On NOIC Days 1 and 3, object exploration, defined as the rat sniffing or touching the object with the nose or forepaws, was quantified by hand-scoring of videos by an experimenter blinded to animal group assignments. The discrimination index for Object B was calculated for NOIC days 1 and 3 as follows: [(time spent exploring Object B, i.e., the ‘novel object in context’ in Context 2) / (time spent exploring Object A + time spent exploring Object B)]. Data were then expressed as a percent shift from baseline as: [day 3 discrimination index – day 1 discrimination index] × 100. Rats with proper hippocampus function will preferentially explore the ‘novel object in context’ on NOIC day 3, while an impairment in hippocampus-dependent memory function will impede such preferential exploration^7,8^. Total object exploration time on NOIC Days 1 and 3 were also analyzed.

#### Novel location recognition (NLR)

NLR was performed to assess hippocampal-dependent spatial recognition memory^13,14^, as performed previously by our group^4,6,12^. A grey opaque box (38.1 cm L × 56.5 cm W × 31.8 cm H), placed on a table, was used as the NLR apparatus in a room in which two adjacent desk lamps were pointed toward the tabletop to provide dim lighting. Rats were habituated to the empty apparatus and conditions for 10 min 1 or 2 days prior to testing. Testing consisted of a familiarization phase (‘initial exposure’) during which rats were placed in the center of the apparatus (facing a neutral wall to prevent biasing them toward either object) with two identical objects placed in two corners of the apparatus.

Rats were allowed to explore for 5 min during this initial exposure period. Objects consisted of either two identical empty glass salt-shakers (that never contained salt) or two identical empty soap dispensers (that were never used with soap; first NLR time point), counterbalanced across rats per group. Following the initial exposure period, rats were removed from the apparatus and placed in their home cage for 5 min, during which time the apparatus and objects were cleaned with 10% ethanol solution and one of the objects was moved to a different corner location in the apparatus (i.e., the object was moved but not replaced). Rats were then placed in the center of the apparatus again and allowed to explore for 3 min, which constituted the ‘final exposure’ (or test exposure). The types of objects used and the novel location placements were counterbalanced by group. The time each rat spent exploring the objects was quantified by hand-scoring of video recordings by an experimenter blinded to animal group assignments.

Object exploration was defined as the rat sniffing or touching the object with its nose or forepaws. An ‘exploration index’ was calculated for each rat as [amount of time spent exploring the object moved to a novel location (s)/total object exploration time (s)] during the test exposure period. Total object exploration time during the initial and final (test) exposure periods were also analyzed.

#### Novel object recognition (NOR)

Evaluation of object-novelty recognition was performed using the NOR task. We performed this task in such a manner that it engaged the perirhinal cortex and not the hippocampus^15^. A grey opaque box (38.1 cm L × 56.5 cm W × 31.8 cm H; the same box that was used for NLR), placed on a table in a dimly lit room in which two adjacent desk lamps were pointed toward the floor, was used as the NOR apparatus. Previous procedures were followed^4,6,11,16^, modified from Beilharz et al.^17^ Rats were habituated to the empty apparatus and conditions for 10 min 1 or 2 days prior to testing. The test consisted of a 5-min familiarization phase (‘initial exposure’) during which rats were placed in the center of the apparatus (facing a neutral wall to prevent biasing them toward either object) with two identical objects and allowed to explore. The objects used were either two identical cans or two identical stem-less wine glasses. Rats were then removed from the apparatus and placed in their home cage for 5 min. During this period, the apparatus and objects were cleaned with 10% ethanol solution and one of the objects was replaced with a different one (i.e., either the can or glass – whichever the animal had not previously been exposed to – i.e., the ‘novel object’). Rats were then placed in the center of the apparatus again and allowed to explore for 3 min for the final (or test) exposure. The novel object and side on which the novel object was placed were counterbalanced per treatment group. The time each rat spent exploring the objects was quantified by hand-scoring of video recordings by an experimenter blinded to animal group assignments. Object exploration was defined as the rat sniffing or touching the object with the nose or forepaws. An ‘exploration index’ was calculated for each rat as [amount of time spent exploring the novel object (s)/total object exploration time (s)] during the test exposure period. Total object exploration time during the initial and final (test) exposure periods were also analyzed.

#### Zero maze

The zero maze procedure was used to examine anxiety-like behavior according to established procedures^1,2,4,9,11^. The zero maze apparatus consisted of an elevated circular track (11.4 cm wide track, 73.7 cm height from track to the ground, 92.7 cm exterior diameter) divided into four segments of equal lengths: two sections with 3-cm high curbs (open), and two sections with 17.5 cm high walls (closed). Ambient lighting was used during testing.

Rats were individually placed onto the maze on an open section of the track and allowed to roam for 5 min, during which time they could freely ambulate through the different segments of the track. The apparatus was cleaned with 10% ethanol between rats. The time each rat spent in the open segments of the track (defined as the center of the rat in an open arm) as well as the number of entries into the open segments of the track were measured via video recording using ANYmaze activity tracking software (Stoelting Co., Wood Dale, IL, USA).

#### Open field

Anxiety-like behavior and locomotor activity were assessed using the open field test according to previous procedures^3,4,9–11,18^. A gray arena (53.5 cm L × 54.6 cm W × 36.8 H) with a designated center zone within the arena (19 cm × 17.5 cm) was used as an apparatus. The center zone was maintained under diffused lighting (∼44 lux) compared to the corners and edges (∼30 lux). For testing, rats were placed in the center of the apparatus and allowed to freely explore for 10 min. The apparatus was cleaned with 10% ethanol between rats. Video recording with ANYmaze activity tracking software (Stoelting Co., Wood Dale, IL, USA) was used to quantify time spent in the center zone (a measurement of anxiety-like behavior) and distance travelled in the apparatus during the test (an indicator of locomotor activity).

### Euthanasia and tissue collection

#### Euthanasia

With the exception of the electrophysiology cohort described below, all other cohorts were euthanized as follows: After deep anesthesia using an intramuscular injection of a cocktail of ketamine (90.1 mg/kg body weight), xylazine (2.8 mg/kg body weight), and acepromazine (0.72 mg/kg body weight), animals were rapidly decapitated.

#### Brain

Brains were quickly extracted, flash-frozen in a beaker containing −30 °C isopentane surrounded by dry ice for 3 min, transferred to pre-labeled bags, and kept on dry ice until being placed in a −80 °C freezer for longer-term storage until further processing and analysis. Tissue punches of the dorsal hippocampus (atlas levels 28-32; one 2 mm diameter punch in the CA1 / dentate gyrus, one 1 mm diameter punch in the CA3) for qPCR analyses were later collected using a Leica CM cryostat (Wetzlar, Germany) and anatomical landmarks were based on the Swanson rat brain atlas^19^. Punches were stored at −80 °C until being used for qPCR analyses as described below.

#### Serum

For serum collection, trunk blood was collected into microcentrifuge tubes after decapitation and either immediately spun in a centrifuge at 4 °C at 2000 × g for 1-4 min or allowed to clot on wet ice for 15 min and then spun in a centrifuge at 4 °C at 2000 × g for 10 min. Supernatants were collected after centrifugation and immediately flash-frozen on dry ice until being transferred to a −80 °C freezer until further processing and analysis.

#### Brown adipose tissue

Brown adipose tissue (BAT) was collected from the tissue cohort for which euthanasia was performed in the fasted state. Accordingly, interscapular BAT was dissected, weighed, and flash-frozen in a beaker filled with isopentane surrounded by dry ice for 3 min. Percentage of BAT per rat was calculated as [g BAT/g body weight] × 100.

#### Duodenal and jejunal scrapes

Intestinal scrapes for quantification of tight junction proteins in the duodenum and jejunum were collected from the tissue cohort euthanized in the fed state according to previous procedures^2,20^. Briefly, an abdominal incision was made to expose the intestine. The small intestine was excised. A 1-cm segment distal from the pyloric sphincter as well as a 1-cm segment proximal from the cecum were removed and discarded. The remaining small intestine was folded in half, and the half-way point was identified as the jejunum. An incision was made at the half-way point. Two 1-cm segments distal from the identified half-way point were cut; the first was discarded and the second (more distal segment) was kept as the jejunum sample. For the duodenum sample, the next-most-distal 1-cm segment from the pyloric sphincter was cut. Both segments from the duodenum and jejunum were flushed with saline. Intestinal villi from each segment were then collected by scraping with a metal spatula, storing in RNAlater overnight, transferring to a Dnase/Rnase-free 2 mL tube, and then storing in a −80 °C freezer until further processing and analysis via qPCR.

#### Stomach content

For the *Lactococcus* tissue cohort, contents from the stomach were collected in cryotubes, flash-frozen in dry ice, and transferred to a −80 °C freezer until further quantification of *Lactococcus lactis* subspecies *lactis* via qPCR.

#### Cecal content

Cecal content was collected as described in the Microbiome section below.

### Quantitation of protein-bound advanced glycation end-products (AGEs) in diets

#### Diet sample preparation

Three pellets each of baked and unbaked AIN-93G rodent chow were individually processed for determination of protein-bound AGEs (carboxymethyllysine [CML], carboxyethyllysine (CEL), methylglyoxal-derived hydroimidazolone [MG-H1]) by LC/MS. One gram of powdered pellet was extracted overnight with 10 mL chloroform/methanol 3:1 and the residue was dried under a stream of nitrogen. 50 mg of the defatted pellets were subjected to total proteolytic digestion, i.e. sequential treatment with each trypsin (0.5% w/v), soybean trypsin inhibitor, collagenase, peptidase, pronase E, and aminopeptidase in HEPES buffer initially containing 1M CaCl_2_ at the trypsin step. The final solution (about 1.1 mL) was passed over SpinEx filters and 10kDa ultra centrifugal filters. A 1:10 dilution was spiked with isotopically labeled internal AGE standards of d2-CML, d3-CEL, d3-MG-H1, d3-methionine sulfoxide, d2-tyrosine, d6-fructose-lysine and 2% final heptafluorobutyric acid (HFBA). 20 µL were used for AGE analysis by mass spectrometry. CML.

#### High Pressure Liquid Chromatography (HPLC) - Mass Spectrometry (MS)

The LC system consisted of an Agilent Series 1100 HPLC equipped with a quaternary pump (G1311A), autosampler (G1313A), degasser (G1322A) and a temperature controlled column compartment (G1322A). The mass spectrometer was a Thermo TSQ Quantum Access (Thermo Electron Corp.) interfaced to the HPLC using a Dell Vostro 220 computer operated by Windows XP and loaded with Thermo LC-MS software including Xcalibur v2.07, TSQ v2.2 and LC Devices v2.0.2.

#### Quantitative Analyses of AGEs by HPLC-MS/MS

Markers of glycation were quantitated by liquid chromatography mass spectrometry using the HPLC-MS system described above. Analytical separations were made at flow rates 0.2 mL/min on a Discovery HS C18, 150 × 2.1 mm, 3 µm column using a 2 cm × 2.1 mm guard column with same resin (Millipore Sigma). The column temperature compartment was 35 °C. The mobile phases were: (A) water + 0.1% heptafluorobutyric acid (HFBA, Product 25003, Thermo); (B) 90% acetonitrile /10% water (Burdick & Jackson, Honeywell, Muskegon, MI) with 0.1% HFBA same. Samples consisting of 10 KD filtrates of food digests ∼5 µg protein were injected onto the column every 40 min with the following gradient program: 0-5 min, 100% A; 5.1-15 min, 1% - 25% B (linear); 15-15.5 min, 25% B; 15.5-17 min, 25% - 100% B (linear); 17-27 min, 100% B; 27.1-40 min, 100% A. The TSQ Quantum Instrument method was composed of 4 segments which were programmed by Xcalibur software according to the retention times of measured AGEs and as previously described by Monnier et al.^21^

### Quantification of AGEs in biological samples

AGEs were quantified in serum and urine. Serum samples were collected from the tissue cohort that underwent euthanasia in the fed state as described above. Urine samples were collected between 12:00-15:00 on PN 53 in the same cohort. Rats were kept in a neutral clean cage (without bedding) until urination. Urine was quickly pipetted as aliquots into 2 mL tubes and embedded in dry ice until being transferred to an ultra-low freezer at −80 °C. Serum and urine samples were transported from the University of Southern California (Los Angeles, CA, USA) to City of Hope (Duarte, CA, USA) on dry ice, where they were stored at −80 °C until preparation for AGEs quantification.

Serum and urine samples were diluted with formic acid and centrifuged for 10 min at 4 °C and 17,000 × g, at which point the supernatant was transferred to new tubes. Samples were then purified using Oasis MCX SPE columns, evaporated in a speed vac overnight with no heat, and resuspended in LCMS-grade H_2_O. Samples were then analyzed on a Thermo Scientific TSQ Altis Plus triple quadrupole mass spectrometer in positive ion mode coupled with a Thermo Scientific Vanquish Flex UHPLC system. Samples were stored in a 4 °C autosampler during the analysis and run using three distinct methods for creatinine, MG-amino acids, or MG-nucleosides. All mass spectrometry data was quantified using the Skyline software version 24.1.

Creatinine, MG-H1, and CML were measured using an Acquity Premier BEH Amide 1.7 µm VanGuard FIT 2.1 x 100 mm column using mobile phase A: 0.1% formic acid in LC-MS grade water; and mobile phase B: 0.1% formic acid in LC-MS grade water. Separation was carried out with the following gradient: 0-1 min, 90% B; 1-4 min, 90-30% B; 4-7 min, 30-90% B; 7-11 min, 90% B. Ion analysis was performed with H-ESI positive ion mode with ion transfer tube temperature of 325 °C, vaporizer temp 350 °C, 2000 V spray voltage, 50 sheath gas, 20 aux gas, 1 sweep gas, 1.5 mTorr collision gas pressure, Q1 FWHM of 0.7, and Q3 FWHM of 1.2. Analytes were measured using the following fragmentation protocol: creatinine, 114.000 > 86.000 *m/z*, collision energy 17 V; d3-creatinine, 117.000 > 89.000 *m/z*, collision energy 17.0 V V; CML, 205.080 > 130.050 *m/z*, collision energy 13.020 V; d4-CML, 209.120 > 134.050 *m/z*, collision energy 13.520 V; MG-H1, 229.110 > 166.050 *m/z*, collision energy 16.530 V; d3-MG-H1, 232.110 > 169.050 *m/z*, collision energy 16.530 V.

CEL was separated using an Atlantis Premier BEH Z-HILIC 1.7 µm VanGuard FIT 2.1 x 100 mm column using mobile phase A: 0.1% formic acid in LC-MS grade water; and mobile phase B: 0.1% formic acid in LC-MS grade water. Separation was carried out with the following gradient: 0-1 min, 90% B; 1-7 min, 90-30% B; 7-7.1 min, 30-15% B; 7.1-10.0 min, 15% B; 10.0-10.1 min, 15-90% B; 10.1-15.5 min, 90% B. Ion analysis was performed with identical instrument parameters as mentioned above. Fragmentation was performed using the following protocol: CEL, 219.090 > 130.050 *m/z*, collision energy 13.520 V; d4-CEL, 223.120 > 134.050 *m/z*, collision energy 13.810 V.

CEdG and CEG were separated using an Acquity Premier HSS T3 1.8 µm column using mobile phase A: 0.1% formic acid in LC-MS grade water; and mobile phase B: 0.1% formic acid in LC-MS grade water. Separation was carried out with the following gradient: 0-0.8 min, 0% B, 0.4 mL/min; 0.8-4.5 min, 0-3% B; 4.5-8.2 min, 3-10% B; 8.2-8.5 min; 10-97% B; 8.5-9 min, 97% B; 9-9.5 min; 97-0% B, 0.4-0.5 mL/min; 9.5-14.0 min, 0% B, 0.5 mL/min. Ion analysis was performed with instrument parameters of: 3500 V spray voltage, 300 C ion transfer tube temperature, 350 C vaporizer temp, 50 sheath gas, 10 aux gas, 1 sweep gas, 1.5 mTorr collision gas pressure, 0.4 Q1 FWHM, and 1.2 Q3 FWHM. Analytes were measured using the following fragmentation protocol: CEdG, 340.08 > 224.00 *m/z*, collision energy 10.675 V; ^13^C_10_-^15^N_5_-CEdG, 355.140 > 234.080 m/z; collision energy 10.080 V; CEG 356.100 > 224.083 *m/z*, collision energy 12.230 V; ^15^N_5_-CEG 361.110 > 229.060 *m/z*, collision energy 12.230 V.

### Electrophysiology experiments

#### Brain extraction and slicing

Rats from the electrophysiology cohort were anesthetized with isoflurane and absence of reflex was checked by toe pinch. The animal was decapitated using a small guillotine and the whole brain was extracted as previously described^22^. The brain was quickly placed into an ice-cold oxygenated NMDG-artificial cerebrospinal fluid (aCSF) solution containing (in mM): 135 NMDG, 2.5 KCl, 1.2 NaH_2_PO_4_, 20 HEPES, 10 glucose, 2.5 MgSO4, 0.5 CaCl_2_, 5 sodium ascorbate, 3 sodium pyruvate and cooled for 1 minute^23^.

Afterwards, the brain was placed into a petri dish partially submerged in NMDG-aCSF solution and cut in half to separate the two hemispheres. The hippocampus from both hemispheres was gently removed using a curved spatula, mounted on an agarose block, and placed in a vibratome filled with oxygenated NMDG-aCSF solution. 350 uM sections were cut and slices were placed into a 32C oxygenated recovery chamber filled with aCSF solution containing (in mM): 125 NaCl, 26 NaHCO_3_, 2.5 KCl, 1.2 NaH_2_PO_4_, 10 glucose, 1.3 MgSO_4_, 2.5 CaCl_2_, 5 sodium ascorbate, 3 sodium pyruvate for 1.5 h. Note: NMDG-aCSF and aCSF solutions were calibrated to give osmolarity 305 - 315 mOsm/Kg and pH 7.3 - 7.4.

#### Field potential recordings

Acute slices were placed in recording chamber and maintained in artificial cerebrospinal fluid (aCSF) solution containing (in mM): 125 NaCl, 26 NaHCO_3_, 2.5 KCl, 1.2 NaH_2_PO_4_, 10 glucose, 1.3 MgSO_4_, and 2.5 CaCl_2_ for 10 min before proceeding with field recordings. The stimulating glass electrode was filled with aCSF and placed in the stratum radiatum region of area CA1 to orthodromically stimulate Schaffer collateral/commissural fibers^24^. The recording glass electrode was filled with aCSF and placed 1-2 mm from the stimulating electrode to record field excitatory postsynaptic potentials (fEPSPs). Stimulations consisted of current pulses lasting 100 μs applied at 0.05 Hz at varying stimulation intensities (μA) to achieve fiber volley amplitudes of 0.1 mV, 0.2 mV, 0.3 mV, 0.4 mV, and 0.5 mV. The magnitude of fEPSPs was recorded for each corresponding fiber volley measurement. Only slices exhibiting stable field potentials at each fiber volley amplitude were used for analysis, otherwise they were discarded.

### AGE-complement functional analysis – pERK 1/2 Signaling Assay

#### Cell culture

Chinese hamster ovary cells stably expressing C5aR1 (CHO-C5aR1) were maintained in Ham’s F12 media supplemented with 10% FCS, 100 U/mL penicillin, 100 µg/mL streptomycin and 400 µg/mL G418 (Invivogen, San Diego, USA). The cell line was maintained in T175 flasks (37OC, 5% CO2) and subcultured at 90% confluency using TrypLE Express (Thermo Fisher Scientific, Melbourne, Australia)

#### pERK 1/2 Signaling Assay

Ligand-induced phospho-ERK 1/2 signaling was assessed using the Alpha LISA SureFire Ultra pERK 1/2 (Thr202/Tyr204) assay kit (PerkinElmer, Melbourne, Australia). CHO-C5aR1 cells were seeded (50,000 cells/well) onto 96-well tissue culture-treated plates, incubated for 24 hours and subsequently serum starved overnight.

Ligands were initially dissolved in 10% DMSO in H_2_O, with further dilutions prepared in serum-free medium (SFM). To screen for agonist activity, cells were stimulated with respective ligands for 10 minutes and then immediately lysed using AlphaLISA lysis buffer on a microplate shaker (10 min, 450 rpm). To screen for antagonist activity, cells were pretreated with ligands MG-H1, CML, or CEL (Iris Biotech) for 30 minutes in SFM before stimulation with 0.3 nM C5a (CHO-C5aR1) for 10 minutes, after which they were lysed as described previously. Cell lysate (5 µL/well) was added to a 384-well ProxiPlate (PerkinElmer, Melbourne, Australia) followed by the donor and acceptor reaction mixes (2.5 µL/well each). Following a 2-hour incubation in the dark, the plate was read on a CLARIOstar Plus microplate reader following standard AlphaLISA settings.

#### Data collection and analysis

Experiments were conducted in triplicate and conducted on at least three different days. Data were analyzed using GraphPad software (Prism 10.1.2) and expressed as mean ± standard error of mean (SEM). For each replicate, data was normalized prior to being combined. Logarithmic concentration-response curves were plotted using combined data and analyzed to calculate the potencies of each ligand.

### AGE-complement functional analysis – RG-BRET assay

An Receptor-G protein-Bioluminescence Resonance Energy Transfer (RG-BRET) cell-based signaling assay with HEK 293 cells was used to determine functional reactivity between AGEs and the complement system receptor C5aR1 according to previous methods^33^. Briefly, transient transfections of sequence for wild-type and mutant constructs of full-length human C5aR1 in the pcDNA 3.1(-) vector were performed in mammalian HEK293F cells. Receptors were expressed under the Cytomegalovirus (CMV) promoter with an N-terminal HA signal peptide and a FLAG tag. Rluc8 was fused to the C terminus of the wild-type C5aR1 construct with a four-amino acid linker (SRGG). HEK293F cells were maintained in FreeStyle 293 Expression Medium (Gibco). All cell lines were maintained at 37 °C with 5% CO_2_ and shaking at 110 rpm. C5aR1-Rluc8 and Gαi/Gβ1/Gγ2-GFP2 plasmids were transfected at a 1:1 ratio. The C5aR1-selective antagonist avacopan (1 µM) was used to block interactions and confirm specific binding.

The cells were transferred to 96-well plates containing a distinct AGE ligand (1 µM of either MG-H1, CML, or CEL [all from Iris Biotech]) in ligand buffer (20 mM HEPES, pH 7.5, 1× HBSS, 0.1% BSA). BRET measurements were performed using a Synergy H4 Multi-Mode microplate reader (Agilent Technologies).

### qPCR

Dorsal hippocampus punches and intestinal tissues were collected and stored as described above. mRNA expression of complement component 3 (C3), complement component 5 (C5), C3a receptor (C3aR), C5a receptor 1 (C5aR1), the receptor for AGEs (RAGE), nuclear factor-ĸB (NFĸB), and postsynaptic density-95 (PSD-95) protein was quantified in the dorsal hippocampus from the tissue cohort for which euthanasia was done in the fasted state. mRNA expression of occludin, claudin-1, claudin-5, and mucin 2 was quantified in duodenum and jejunum scrape samples from the tissue cohort for which euthanasia was done in the fed state. A reverse transcriptase quantitative polymerase chain reaction (RT-qPCR) was performed as previously described^5^ and done by our group^3^. Briefly, the total RNA was extracted from hippocampus punches using the RNeasy Lipid Tissue Mini Kit (Cat. no. 1023539, Qiagen, Hilden, Germany) and from intestinal tissues using the RNeasy Mini Kit (Cat. no. 74104, Qiagen), both according to the manufacturer’s instructions. Total RNA concentration per extracted sample was measured using a NanoDrop Spectrophotometer (ND-ONE-W, ThermoFisher Scientific, Waltham, MA, USA). RNA (>250 ng) was reverse transcribed to cDNA using a QuantiTect Reverse Transcription Kit (Cat. no. 205314, Qiagen) according to the manufacturer’s instructions. The following Applied Biosystems probes for use with rats were used: *C3* (Rn00566466_m1), *C5* (Rn01436156_m1), *C3aR* (Rn00583199_m1), *C5aR1* (Rn02134203_s1), *RAGE* (Rn01525753_g1), *NfĸB* (Rn01502266_m1), *PSD-95* (Rn00571479_m1), *occludin* (Rn00580064_m1), *claudin-1* (Rn00581740_m1), *claudin-5* (Rn01753146_s1), and *mucin 2* (Rn01498206_m1). qPCR was performed using Applied Biosystems TaqMan Fast Advanced Master Mix (Cat. no. 444557, ThermoFisher Scientific) and an Applied Biosystems QuantStudio 5 Real-Time PCR System (ThermoFisher Scientific). All reactions were conducted in triplicate, with triplicate cycle threshold (Ct) values averaged and normalized to GAPDH expression (Rn99999916_s1) for hippocampus tissues and to Hprt1 expression (Rn01527840_m1) for intestinal samples. The comparative 2^-ΔΔCt^ method was utilized to quantify relative expression levels between groups for the genes of interest.

### Gut microbiota analyses

#### Collection of fecal samples

Fecal samples were collected on PN 56 and 90 from the metabolic, microbiome, and behavior cohort. Rats were placed into a sterile cage (without bedding) and mildly restrained until defecation. Two fecal samples per rat per time point were collected. Each was weighed immediately upon defecation under sterile conditions and placed into a distinct Dnase/Rnase-free 2 mL cryogenic vial and immediately embedded in dry ice (one sample per vial). All samples were then stored in a −80 °C freezer until further processing and analysis. Between every rat, all cages and sample collection materials were thoroughly cleaned with 70% ethanol. All fecal collections were performed between 12:00 and 15:00 (during the dark cycle).

#### Collection of cecal samples

Collecting cecal content is a terminal procedure, and thus it was performed on the terminal tissue collection days for the metabolic, microbiome, and behavioral cohort as well as for the *Lactococcus* tissue cohort (performed at PN 102 and PN 40 for the two cohorts, respectively). Following euthanasia procedures (described above), an abdominal incision was made to expose the intestines. The cecum was identified and extracted. A small incision was made therein. Cecal content was then collected directly into a pre-tared Dnase/Rnase-free 2 mL cryogenic vial, weighed, and immediately embedded into dry ice. All samples were then stored in a −80 °C freezer until further analysis. Two vials of cecal content were collected per rat. Between every rat, all cecal content collection materials were thoroughly cleaned with 70% ethanol. Cecal content collections were performed between 11:00 and 17:00 (during the dark cycle).

#### Analyses

Demultiplexed reads of the 16S rRNA amplicon sequencing were analyzed with DADA2 and QIIME2 following the developers’ instructions. Denoising, dereplication, ASV inference, and chimera removal were performed with DADA2^25,26^. Taxonomy was assigned with QIIME2 feature classifier using the SILVA database (release 138). The taxonomic abundance was normalized to correct for different sequencing depth as previously described^27^. Statistical analyses were performed with R (4.3.1). Beta-diversity was computed using Bray-Curtis dissimilarity and visualized with principal coordinates analysis (PCoA) using R function ‘capscale’ in the package ‘vegan’. The differences in microbial community profiles between groups were analyzed with PERMANOVA test using function ‘adonis2’ in the same package with 999 permutations. Shannon index was used for analyzing alpha-diversity. Differential abundance of individual taxa between groups were analyzed using non-parametric Wilcoxon rank sum test and correlation tests were performed using Spearman’s correlation. Taxa with presence in less than 25% samples were excluded to avoid spurious associations. P-values were corrected for multiple hypothesis testing using the Benjamini-Hochberg method.

### Gut metabolite analyses

Acetate, propionate, and butyrate (short-chain fatty acids), isobutyrate (a branched-chain fatty acid), and lactate were quantified in serum, cecal, and fecal samples from the tissue cohort for which euthanasia was performed when rats were in the fasted state on PN 56.

Quantifications were performed using LC-MS through the UCLA Analytical Phytochemical Core (Los Angeles, CA, USA) according to previous procedures^28^, modified from established protocols^29,30^.

### Lactococcus experiments

#### Selection and culturing of Lactococcus lactis subspecies lactis

Through 16s sequencing, the sequencing variant that drove the *Lactococcus* genus changes at PN 56 was determined to be: TACGTAGGTCCCGAGCGTTGTCCGGATTTATTGGGCGTAAAGCGAGCGCAGGTGGTTTATTAAGTCTGGTGTAAAAGGCAGTGGCTCAACCATTGTATGCATTGGAAACTGGTAGACTTGAGTGCAGGAG. *Lactococcus lactis* subspecies *lactis* was determined to be a close match via a microbial BLAST search (score of 241, 10 hits; BLAST search tool [Basic Local Alignment Search Tool, http://blast.ncbi.nlm.nih.gov/Blast.cgi]). Thus, we focused our efforts on examining the potentially mediating effects of supplementation with *Lactococcus lactis* subspecies *lactis* on early life AGE-rich diet-related behavioral outcomes.

#### Growth and use of *Lactococcus lactis* subspecies *lactis*

*Lactococcus lactis* subspecies *lactis* strain NCTC 6681 (Lister, Schleifer et al., cat. no. 19435) was purchased from the American Type Culture Collection (ATCC, Manassas, VA, USA), rehydrated, and inoculated according to manufacturer’s instructions. Briefly, the pellet was rehydrated with ATCC Medium #44 (brain heart infusion broth), aseptically transferred to a sterile tube, and mixed, at which point several drops of the resulting suspension were pipetted onto a #44 agar plate and incubated for 24 h at 37 °C until colonies formed. Bacterial stock from these colonies was frozen in 50% glycerol at −80 °C until future use. We determined that the early stationary phase for this strain of *Lactococcus lactis* subspecies *lactis* was between 8-12 h based on a 24-h growth curve experiment in which colony forming units (CFUs) were determined for samples collected every hour after inoculation of brain heart infusion broth via serially diluted plate counts (data not shown). Thus, *Lactococcus lactis* subspecies *lactis* colonies were grown on a 12:12 h schedule, by initially placing one colony from the bacterial stock in 5 mL of brain heart infusion broth for 12 h as the initial inoculation culture, and then transferring 0.4 mL of this inoculation culture to 39.6 mL of broth and allowing it to grow for 12 h. Cultures were then aliquoted and centrifuged at 4.0 × g for 5 min, supernatants were removed and discarded, and the resulting pellets were resuspended in saline (0.9%) to be given as supplements to the rats in the *Lactococcus* cohort. Consistency in obtaining 10^9^ CFU/mL on three separate days was verified by plate counts (Extended Data Fig. 5.2b). *Lactococcus lactis* subspecies *lactis* suspensions (200 µL of 10^9^ CFU/mL) and saline (vehicle; 200 µL) gavages were performed in experimental (AGE-rich - LAC) and control (AGE-rich - SAL) groups of rats between 12:00-1:00 (between 1-2 h after the onset of the dark cycle) as described in the Methods section of the main text for the *Lactococcus* tissue and behavior cohorts.

#### Quantification of *Lactococcus lactis* in stomach and cecal samples via qPCR

A Taqman probe was made based on primer sequences for *Lactococcus lactis* validated for qPCR^31^. DNA was extracted from stomach and cecal samples using a QIAamp PowerFecal Pro DNA Kit (Cat. no. 51804, Qiagen) according to the manufacturer’s instructions. Total double-stranded DNA concentration per extracted sample was measured using a NanoDrop Spectrophotometer (ND-ONE-W, ThermoFisher Scientific). 30 ng DNA for stomach content and 1500 ng DNA for cecal content were used for qPCR, which was performed with Applied Biosystems TaqMan Fast Advanced Master Mix (Cat. no. 444557, ThermoFisher Scientific) and an Applied Biosystems QuantStudio 5 Real-Time PCR System (ThermoFisher Scientific). The thermal cycling protocol consisted of one 2-min cycle at each 50 °C and 90 °C followed by 40 cycles at 95 °C for 1 s and 60 °C for 20 s. A standard curve of *Lactococcus lactis* subspecies *lactis* strain NCTC 6681 (Lister, Schleifer et al., cat. no. 19435) with known concentration - grown as described above with CFUs quantified via plate counts with DNA extracted using the same QIAamp PowerFecal Pro DNA Kit indicated above - was used to determine absolute normalization of log CFUs per 5 ng DNA from stomach content and 250 ng DNA from cecal content.

#### Quantification of *Lactococcus lactis* subspecies *lactis* in diet samples via qPCR

DNA was extracted in triplicate from heat-treated and non-heat-treated diet samples according to previously established procedures^32^ with slight modifications. Briefly, diet samples (11 g per replicate) were mixed with sterile saline (0.9% w/v) and sodium citrate (25% w/v), and polyethylene glycol (PEG) 8000 was added to yield a concentration of 4% w/v PEG 8000 in the final mixture. Samples were then blended using a food processor for 2 min, and the resulting mixtures were centrifuged at 9700 × g for 15 min at 4°C. The supernatant was decanted and the remaining pellet was resuspended in DNAzol BD Reagent (Invitrogen, Carlsbad, CA, USA) in a volume ratio of 1:2 (pellet:DNA) with thorough mixing by vortex. Isopropanol (400 µL) was added to 1.5 mL of the resuspended pellet, and the resulting suspensions were mixed via vortex, centrifuged at 6000 × g, washed with DNAzol BD and ethanol (75% w/v), suspended in sterile deionized water (200 µL), and purified via centrifugation through a QIAshredder column for 2 min at 11750 × g (QIAGEN Inc., Valencia, CA, USA). Total double-stranded DNA concentration per extracted diet sample was measured using a NanoDrop Spectrophotometer (ND-ONE-W, ThermoFisher Scientific). 100 ng DNA was used for qPCR, conducted as described above for the quantification of *Lactococcus lactis* in stomach and cecal samples via qPCR with the exception that absolute normalization of log CFUs was calculated per 100 ng DNA for diet samples.

## Supplementary Discussion

Given the differences in gut microbial taxonomic composition between rats receiving the AGE-rich vs. control diets, we examined gut microbial metabolites. No differences were observed in short-chain fatty acid or lactate levels in serum, cecal content, or fecal content at postnatal day (PN) 56 (i.e., when the gut microbiome differed between groups, during the experimental diet period), although there was a statistically significant increase in fecal levels of the branched-chain fatty acid, isobutyrate (Extended Data Figure 5.3a-o). Elevated isobutyrate levels have previously been linked with increased protein fermentation^33,34^. It is possible that heat treatment ultimately decreased protein digestibility through the Maillard reaction, especially given that AGEs are glycated amino acids. Future research is needed to determine whether elevated gut isobutyrate levels may play a role in AGE-induced disturbances in gut barrier integrity or in the brain.

Although we found that consumption of the heat-treated AGE-rich diet promoted levels of AGEs in serum, it is unknown specifically how they entered into circulation. Given previous evidence linking prolonged dietary AGE consumption and increased intestinal permeability in rodents^35^, high early life dietary AGE exposure could also be disrupting intestinal barrier function and thereby allowing more AGEs, and even physiologically active bacterial products such as LPS, to pass into the bloodstream. Once in circulation, the majority of low molecular weight AGEs are sufficiently small (<400 Da) to pass through the blood-brain barrier^36,37^, which could enable them to act directly in the hippocampus. Vice-versa, other gut-derived proinflammatory molecules triggered by a high AGE diet might indirectly act on the hippocampus^38^. While we only observed altered expression of the tight junction protein claudin-5 in the duodenum region of the small intestine (Extended Data Fig. 3d-k), we did not measure tight junction proteins in the ileum and colon, which arguably could be sites of greater action between AGEs and the gut microbiota. Future work is needed to determine whether dietary AGEs, especially when unbound to proteins and of relatively small molecular weight, can pass into the brain.

We initially hypothesized that an over-pruning of synapses would likely be giving rise to the memory impairments we observed for rats receiving the AGE-rich diet, which would have been marked by an over-activation (and thus higher expression) of one or more components of the complement system. However, our findings of decreased expression of C3 in the hippocampus suggest that complement system expression/activation may follow an inverted U-shaped curve, with extremes of *either too high* or *too low* of activation negatively impacting synaptic pruning and behavioral phenotypes. Yet another explanation is that additional factor(s) are at play. The receptor for AGEs (RAGE) is expressed in various locations throughout the body and has been implicated in inflammatory responses, although mounting evidence indicates that it has numerous ligands beyond AGEs^39–41^. We did not observe differences in expression of RAGE or nuclear factor kappa-light-chain-enhancer of activated B cells (NFĸB) in the dorsal hippocampus, although we did observe a trending decrease in expression of postsynaptic density protein 95 (PSD-95) in rats that received the AGE-rich diet (P=0.0532; Extended Data Figure 3a-c). These findings continue to point to the complement system as a mediator of the early life AGE diet-induced memory deficits observed.

---

**Extended Data Figure 1.**
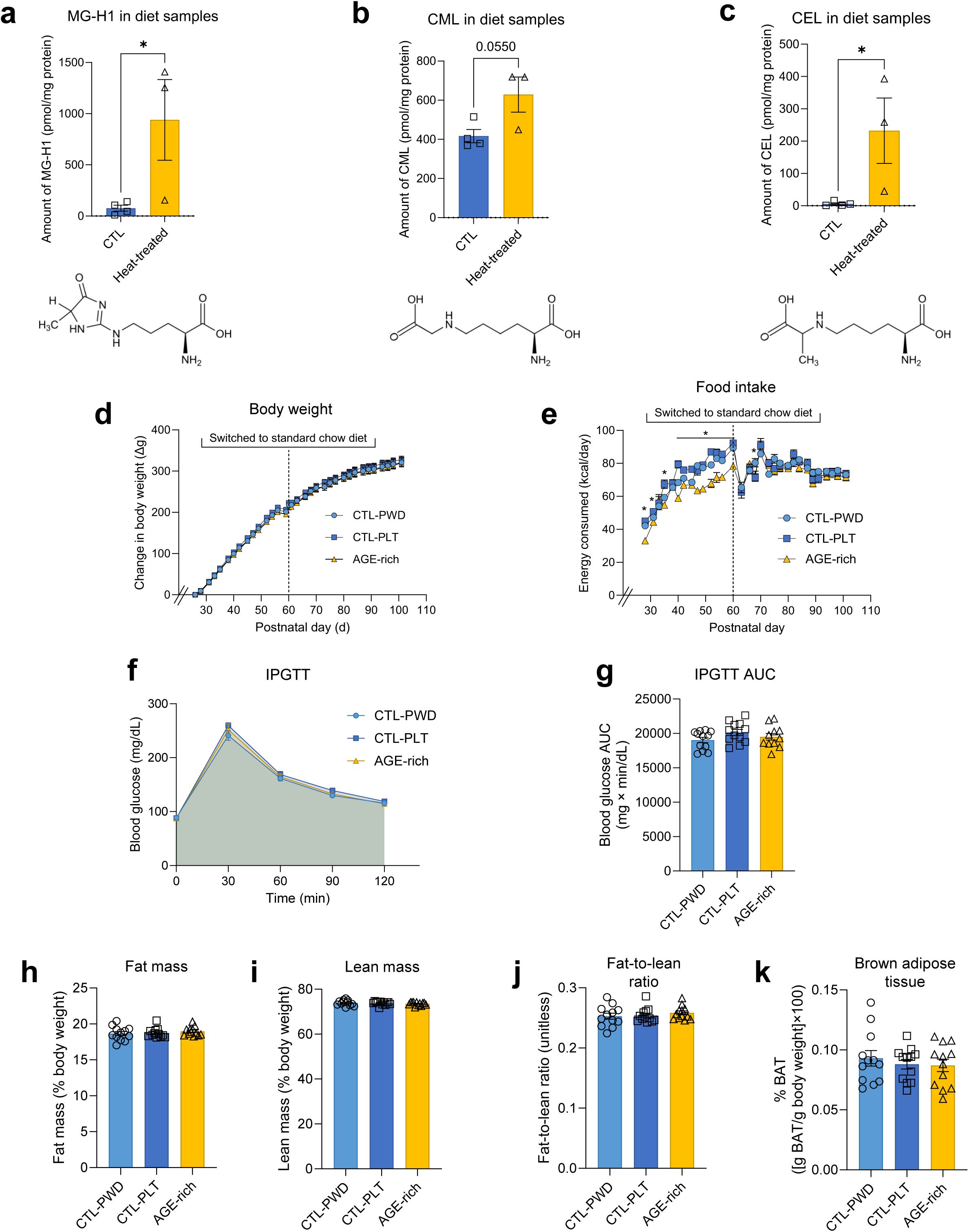
Heat treatment of AIN-93G diet increases levels of advanced glycation end-products, and early life consumption of such heat-treated diet does not affect body weight trajectory, glucose tolerance, or body composition compared to powdered and pelleted control AIN-93G diets. **a**, Concentration of MG-H1 in heat-treated vs. control non-heat-treated pelleted AIN-93G diets (n=4 CTL, n=3 Heat-treated; unpaired two-tailed t-test, P=0.048). The structure of MG-H1 is also shown. **b**, Concentration of CML in heat-treated vs. control pelleted AIN-93G diets (n=4 CTL, n=3 Heat-treated; unpaired two-tailed t-test, P=0.0550). The structure of CML is also shown. **c**, Concentration of CEL in heat-treated vs. control pelleted AIN-93G diets (n=4 CTL, n=3 Heat-treated; unpaired two-tailed t-test, P=0.0445). The structure of CEL is also shown. **d**, Change in body weight over time in rats receiving the CTL-PWD, CTL-PLT, and AGE-rich diets from PN 26-60 followed by standard chow (n=12 CTL-PWD, n=12 CTL-PLT, n=12 AGE-rich; 2-way repeated measures ANOVA, diet group [P=0.6272], time [repeated, P<0.0001], diet group × time [P>0.9999]). **e**, Food intake (kcal) over time in rats receiving the CTL-PWD, CTL-PLT, and AGE-rich diets from PN 26-60 followed by standard chow (n=12 CTL-PWD, n=12 CTL-PLT, n=12 AGE-rich; 2-way repeated measures ANOVA, diet group [P=0.0144], time [repeated, P<0.0001], diet group × time [P<0.0001]; various instances of statistically significant differences indicated by Tukey’s post-hoc multiple comparisons test). **f**, Blood glucose values over time from IPGTT performed on PN 59 (n=12 CTL-PWD, n=12 CTL-PLT, n=11 AGE rich; 2-way repeated measures ANOVA, diet group [P=0.2273], time [repeated, P<0.0001], diet group × time [P=0.7087]). **g**, Blood glucose AUC from IPGTT performed on PN 59 (n=12 CTL-PWD, n=12 CTL-PLT, n=11 AGE-rich; 1-way ANOVA, P=0.2123). **h**, Body fat mass on PN 60 (n=12 CTL-PWD, n=12 CTL-PLT, n=12 AGE-rich; 1-way ANOVA, P=0.5795). **i**, Body lean mass on PN 60 (n=12 CTL-PWD, n=12 CTL-PLT, n=12 AGE-rich; 1-way ANOVA, P=0.6140). **j**, Fat-to-lean ratio on PN 60 (n=12 CTL-PWD, n=12 CTL-PLT, n=12 AGE-rich; 1-way ANOVA, P=0.6045). **k**, Brown adipose tissue on PN 102 [after the standard chow exposure period during adulthood] (n=12 CTL-PWD, n=12 CTL-PLT, n=12 AGE-rich; 1-way ANOVA, P=0.7017). Error bars represent standard error of the mean (SEM). *P<0.05. All n’s indicate number of rats per group. Additional details about the statistical analyses for each subpanel can be found in Supplementary Table S1. AGE, advanced glycation end-product; ANOVA, analysis of variance; AUC, area under the curve; BAT, brown adipose tissue; CEL, carboxyethyllysine; CML, carboxymethyllysine; CTL-PLT, control pelleted [diet]; CTL-PWD, control powdered [diet]; IPGTT, intraperitoneal glucose tolerance test; MG-H1, methylglyoxal-derived hydroimidazolone; PN, postnatal day.

**Extended Data Figure 2.**
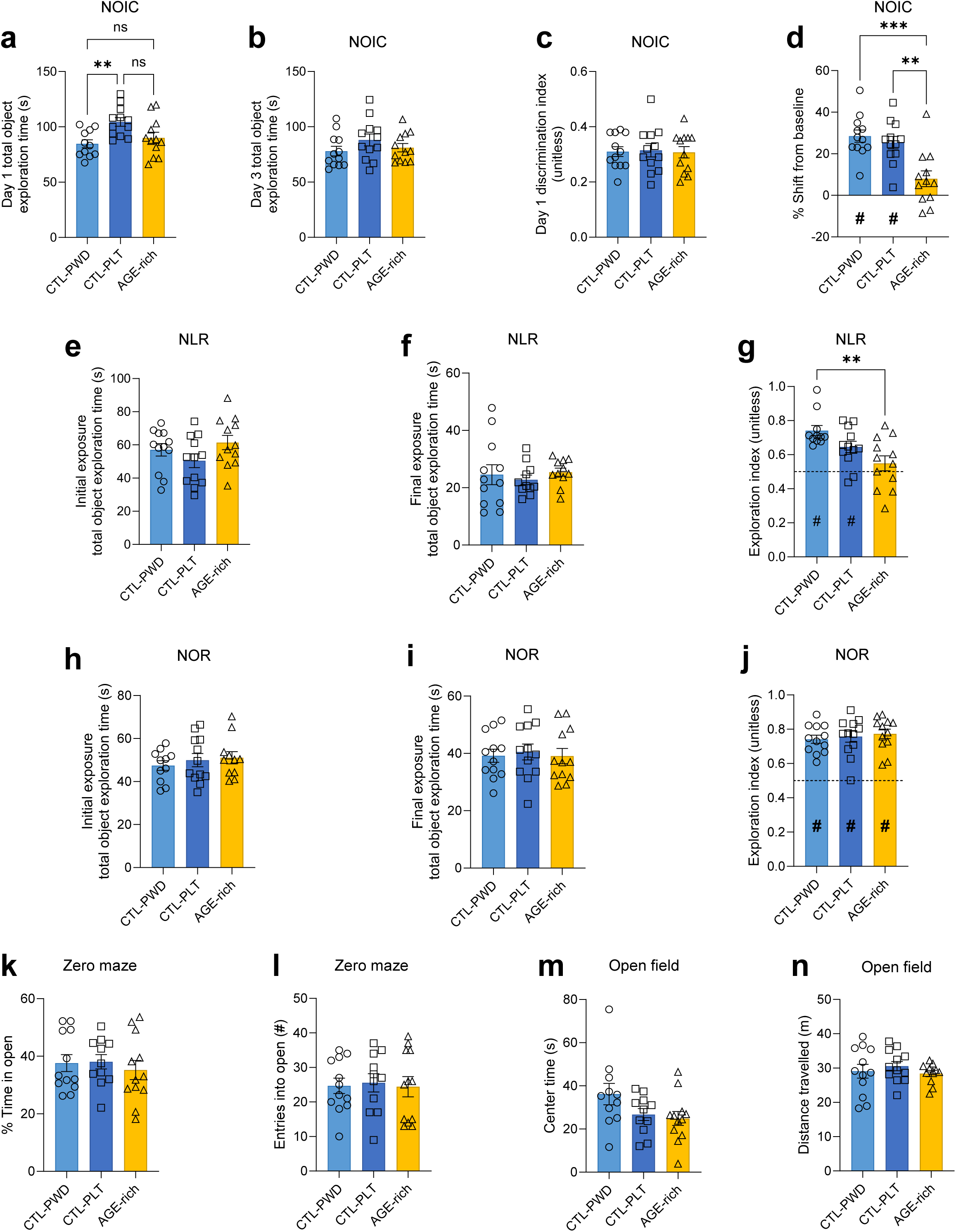
Consumption of a heat-treated AGE-rich diet during early life compared to non-heat-treated powdered and pelleted control diets leads to hippocampal-dependent memory impairments in adulthood, without impacting perirhinal cortex-dependent memory, anxiety-like behavior, or locomotor activity. **a**, Total object exploration time during NOIC day 1 (n=11 CTL-PWD, n=12 CTL-PLT, n=12 AGE-rich; 1-way ANOVA, P=0.0071; post-hoc Tukey’s multiple comparisons – CTL-PWD vs. CTL-PLT: P=0.0070, CTL-PWD vs. AGE-rich: P=0.6477, CTL-PLT vs. AGE-rich: P=0.0530). **b**, Total object exploration time during NOIC day 3 (n=12 CTL-PWD, n=12 CTL-PLT, n=12 AGE-rich; 1-way ANOVA, P=0.2834). **c**, NOIC day 1 object discrimination index (n=12 CTL-PWD, n=12 CTL-PLT, n=12 AGE-rich; 1-way ANOVA, P=0.9679). **d**, NOIC percent shift from baseline object exploration (n=12 CTL-PWD, n=12 CTL-PLT, n=12 AGE-rich; 1-way ANOVA, P=0.0002; post-hoc Tukey’s multiple comparisons – CTL-PWD vs. CTL-PLT: P=0.7816, CTL-PWD vs. AGE-rich: P=0.0003, CTL-PLT vs. AGE-rich: P=0.0019; one-sample t-test for difference from 0 [chance], P<0.0001 CTL-PWD, P<0.0001 CTL-PLT, P=0.06 AGE-rich). **e**, Total object exploration time during the initial exposure period for NLR (n=12 CTL-PWD, n=12 CTL-PLT, n=12 AGE-rich; 1-way ANOVA, P=0.1778). **f**, Total object exploration time during the final exposure period for NLR (n=12 CTL-PWD, n=11 CTL-PLT, n=11 AGE-rich; Brown-Forsythe ANOVA, P=0.4939). **g**, NLR object exploration index during the test exposure period (n=11 CTL-PWD, n=12 CTL-PLT, n=12 AGE-rich; 1-way ANOVA, P=0.0034; post-hoc Tukey’s multiple comparisons – CTL-PWD vs. CTL-PLT: P=0.1737, CTL-PWD vs. AGE-rich: P=0.0023, CTL-PLT vs. AGE-rich: P=0.1551; one-sample t-test for difference from 0.5 [chance], P<0.0001 CTL-PWD, P=0.0011 CTL-PLT, P=0.2834 AGE-rich). **h**, Total object exploration time during the initial exposure period for NOR (n=11 CTL-PWD, n=12 CTL-PLT, n=11 AGE-rich; 1-way ANOVA, P=0.6472). **i**, Total object exploration time during the final exposure period for NOR (n=12 CTL-PWD, n=12 CTL-PLT, n=11 AGE-rich; 1-way ANOVA, P=0.9070). **j**, NOR object exploration index during the test exposure period (n=12 CTL-PWD, n=12 CTL-PLT, n=12 AGE-rich; 1-way ANOVA, P=0.7356; one-sample t-test for difference from 0.5 [chance], P<0.0001 CTL-PWD, P<0.0001 CTL-PLT, P<0.0001 AGE-rich). **k**, Percent time in the open arm zones of the zero maze apparatus (n=12 CTL-PWD, n=11 CTL-PLT, n=12 AGE-rich; 1-way ANOVA, P=0.7155). **l**, Number of entries into the open arm zones of the zero maze apparatus (n=12 CTL-PWD, n=11 CTL-PLT, n=12 AGE-rich; 1-way ANOVA, P=0.9515). **m**, Time spent in the center zone during the open field test (n=11 CTL-PWD, n=11 CTL-PLT, n=12 AGE-rich; 1-way ANOVA, P=0.0881). **n**, Distance travelled during the open field test (n=12 CTL-PWD, n=12 CTL-PLT, n=11 AGE-rich; Brown-Forsythe ANOVA, P=0.4369). Error bars represent standard error of the mean (SEM). **P<0.01, ***P<0.001, # indicates statistically significant difference from chance (P<0.05), which was 0.0 for NOIC and 0.5 for NLR and NOR. All n’s indicate number of rats per group. Additional details about the statistical analyses for each subpanel can be found in Supplementary Table S1. AGE, advanced glycation end-product; ANOVA, analysis of variance; CTL-PLT, control pelleted [diet]; CTL-PWD, control powdered [diet]; NLR, novel location recognition; NOIC, novel object in context; NOR, novel object recognition; PN, postnatal day.

**Extended Data Figure 3.**
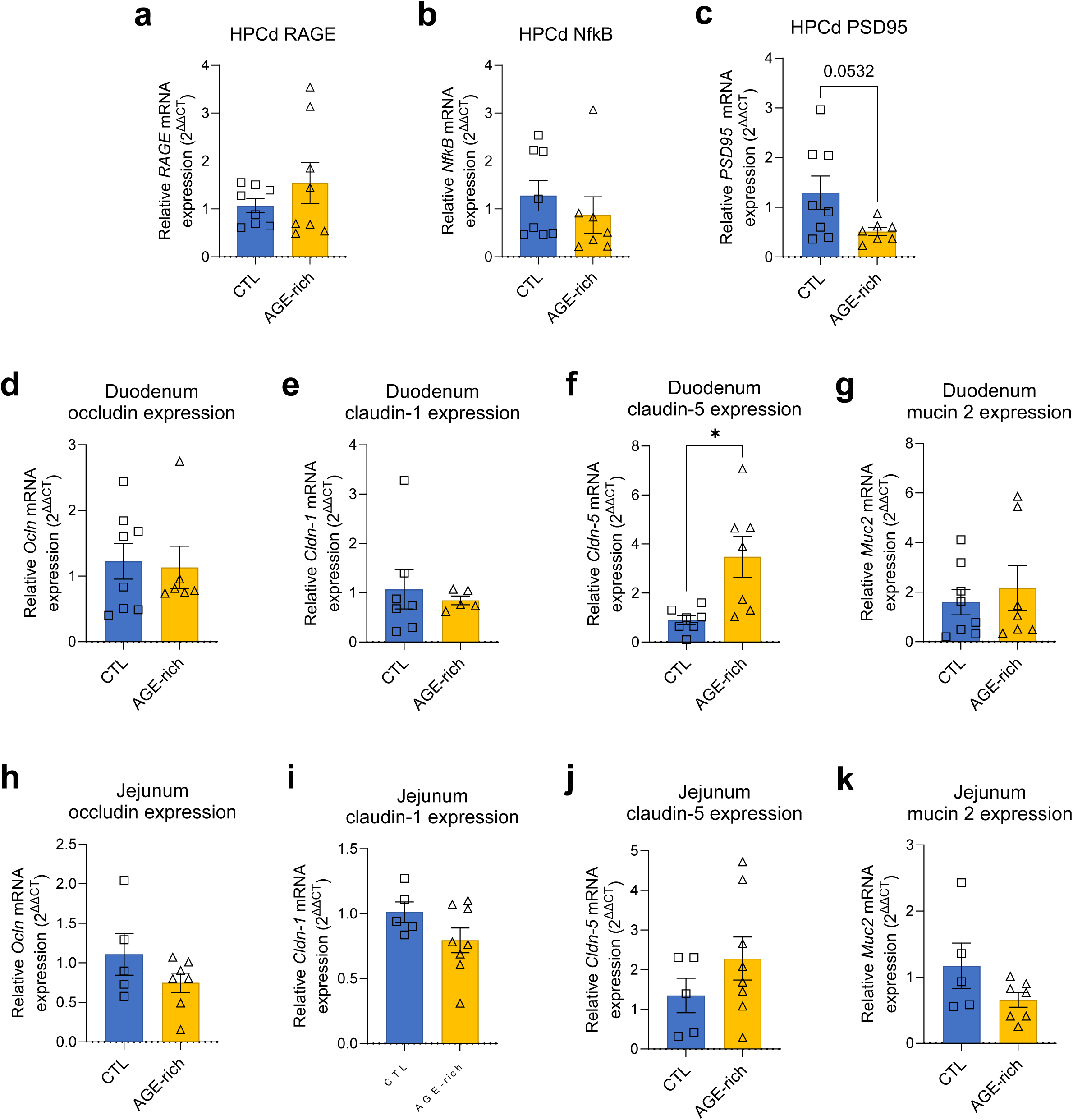
Early life consumption of an AGE-rich diet leads to trending-to-no differences in expression of RAGE and, NFĸB, and PSD95 in the dorsal hippocampus and minor differences in expression of tight junction proteins in the small intestine. **a,** mRNA expression of *RAGE* in the HPCd (n=8 CTL, n=8 AGE-rich; unpaired two-tailed t-test, P=0.3075). **b,** mRNA expression of *NFĸB* in the HPCd (n=8 CTL, n=7 AGE-rich; unpaired two-tailed t-test, P=0.4285). **c,** mRNA expression of *PSD95* in the HPCd (n=8 CTL, n=7 AGE-rich; unpaired two-tailed t-test, P=0.0532). **d,** mRNA expression of *occludin* in the duodenum (n=8 CTL, n=6 AGE-rich; unpaired two-tailed t-test, P=0.8278). **e,** mRNA expression of *claudin-1* in the duodenum (n=7 CTL, n=5 AGE-rich; unpaired two-tailed t-test, P=0.6547). **f,** mRNA expression of *claudin-5* in the duodenum (n=7 CTL, n=7 AGE-rich; unpaired two-tailed t-test, P=0.0111). **g,** mRNA expression of *mucin 2* in the duodenum (n=8 CTL, n=7 AGE-rich; unpaired two-tailed t-test, P=0.5817). **h,** mRNA expression of *occludin* in the jejunum (n=5 CTL, n=7 AGE-rich; unpaired two-tailed t-test, P=0.1992). **i,** mRNA expression of *claudin-1* in the jejunum (n=5 CTL, n=7 AGE-rich; unpaired two-tailed t-test, P=0.1429). **j,** mRNA expression of *claudin-5* in the jejunum (n=5 CTL, n=7 AGE-rich; unpaired two-tailed t-test, P=0.2541). **k,** mRNA expression of *mucin 2* in the jejunum (n=5 CTL, n=8 AGE-rich; unpaired two-tailed t-test, P=0.1327). Error bars represent standard error of the mean (SEM). *P<0.05. All n’s indicate number of rats per group. Additional details about the statistical analyses for each subpanel can be found in Supplementary Table S1. AGE, advanced glycation end-product; CTL, control [diet]; mRNA, messenger RNA; NFĸB, nuclear factor kappa-light-chain-enhancer of activated B cells; PN, postnatal day; PSD95, postsynaptic density protein-95; RAGE, receptor for advanced glycation end-products.

**Extended Data Figure 4.**
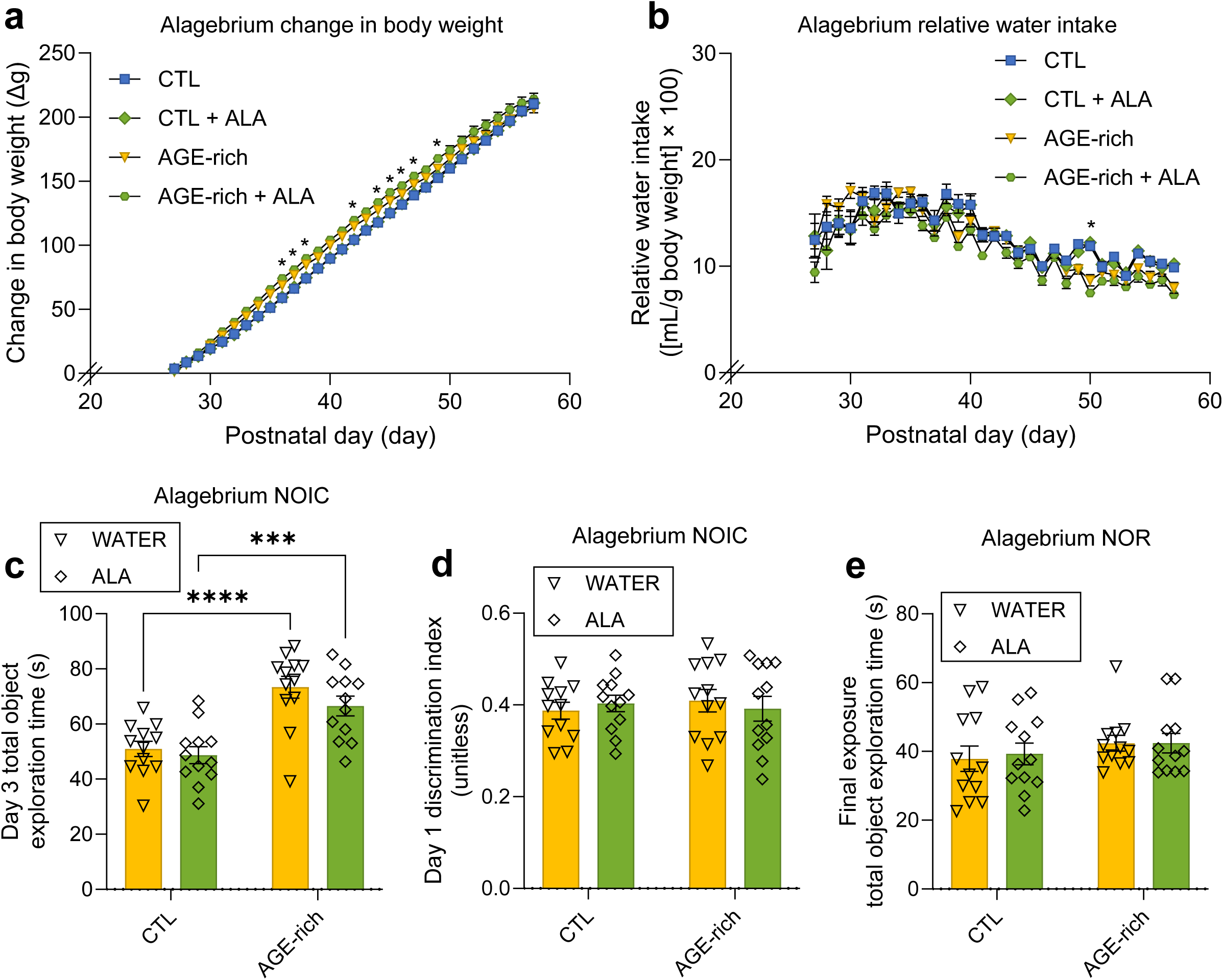
Body weight, food intake, and expanded NOIC behavioral data for rats receiving the AGE-inhibitor drug alagebrium (ALA) in early life with AGE-rich and control diets. **a,** Change in body weight over time in rats receiving the CTL and AGE-rich diets with and without ALA from PN 26-60 (n=12 CTL w/water, n=12 CTL w/ALA, n=12 AGE-rich w/water, n=12 AGE-rich w/ALA; 3-way ANOVA with factors of diet group, drug [water vs. ALA], time, diet group × drug, diet group × time, drug × time, diet group × drug × time; P=0.0001 diet group, P=0.1753 drug [water vs. ALA], P<0.0001 time, P=0.2413 diet group × drug, P<0.0001 diet group × time, P=0.7737 drug × time, P=0.9948 diet group × drug × time. **b,** Relative water intake over time in rats receiving the CTL and AGE-rich diets with and without ALA from PN 26-60 (n=12 CTL w/water, n=12 CTL w/ALA, n=12 AGE-rich w/water, n=12 AGE-rich w/ALA; 3-way ANOVA with factors of diet group, drug [water vs. ALA], time, diet group × drug, diet group × time, drug × time, diet group × drug × time; P=0.0309 diet group, P=0.1262 drug [water vs. ALA], P<0.0001 time, P=0.3110 diet group × drug, P<0.0001 diet group × time, P=0.9290 drug × time, P=0.9056 diet group × drug × time. **c,** Total object exploration time during NOIC day 3 (n=12 CTL w/water, n=12 CTL w/ALA, n=12 AGE-rich w/water, n=12 AGE-rich w/ALA; 2-way ANOVA with factors of diet group, drug [water vs. ALA], and diet group × drug interaction; P<0.0001 diet group, P=0.2087 drug, P=0.4891 diet group × drug; post-hoc test results for multiple comparisons using uncorrected Fisher’s LSD – CTL water vs. ALA: P=0.6388, AGE-rich water vs. ALA: P=0.1602, water CTL vs. AGE-rich: P<0.0001, ALA CTL vs. AGE-rich: P=0.0008). **d,** NOIC day 1 discrimination index (n=12 CTL w/water, n=12 CTL w/ALA, n=12 AGE-rich w/water, n=12 AGE-rich w/ALA; 2-way ANOVA with factors of diet group, drug [water vs. ALA], and diet group × drug interaction; P=0.8468 diet group, P=0.9620 drug, P=0.5381 diet group × drug). **e,** NOR total object exploration time during the final exposure period (n=12 CTL w/water, n=12 CTL w/ALA, n=12 AGE-rich w/water, n=12 AGE-rich w/ALA; 2-way ANOVA with factors of diet group, drug [water vs. ALA], and diet group × drug interaction; P=0.0837 diet group, P=0.8493 drug, P=0.7392 diet group × drug). Error bars represent standard error of the mean (SEM).*P<0.05, ***P<0.001, ****P<0.0001. # indicates statistically significant difference from chance (P<0.05), which was 0.0 for NOIC and 0.5 for NOR. All n’s indicate number of rats per group. Additional details about the statistical analyses for each subpanel can be found in Supplementary Table S1. Additional details about the statistical analyses for each subpanel can be found in Supplementary Table S1. AGE, advanced glycation end-product; ALA, alagebrium; ANOVA, analysis of variance; CTL, control [diet]; NOIC, novel object in context; NOR, novel object recognition; PN, postnatal day.

**Extended Data Figure 5.1.**
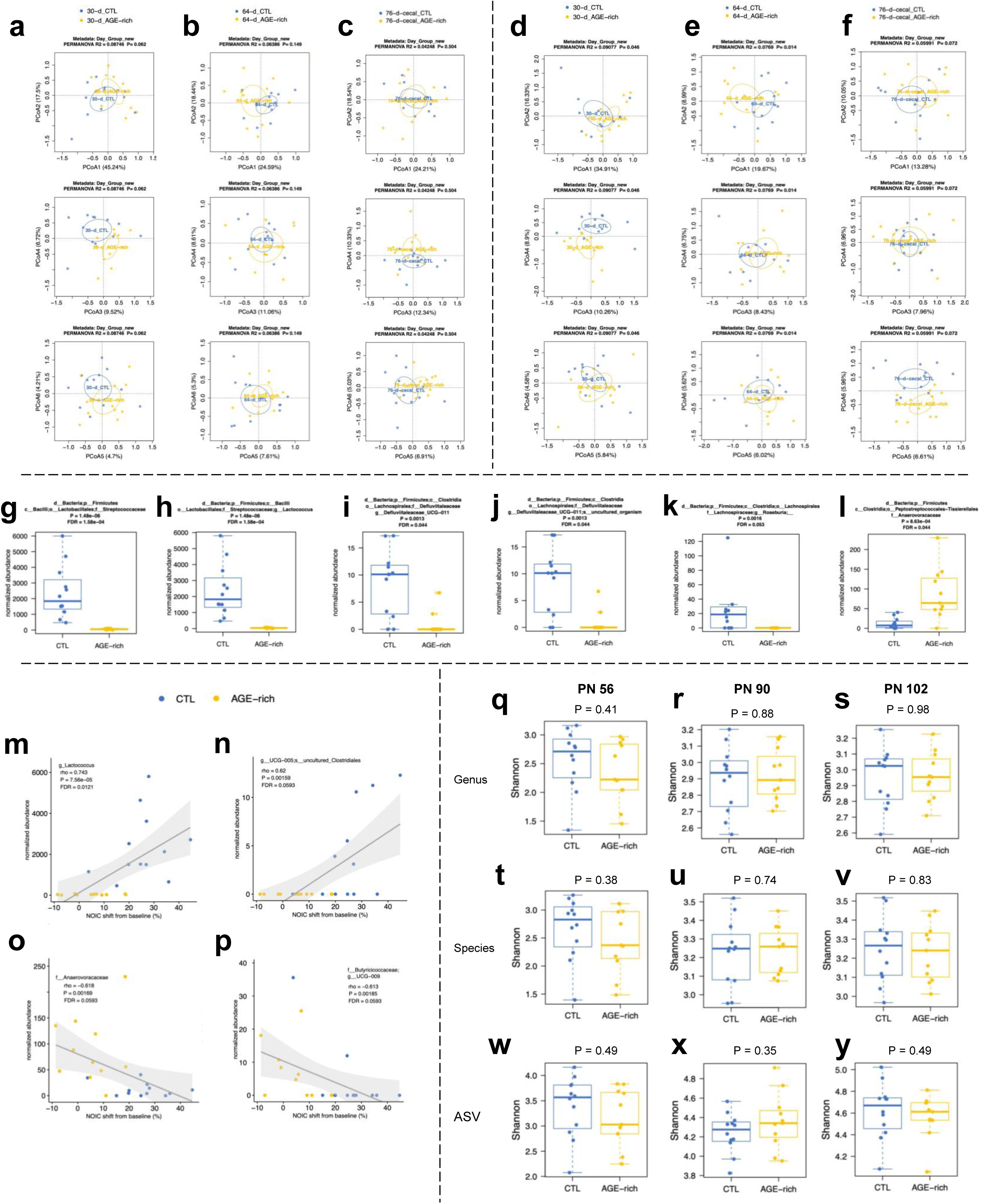
Gut microbiome changes due to early life consumption of an AGE-rich diet. **a**, Microbiome principal coordinate analysis (PCoA) plot for the genus level from fecal samples collected at PN 56 in rats that consumed a heat-treated AGE-rich diet vs. non-heat-treated diet during early life (n=12 CTL, n=12 AGE-rich; Bray-Curtis dissimilarity; R^2^=0.08746, P=0.062). **b**, Microbiome PCoA plot for the genus level from fecal samples collected at PN 90 in rats that consumed a heat-treated AGE-rich diet vs. non-heat-treated diet during early life (n=12 CTL, n=12 AGE-rich; Bray-Curtis dissimilarity; R^2^=0.06386, P=0.149). **c**, Microbiome PCoA plot for the genus level from fecal samples collected at PN 102 in rats that consumed a heat-treated AGE-rich diet vs. non-heat-treated diet during early life (n=12 CTL, n=12 AGE-rich; Bray-Curtis dissimilarity; R^2^=0.09077, P=0.046). **d**, Microbiome PCoA plot for the ASV level from fecal samples collected at PN 56 in rats that consumed a heat-treated AGE-rich diet vs. non-heat-treated diet during early life (n=12 CTL, n=12 AGE-rich; Bray-Curtis dissimilarity; R^2^=0.08746, P=0.062). **e**, Microbiome PCoA plot for the ASV level from fecal samples collected at PN 90 in rats that consumed a heat-treated AGE-rich diet vs. non-heat-treated diet during early life (n=12 CTL, n=12 AGE-rich; Bray-Curtis dissimilarity; R^2^=0.0769, P=0.014). **f**, Microbiome PCoA plot for the ASV level from fecal samples collected at PN 102 in rats that consumed a heat-treated AGE-rich diet vs. non-heat-treated diet during early life (n=12 CTL, n=12 AGE-rich; Bray-Curtis dissimilarity; R^2^=0.05991, P=0.072). **g**, *Streptococcaceae* (family) abundance in fecal samples at PN 56 (n=12 CTL, n=12 AGE-rich; Non-parametric Wilcoxon rank sum test; P=1.48×10^-6^, FDR=1.58×10^-4^). **h**, *Lactococcus* (genus) abundance in fecal samples at PN 56 (n=12 CTL, n=12 AGE-rich; Non-parametric Wilcoxon rank sum test; P=1.48×10^-6^, FDR=1.58×10^-4^). **i**, *Defluviitaleaceae_UCG-011* (genus) abundance in fecal samples at PN 56 (n=12 CTL, n=12 AGE-rich; Non-parametric Wilcoxon rank sum test; P=0.0013, FDR=0.044). **j**, Uncultured organism (species) abundance in fecal samples at PN 56 (n=12 CTL, n=12 AGE-rich; Non-parametric Wilcoxon rank sum test; P=0.0013, FDR=0.044). **k**, *Roseburia* (genus) abundance in fecal samples at PN 56 (n=12 CTL, n=12 AGE-rich; Non-parametric Wilcoxon rank sum test; P=0.0016, FDR=0.053). **l**, *Anaerovoracaceae* (family) abundance in fecal samples at PN 56 (n=12 CTL, n=12 AGE-rich; Non-parametric Wilcoxon rank sum test; P=8.63×10^-4^, FDR=0.044). **m**, Correlation between NOIC performance and *Lactococcus* (genus) abundance (n=12 CTL, n=12 AGE-rich; Spearman’s correlation; rho=0.743, P=7.56×10^-6^, FDR=0.0121). **n**, Correlation between NOIC performance and *UCG-005* (genus) uncultured *Clostridiales* (species) abundance (n=12 CTL, n=12 AGE-rich; Spearman’s correlation; rho=0.62, P=0.00159, FDR=0.0593). **o**, Correlation between NOIC performance and *Anaerovoracaceae* (family) abundance (n=12 CTL, n=12 AGE-rich; Spearman’s correlation; rho=-0.618, P=0.00169, FDR=0.0593). **p**, Correlation between NOIC performance and *UCG-005* (genus) abundance (n=12 CTL, n=12 AGE-rich; Spearman’s correlation; rho=-0.613, P=0.00185, FDR=0.0593). **q**, Shannon index for genus level abundances at PN 56 (n=12 CTL, n=12 AGE-rich; Shannon index (alpha-diversity); P=0.41). **r**, Shannon index for genus level abundances at PN 90 (n=12 CTL, n=12 AGE-rich; Shannon index (alpha-diversity); P=0.88). **s**, Shannon index for genus level abundances at PN 102 (n=12 CTL, n=12 AGE-rich; Shannon index (alpha-diversity); P=0.98). **t**, Shannon index for species level abundances at PN 56 (n=12 CTL, n=12 AGE-rich; Shannon index (alpha-diversity); P=0.38). **u**, Shannon index for species level abundances at PN 90 (n=12 CTL, n=12 AGE-rich; Shannon index (alpha-diversity); P=0.74). **v**, Shannon index for species level abundances at PN 102 (n=12 CTL, n=12 AGE-rich; Shannon index (alpha-diversity); P=0.83). **w**, Shannon index for ASV level abundances at PN 56 (n=12 CTL, n=12 AGE-rich; Shannon index (alpha-diversity); P=0.49). **x**, Shannon index for ASV level abundances at PN 90 (n=12 CTL, n=12 AGE-rich; Shannon index (alpha-diversity); P=0.35). **y**, Shannon index for ASV level abundances at PN 102 (n=12 CTL, n=12 AGE-rich; Shannon index (alpha-diversity); P=0.49). Error bars represent standard error of the mean (SEM).*P<0.05, ***P<0.001, ****P<0.0001. All n’s indicate number of rats or samples per group. Additional details about the statistical analyses for each subpanel can be found in Supplementary Table S1. AGE, advanced glycation end-product; ASV, amplicon sequence variant; CTL, control [diet]; FDR, false discovery rate; PN, postnatal day; PCoA, principal coordinate analysis.

**Extended Data Figure 5.2.**
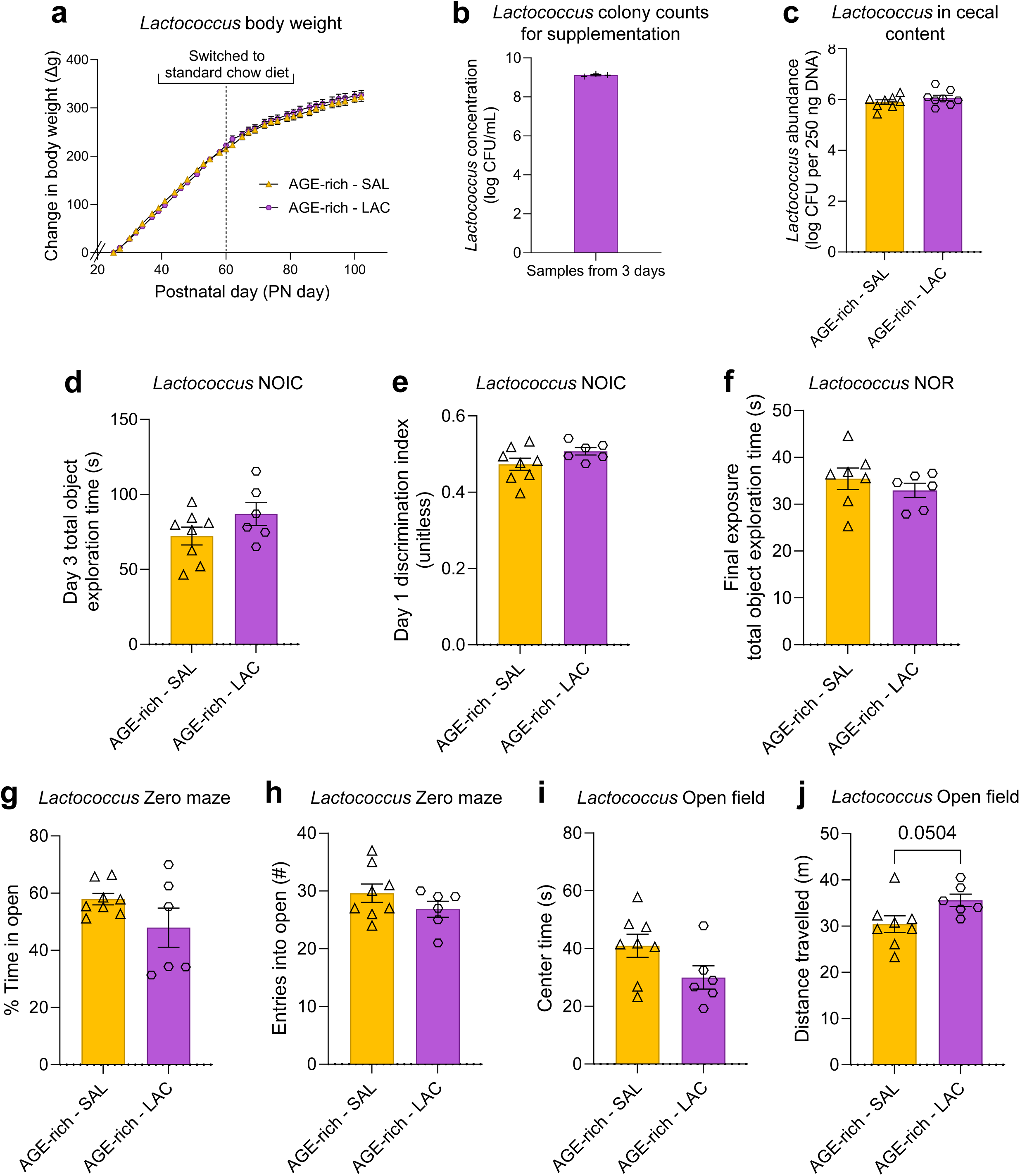
Early life supplementation with *Lactococcus lactis* subspecies *lactis* does not significantly alter body weight, total object exploration time, or anxiety-like behavior. **a**, Change in body weight over time in rats receiving the AGE-rich diet with or without Lactococcus supplementation (PN 26-41) from PN 26-60 followed by standard chow (n=8 AGE-rich - SAL, n=6 AGE-rich - LAC; 2-way ANOVA, supplement group [P=0.8345], time [P<0.0001], supplement group × time [P=0.0277]; no post-hoc comparisons were statistically significant). **b**, *Lactococcus lactis* subspecies *lactis* colony counts across supplementation days (count performed on 3 samples, no statistical analyses performed). **c**, *Lactococcus lactis* subspecies *lactis* abundance in cecal content (n=8 AGE-rich - SAL, n=6 AGE-rich - LAC; Welch’s t-test, P=0.2709). **d**, Total object exploration time during NOIC day 3 (n=8 AGE-rich - SAL, n=6 AGE-rich - LAC; unpaired two-tailed t-test, P=0.1477). **e**, Discrimination index during NOIC day 1 (n=8 AGE-rich - SAL, n=6 AGE-rich - LAC; unpaired two-tailed t-test, P=0.1198). **f**, Total object exploration time during the final exposure period of NOR (n=7 AGE-rich - SAL, n=6 AGE-rich - LAC; unpaired two-tailed t-test, P=0.4052). **g**, Percent time in the open arm zones during the zero maze test (n=7 AGE-rich - SAL, n=6 AGE-rich - LAC; Welch’s t-test, P=0.2131). **h**, Entries into the open arm zones during the zero maze test (n=8 AGE-rich - SAL, n=6 AGE-rich - LAC; Welch’s t-test, P=0.2303). **i**, Time spent in the center zone during the open field test (n=8 AGE-rich - SAL, n=6 AGE-rich - LAC; Welch’s t-test, P=0.0815). **j**, Distance travelled during the open field test (n=8 AGE-rich - SAL, n=6 AGE-rich - LAC; Welch’s t-test, P=0.0504). Error bars represent standard error of the mean (SEM). All n’s indicate number of rats per group. Additional details about the statistical analyses for each subpanel can be found in Supplementary Table S1. AGE, advanced glycation end-product; CTL, control [diet]; LAC, *Lactococcus*; NOIC, novel object in context; NOR, novel object recognition; PN, postnatal day; SAL, saline [vehicle].

**Extended Data Figure 5.3.**
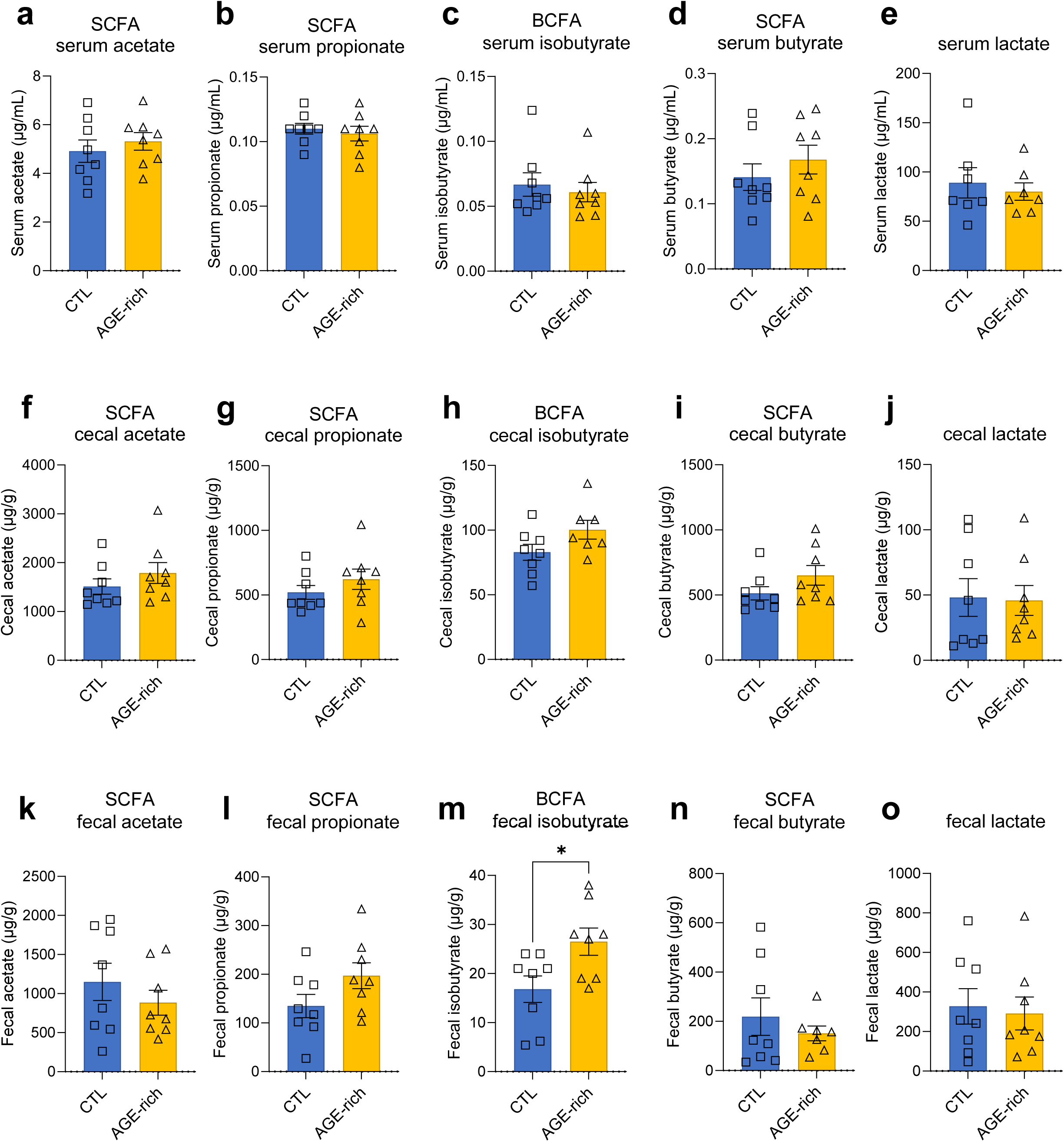
Effects of early life AGE-rich diet consumption on serum, cecal, and fecal levels of short-chain fatty acids (SCFAs). **a**, Serum acetate levels on PN 56 (n=8 CTL, n=8 AGE-rich; unpaired two-tailed t-test, P=0.5026). **b**, Serum propionate levels on PN 56 (n=8 CTL, n=8 AGE-rich; unpaired two-tailed t-test, P=0.6034). **c**, Serum isobutyrate levels on PN 56 (n=8 CTL, n=8 AGE-rich; unpaired two-tailed t-test, P=0.6222). **d**, Serum butyrate levels on PN 56 (n=8 CTL, n=8 AGE-rich; unpaired two-tailed t-test, P=0.3855). **e**, Serum lactate levels on PN 56 (n=7 CTL, n=7 AGE-rich; unpaired two-tailed t-test, P=0.6259). **f**, Cecal acetate levels on PN 56 (n=8 CTL, n=8 AGE-rich; unpaired two-tailed t-test, P=0.3129). **g**, Cecal propionate levels on PN 56 (n=8 CTL, n=8 AGE-rich; unpaired two-tailed t-test, P=0.3026). **h**, Cecal isobutyrate levels on PN 56 (n=8 CTL, n=7 AGE-rich; unpaired two-tailed t-test, P=0.0881). **i**, Cecal butyrate levels on PN 56 (n=8 CTL, n=8 AGE-rich; unpaired two-tailed t-test, P=0.1526). **j**, Cecal lactate levels on PN 56 (n=8 CTL, n=8 AGE-rich; unpaired two-tailed t-test, P=0.9045). **k**, Fecal acetate levels on PN 55 (n=8 CTL, n=8 AGE-rich; unpaired two-tailed t-test, P=0.3699). **l**, Fecal propionate levels on PN 55 (n=8 CTL, n=8 AGE-rich; unpaired two-tailed t-test, P=0.1021). **m**, Fecal isobutyrate levels on PN 55 (n=8 CTL, n=8 AGE-rich; unpaired two-tailed t-test, P=0.0253). **n**, Fecal butyrate levels on PN 55 (n=8 CTL, n=8 AGE-rich; unpaired two-tailed t-test, P=0.4452). **o**, Fecal lactate levels on PN 55 (n=8 CTL, n=8 AGE-rich; unpaired two-tailed t-test, P=0.7688). Error bars represent standard error of the mean (SEM).*P<0.05. All n’s indicate number of rats or samples per group. Additional details about the statistical analyses for each subpanel can be found in Supplementary Table S1. AGE, advanced glycation end-product; CTL, control [diet]; PN, postnatal day; SCFA, short-chain fatty acid.

**Table S1.**
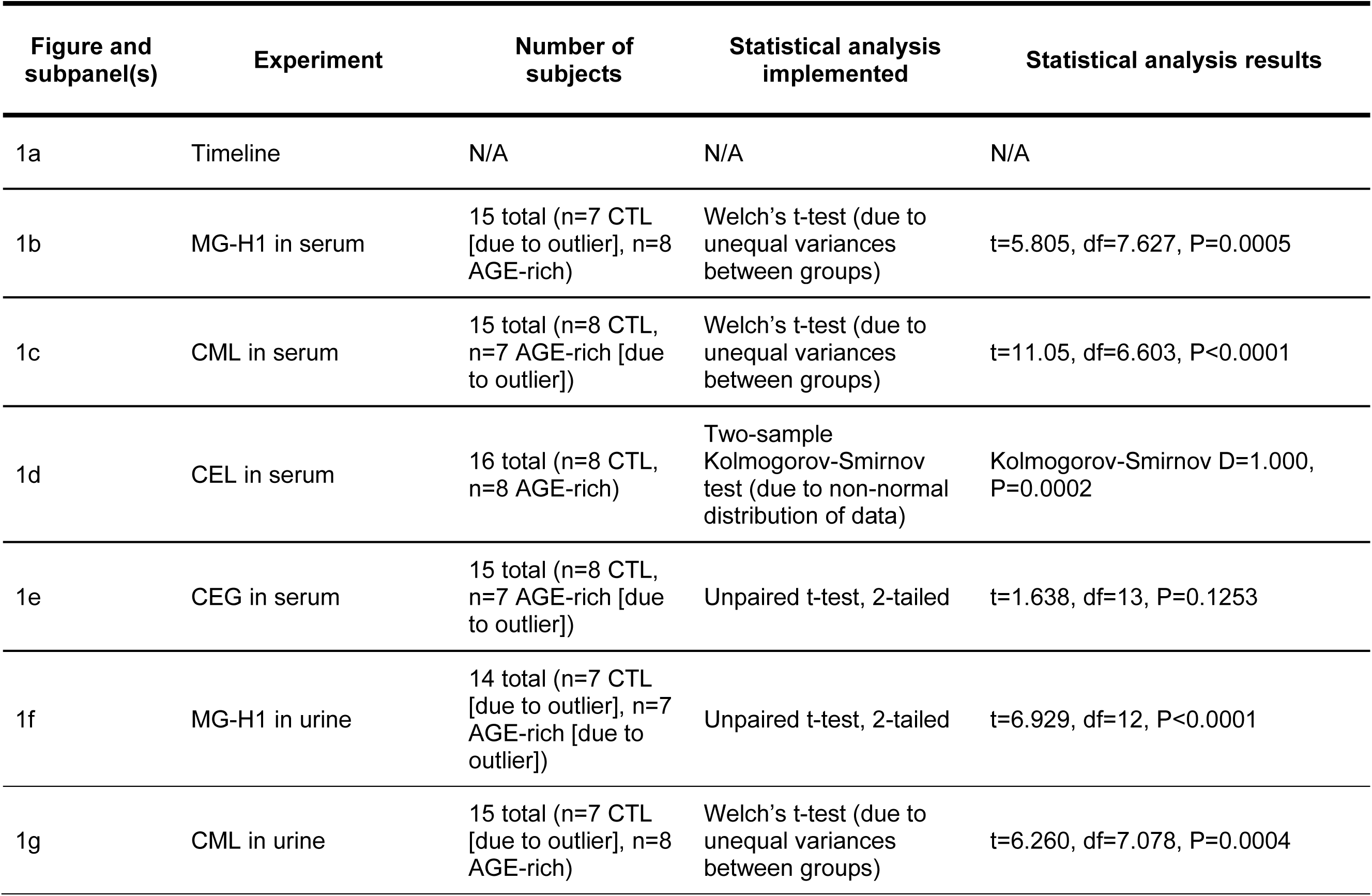
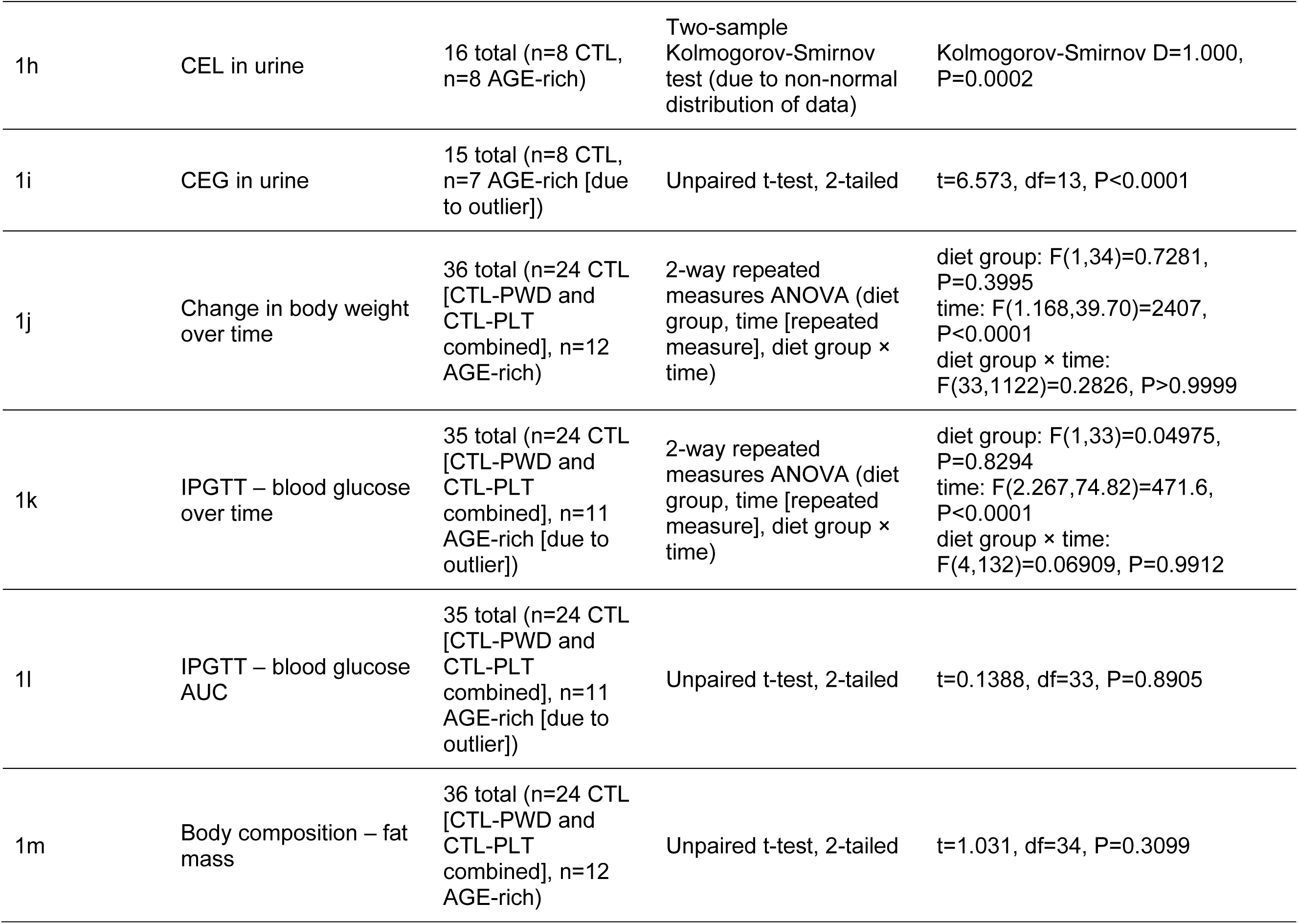
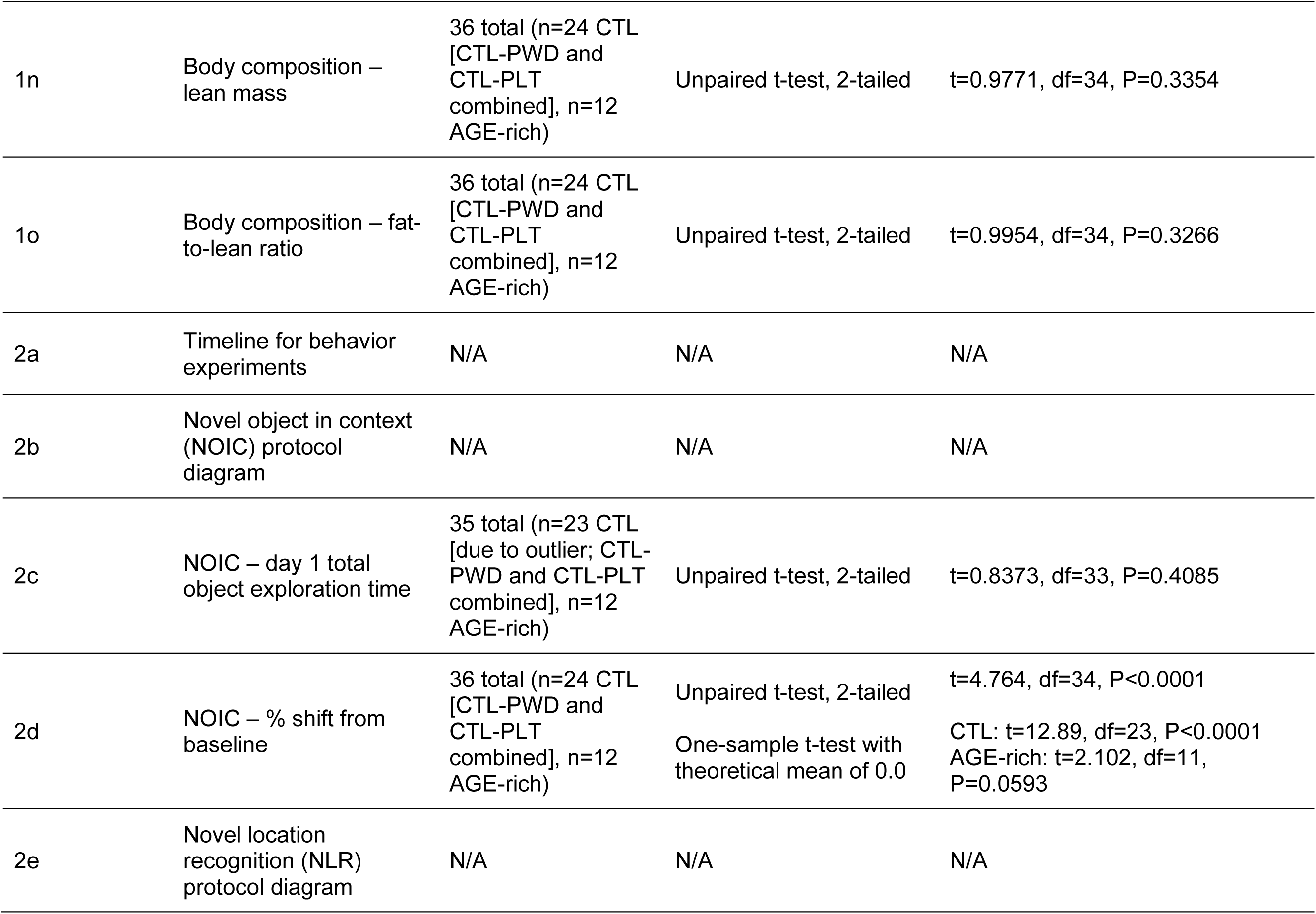
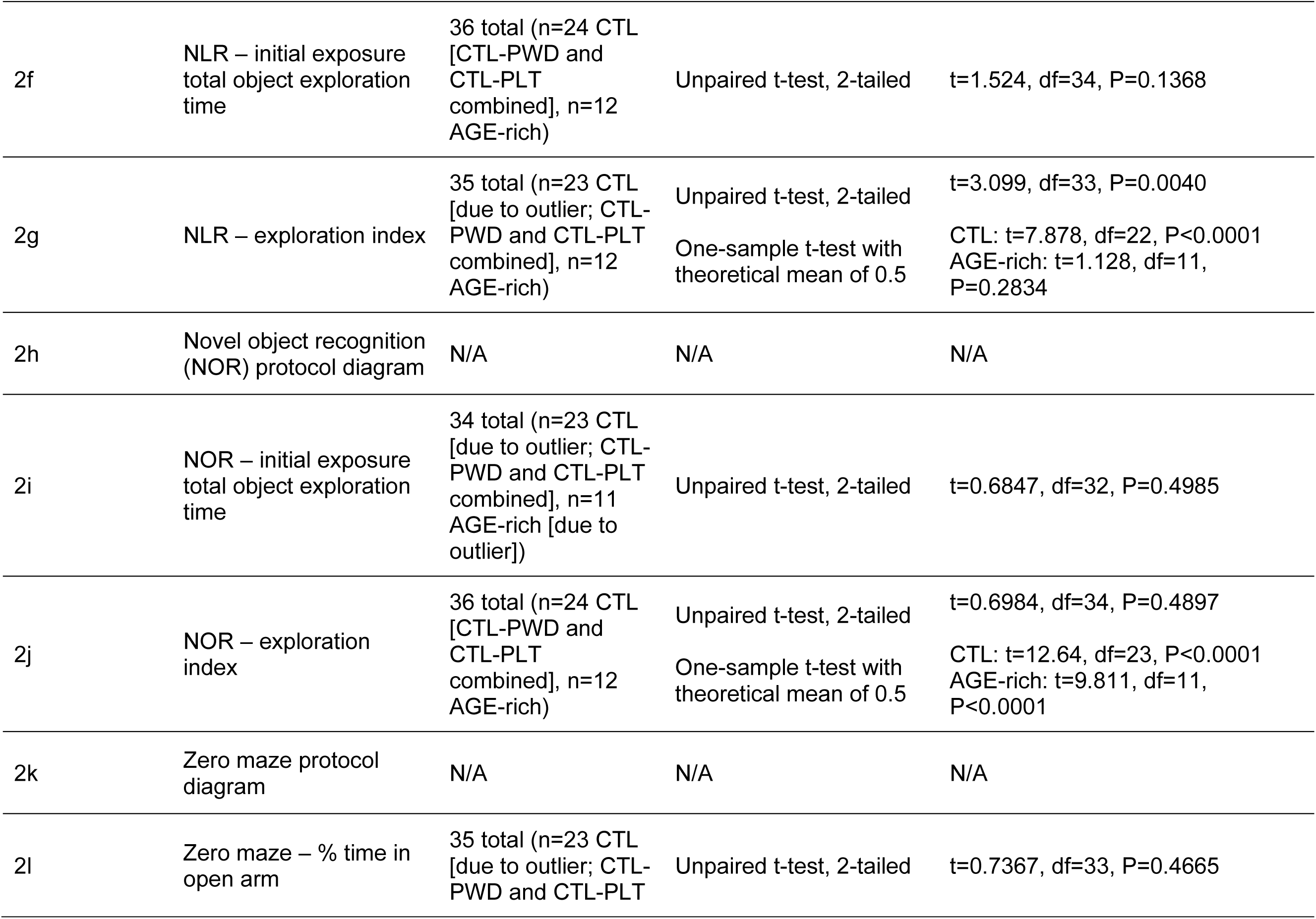
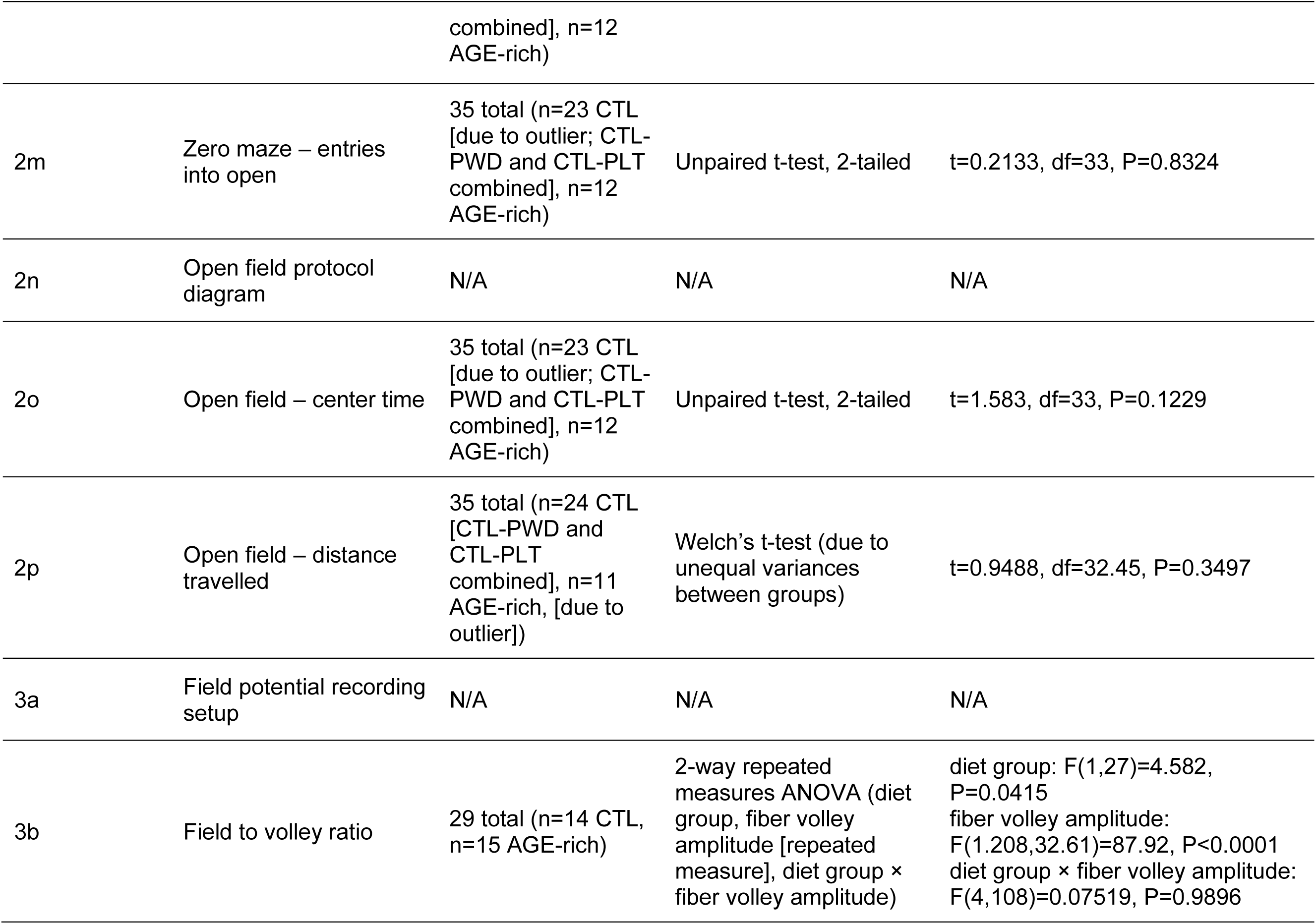
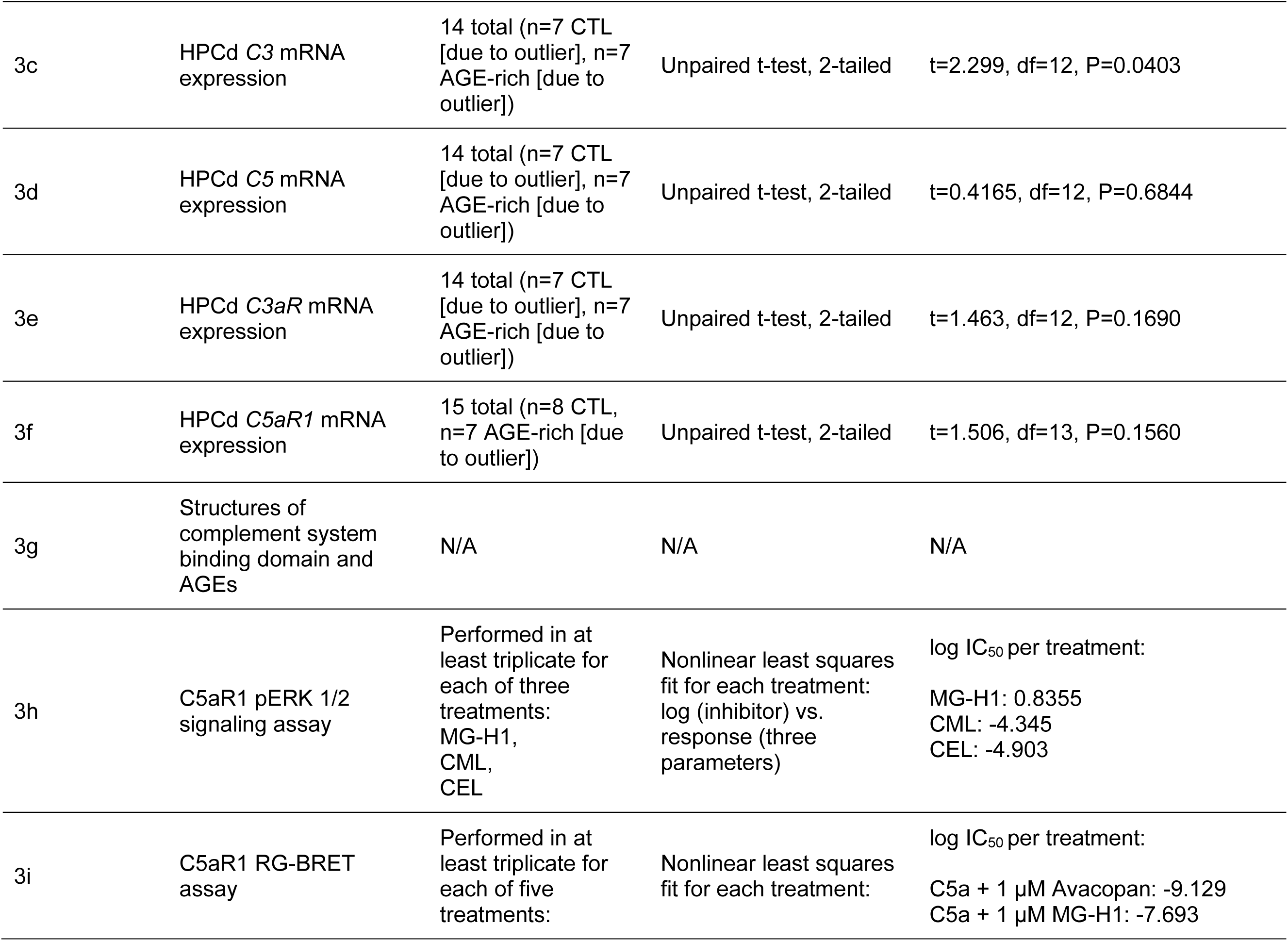
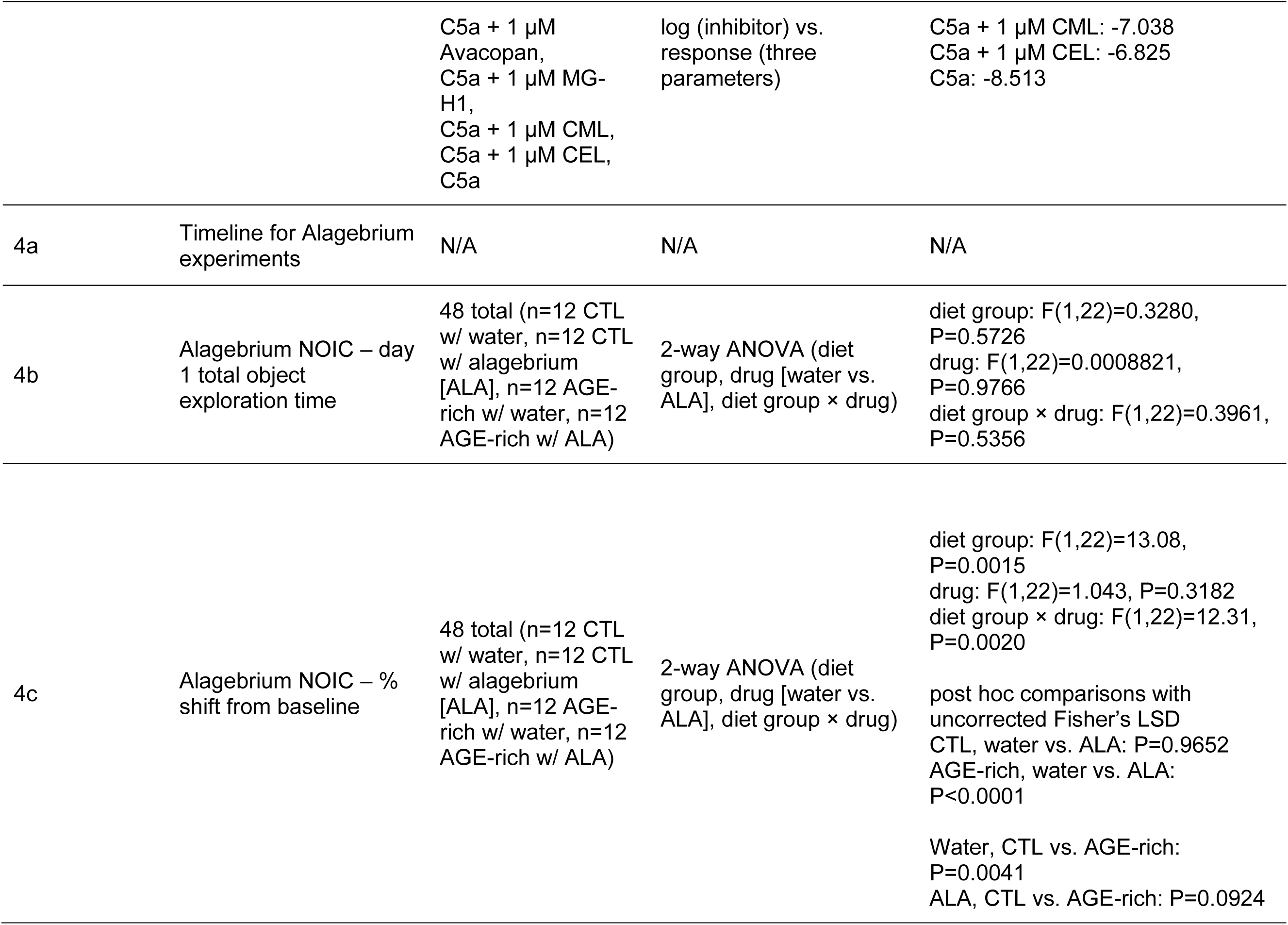
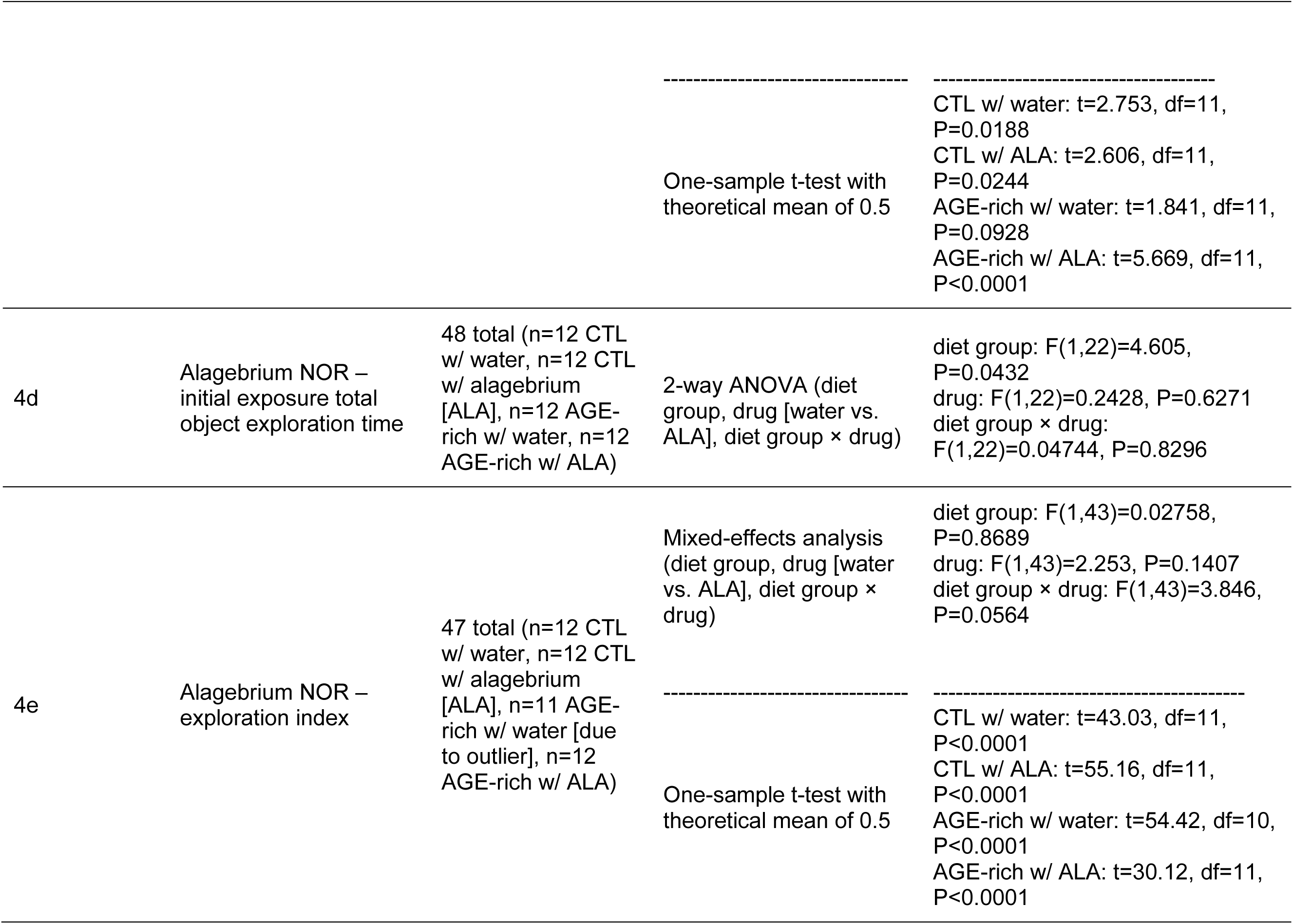
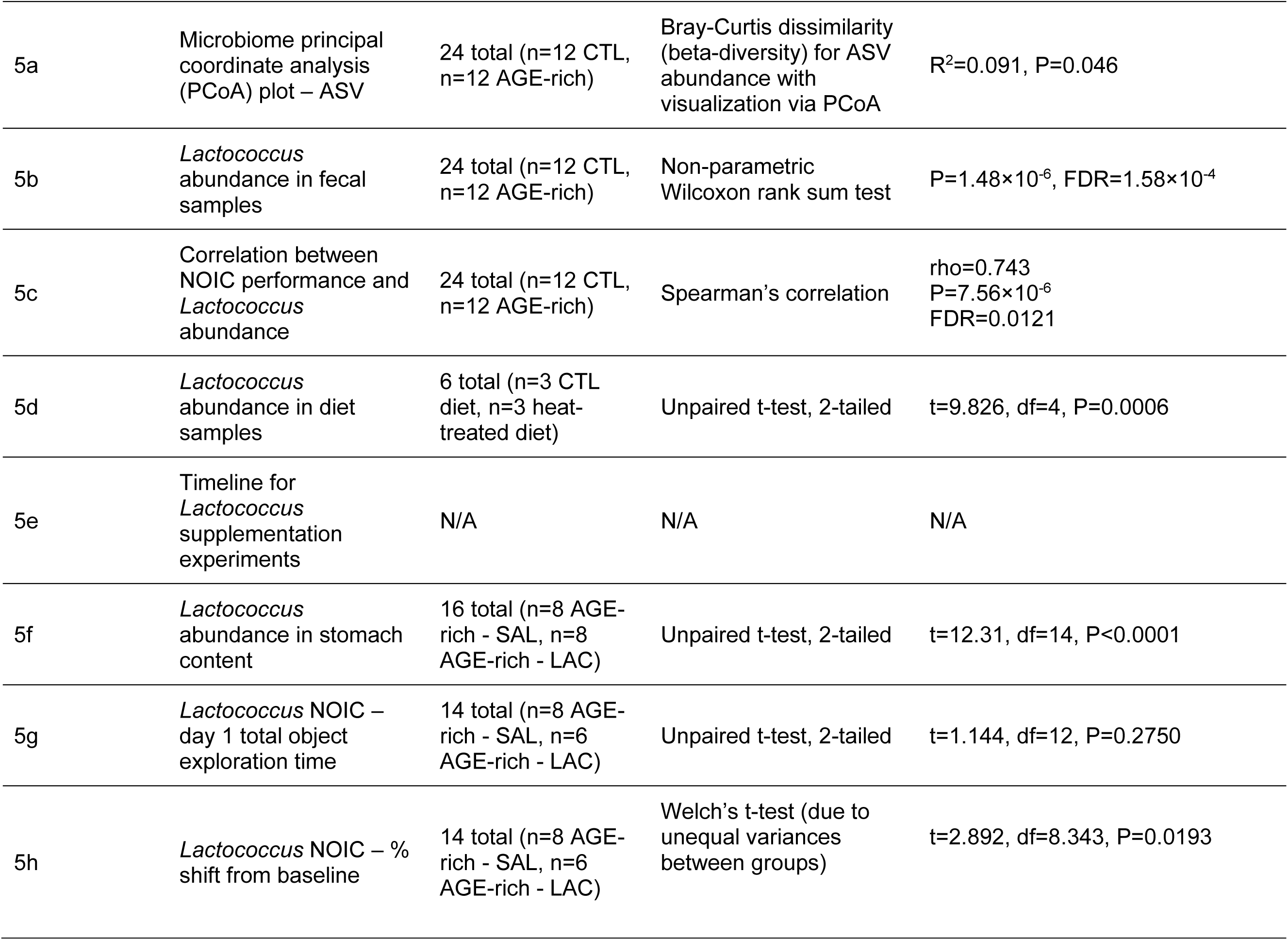
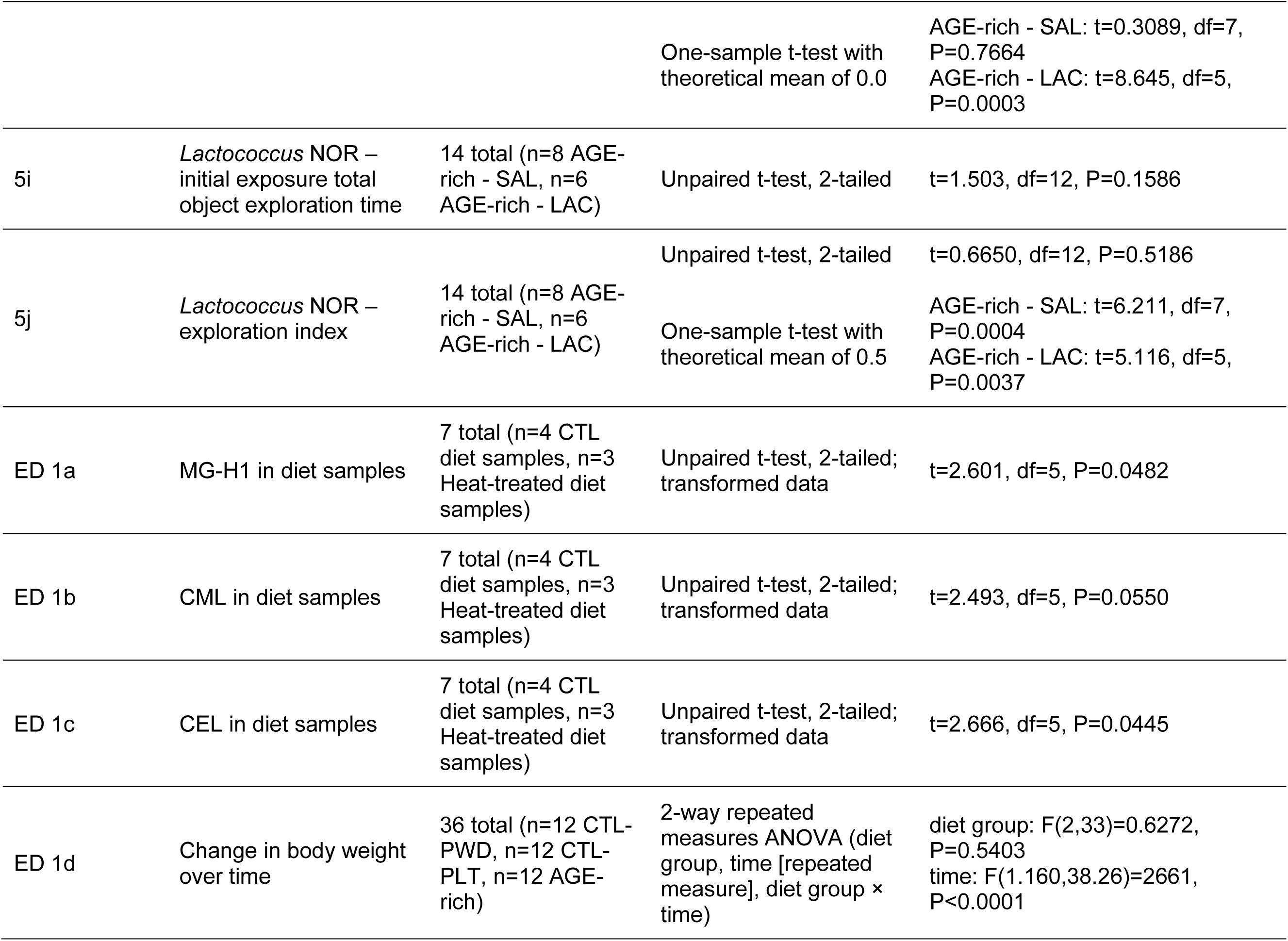
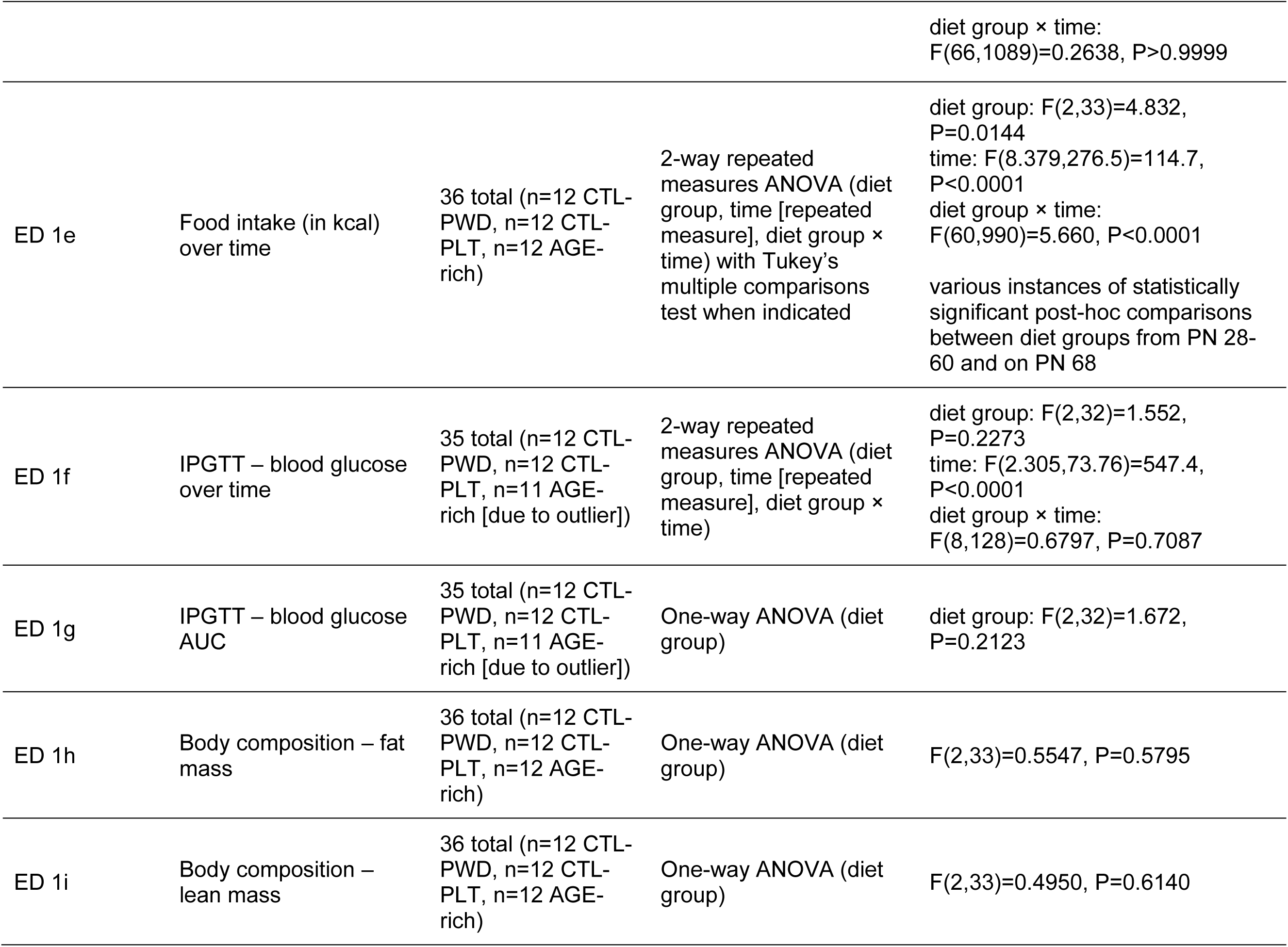
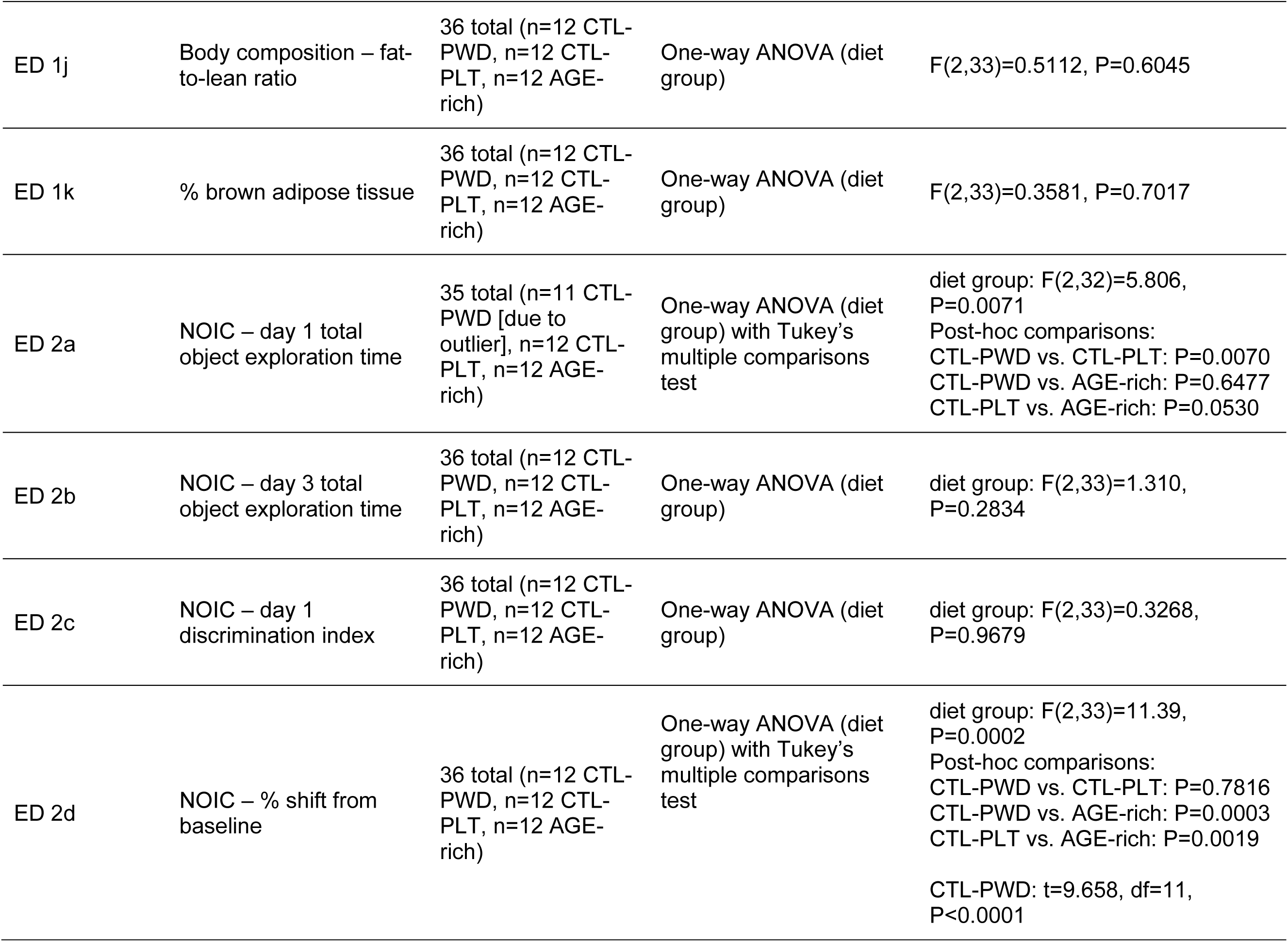
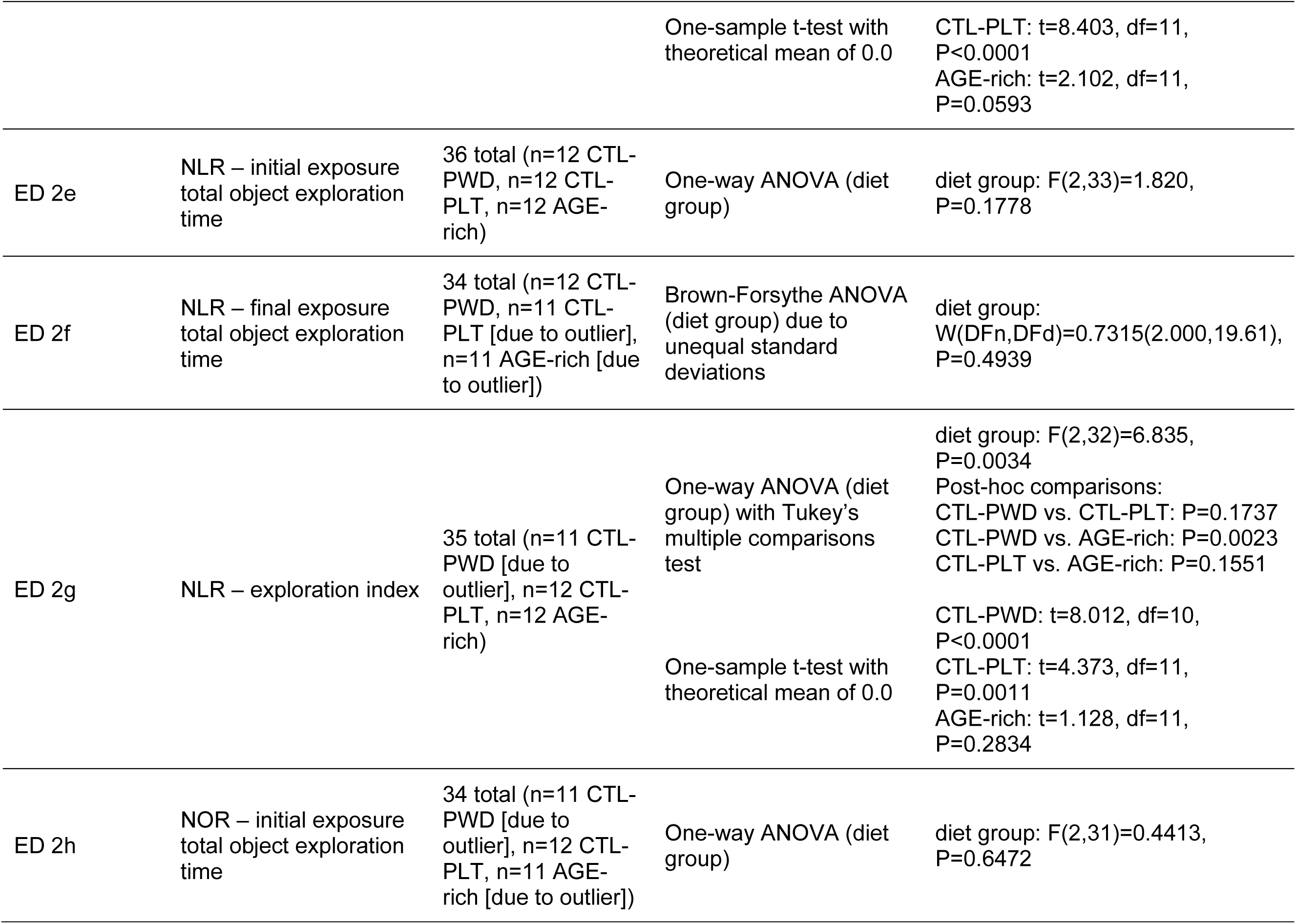
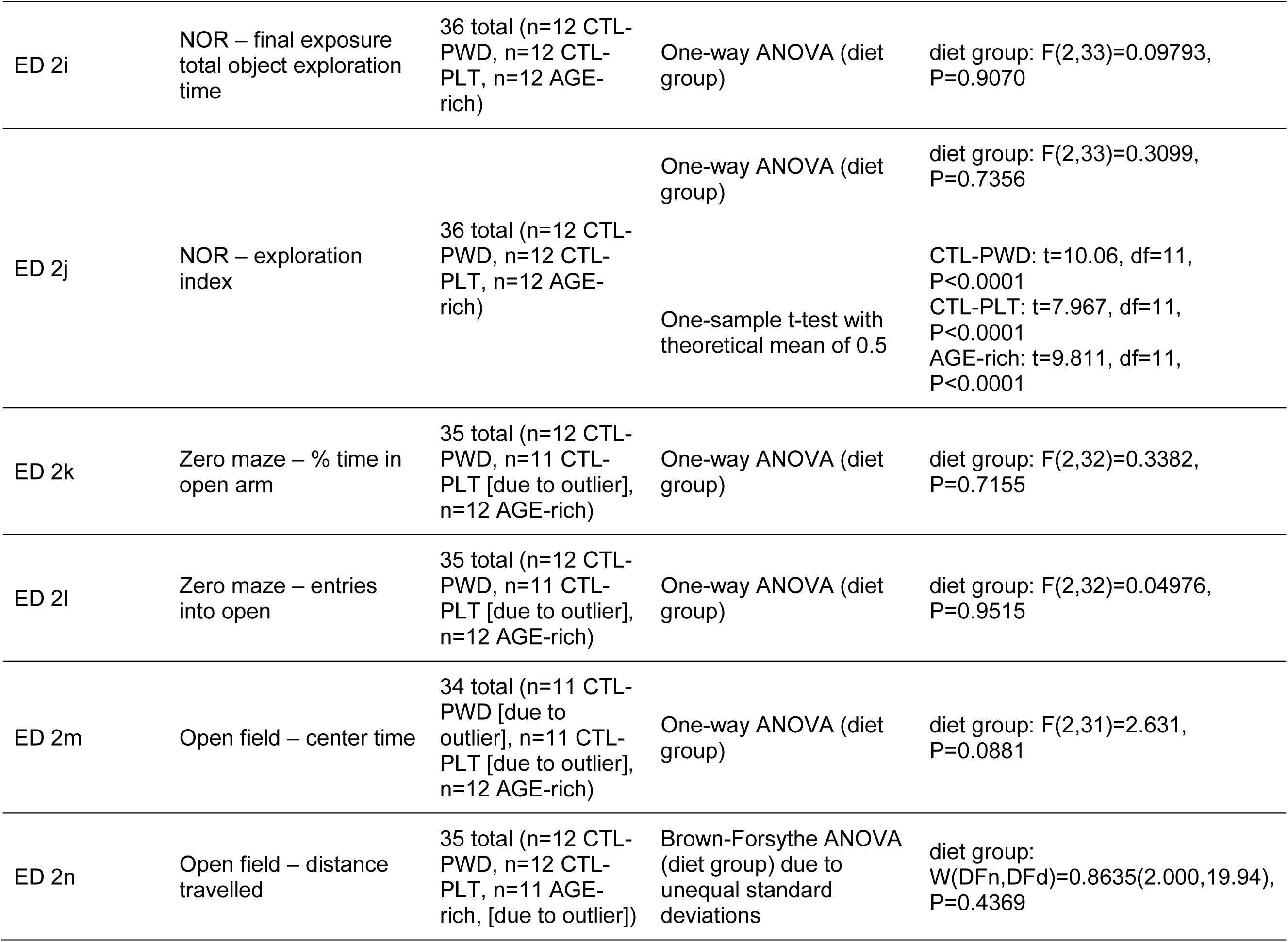
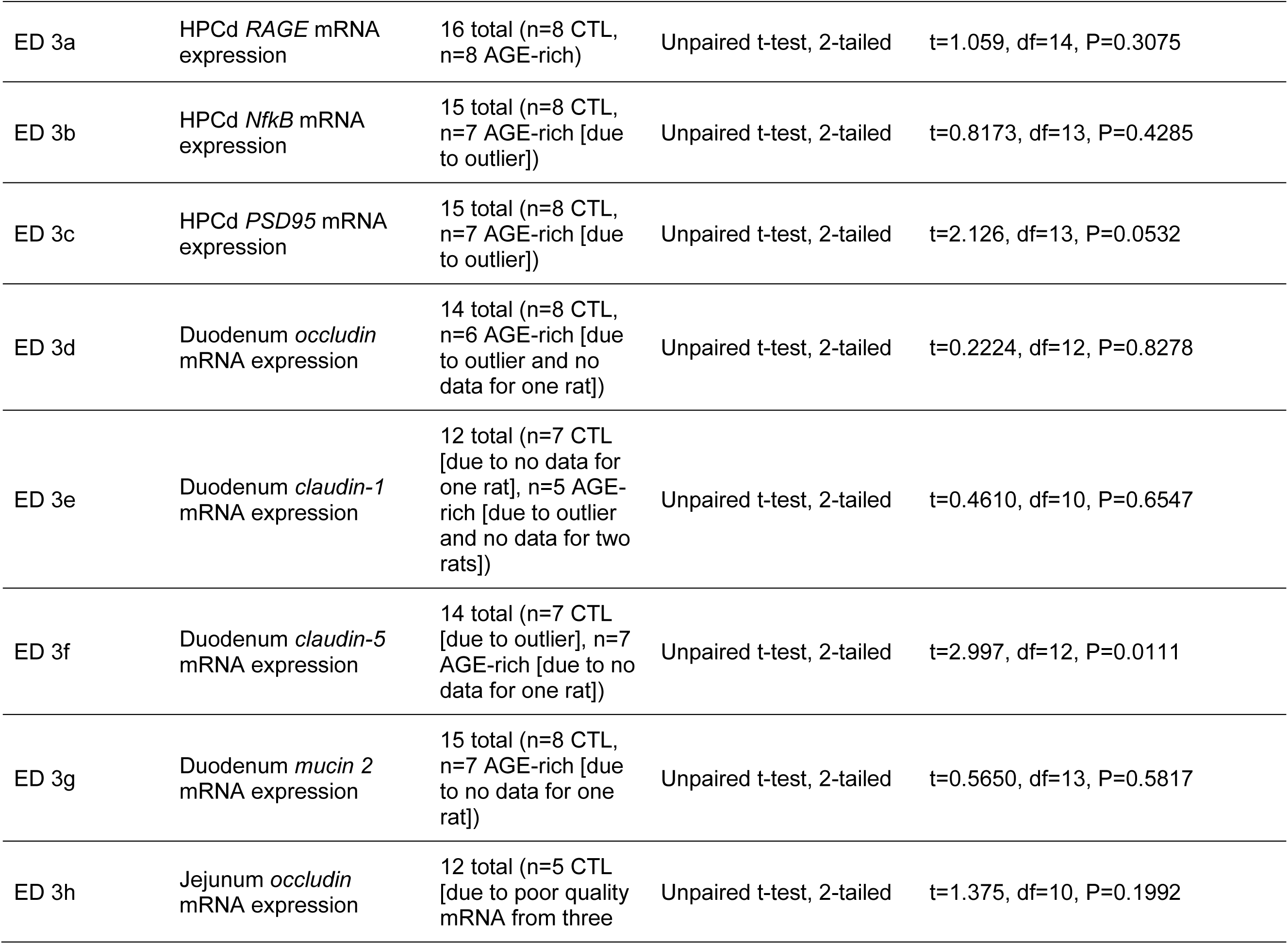
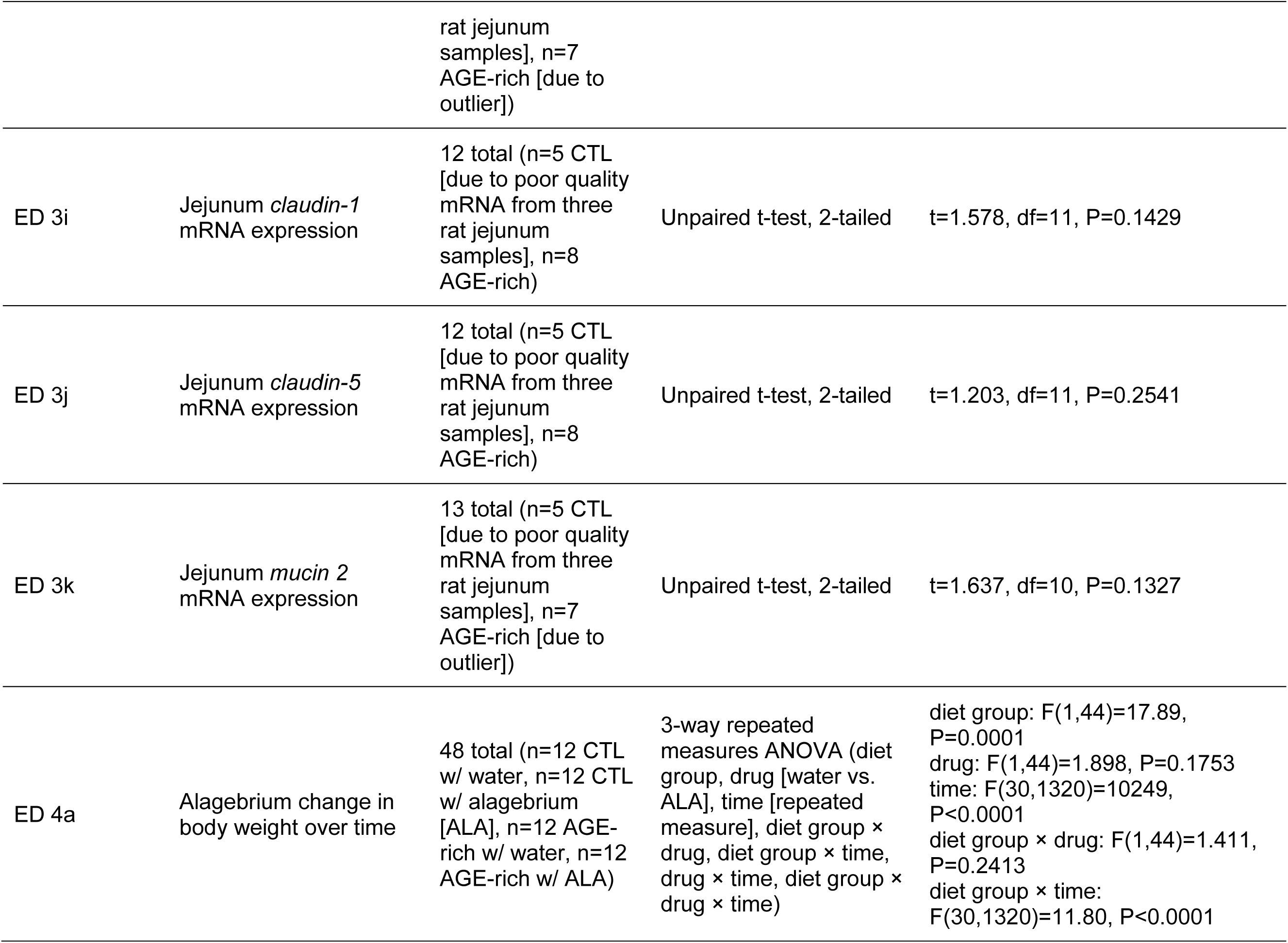
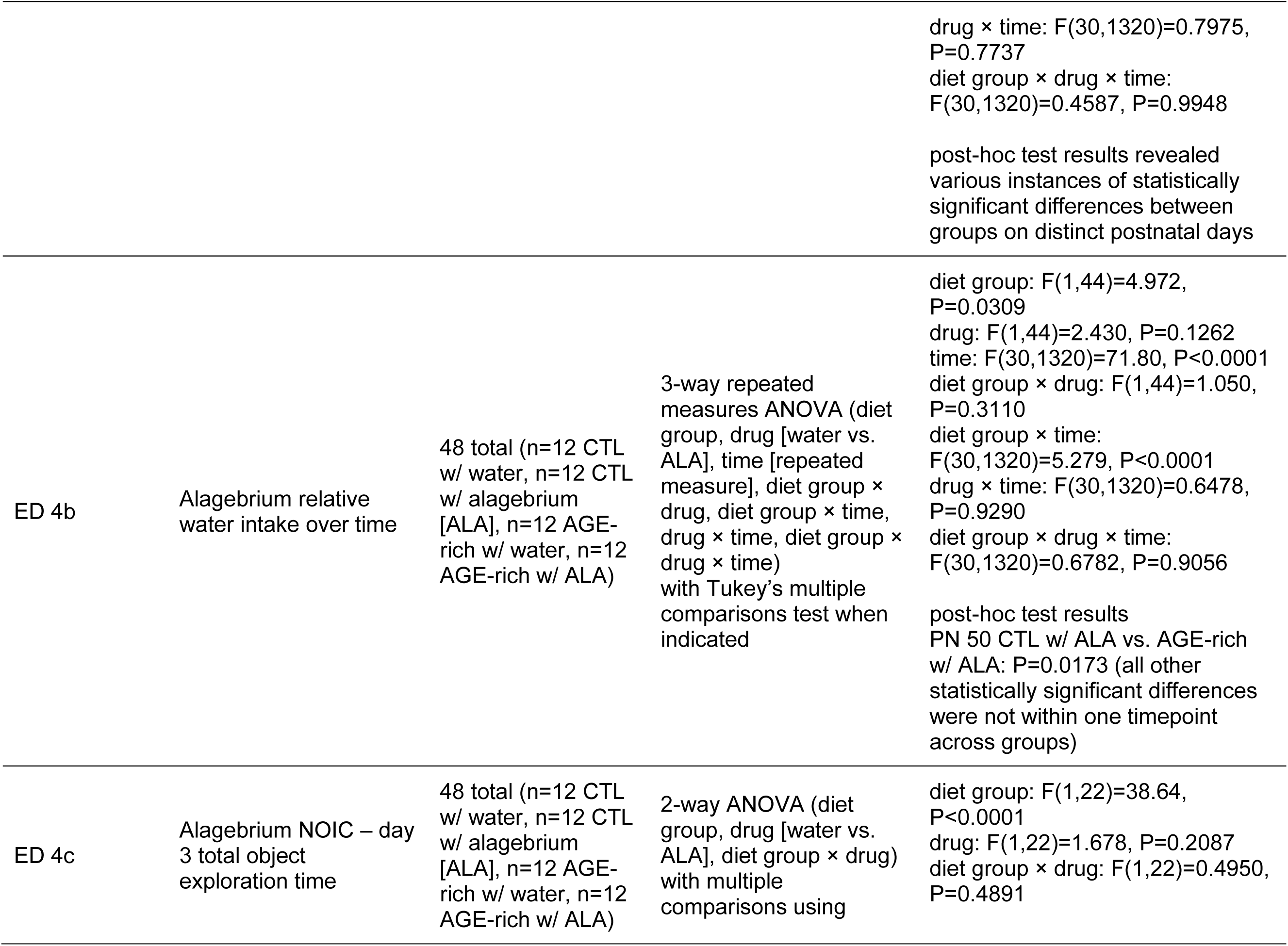
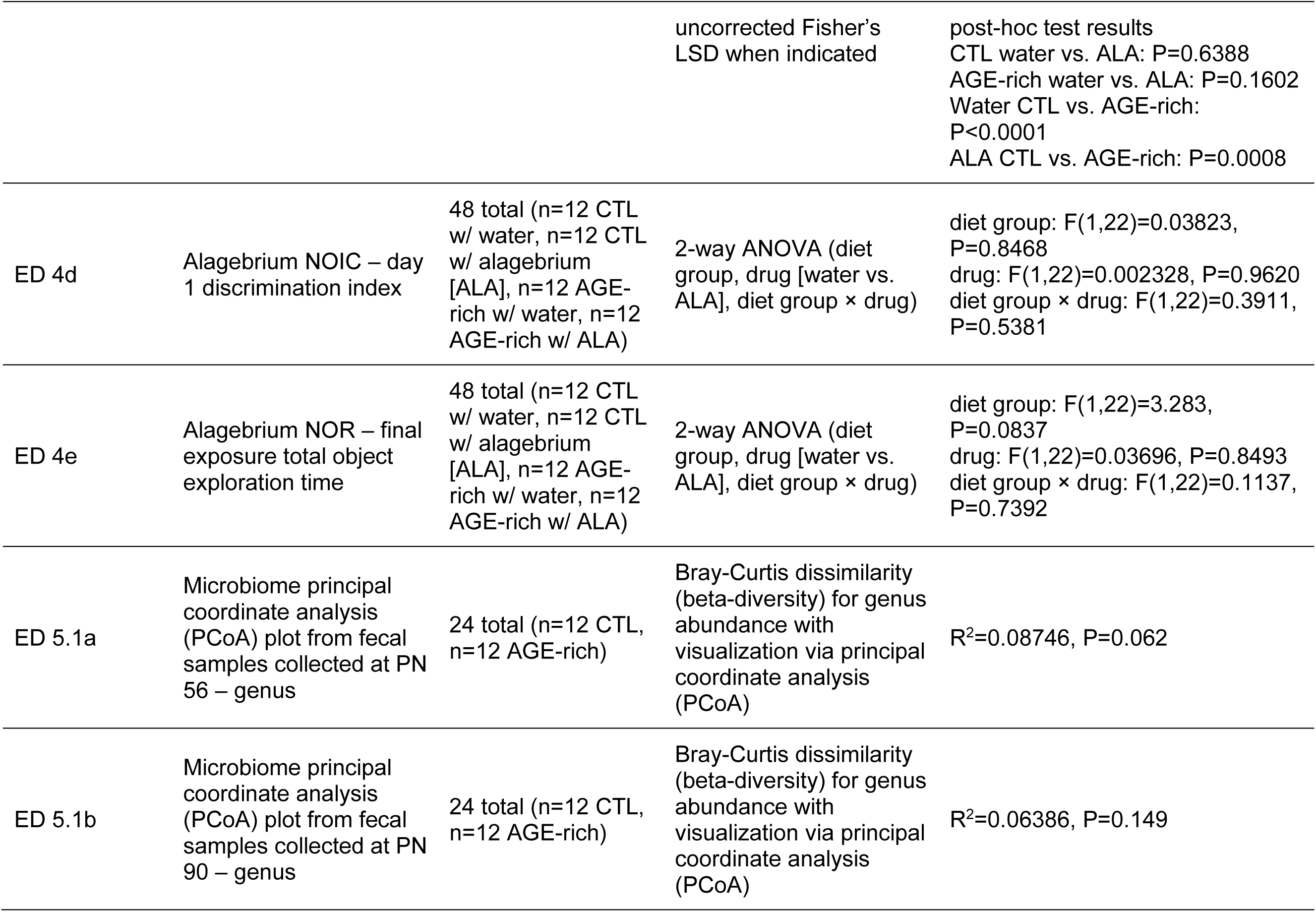
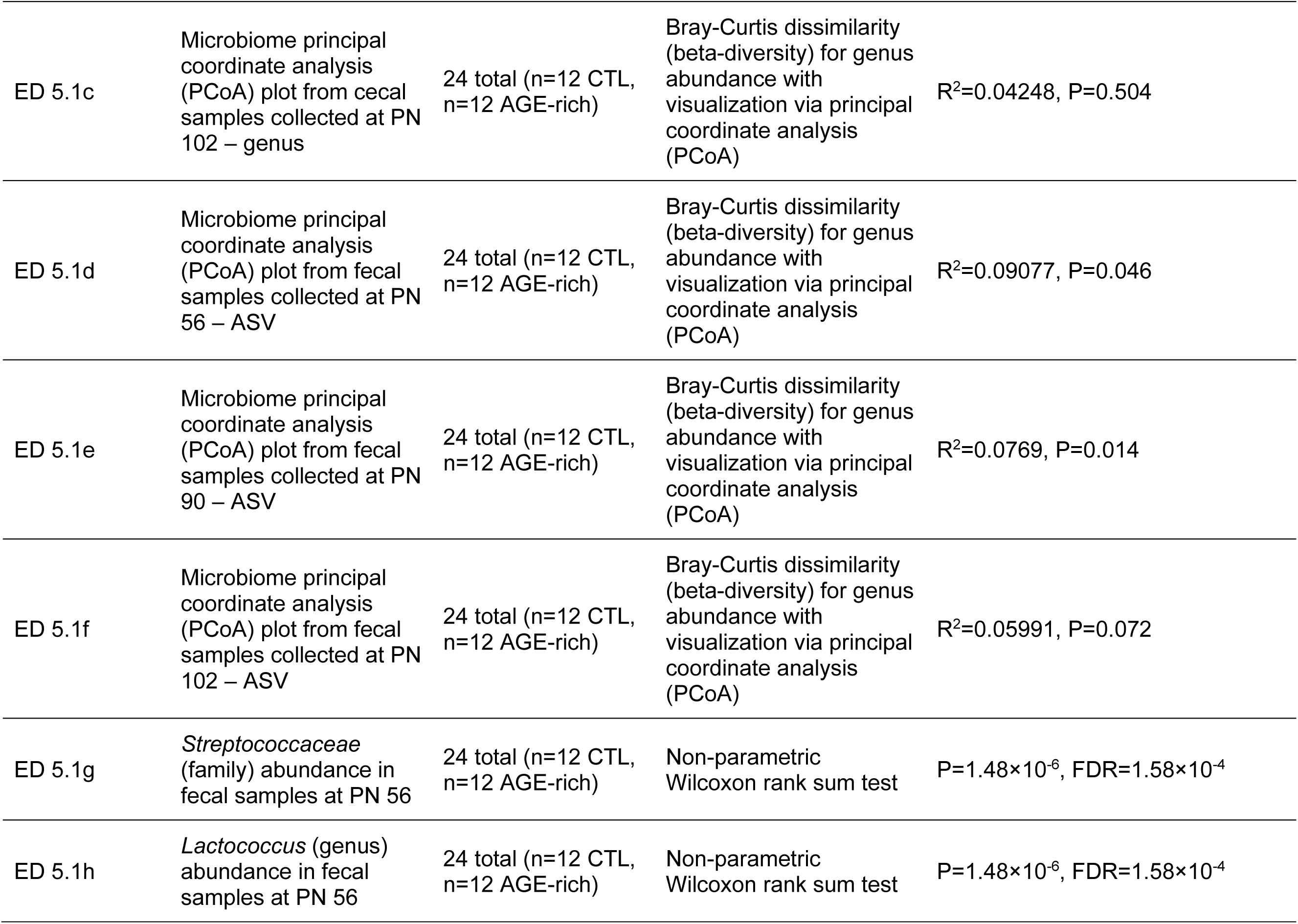
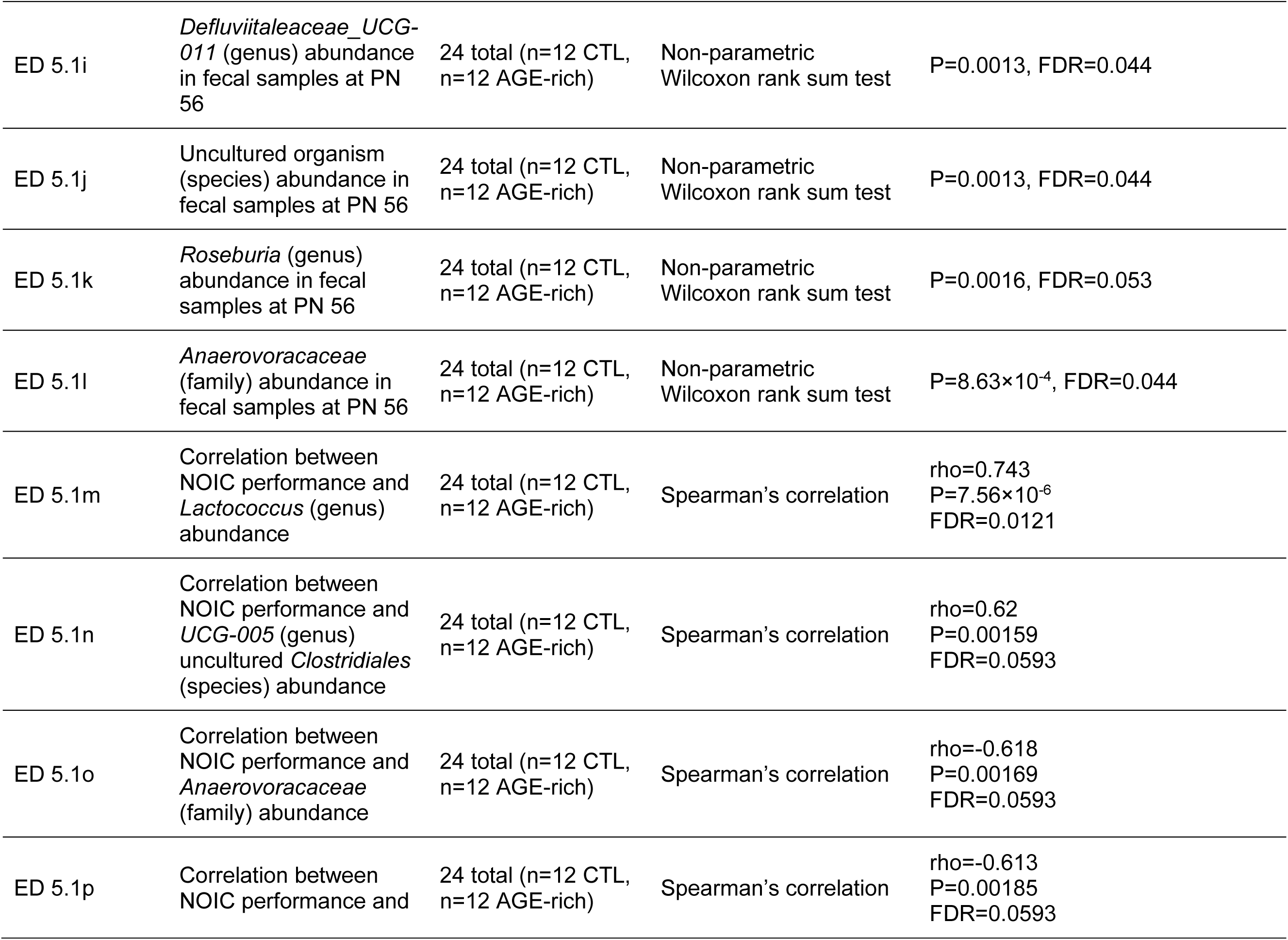
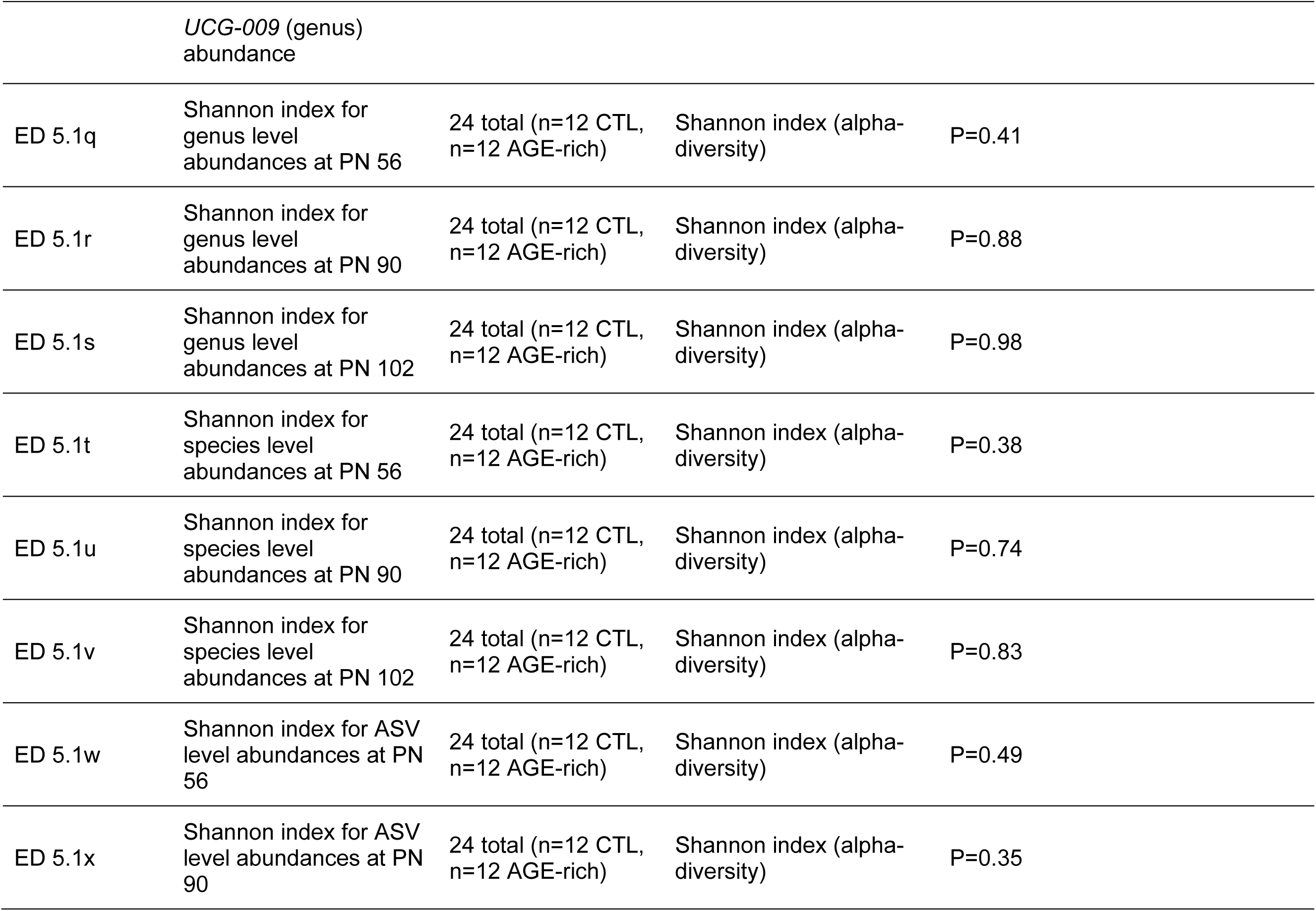
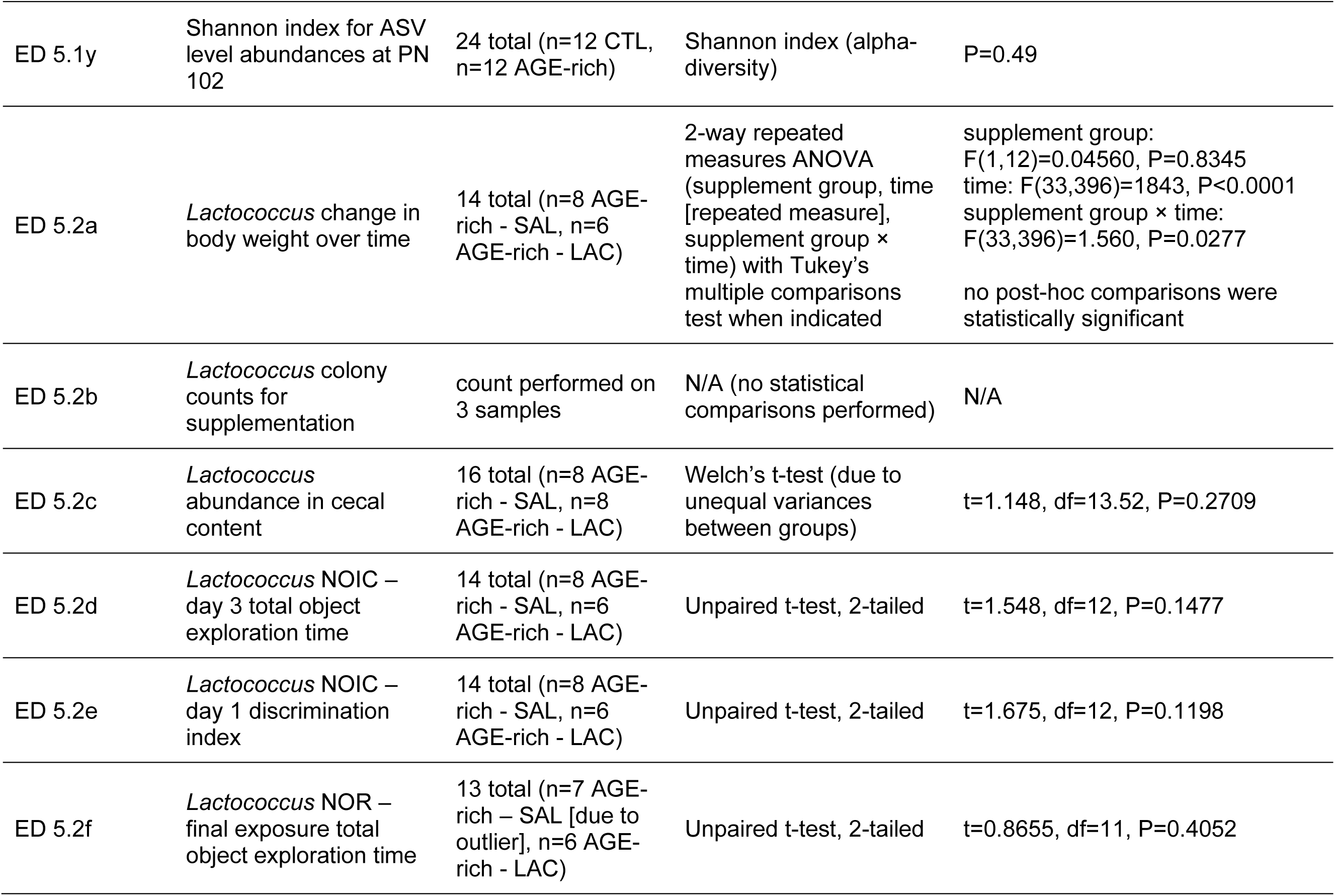
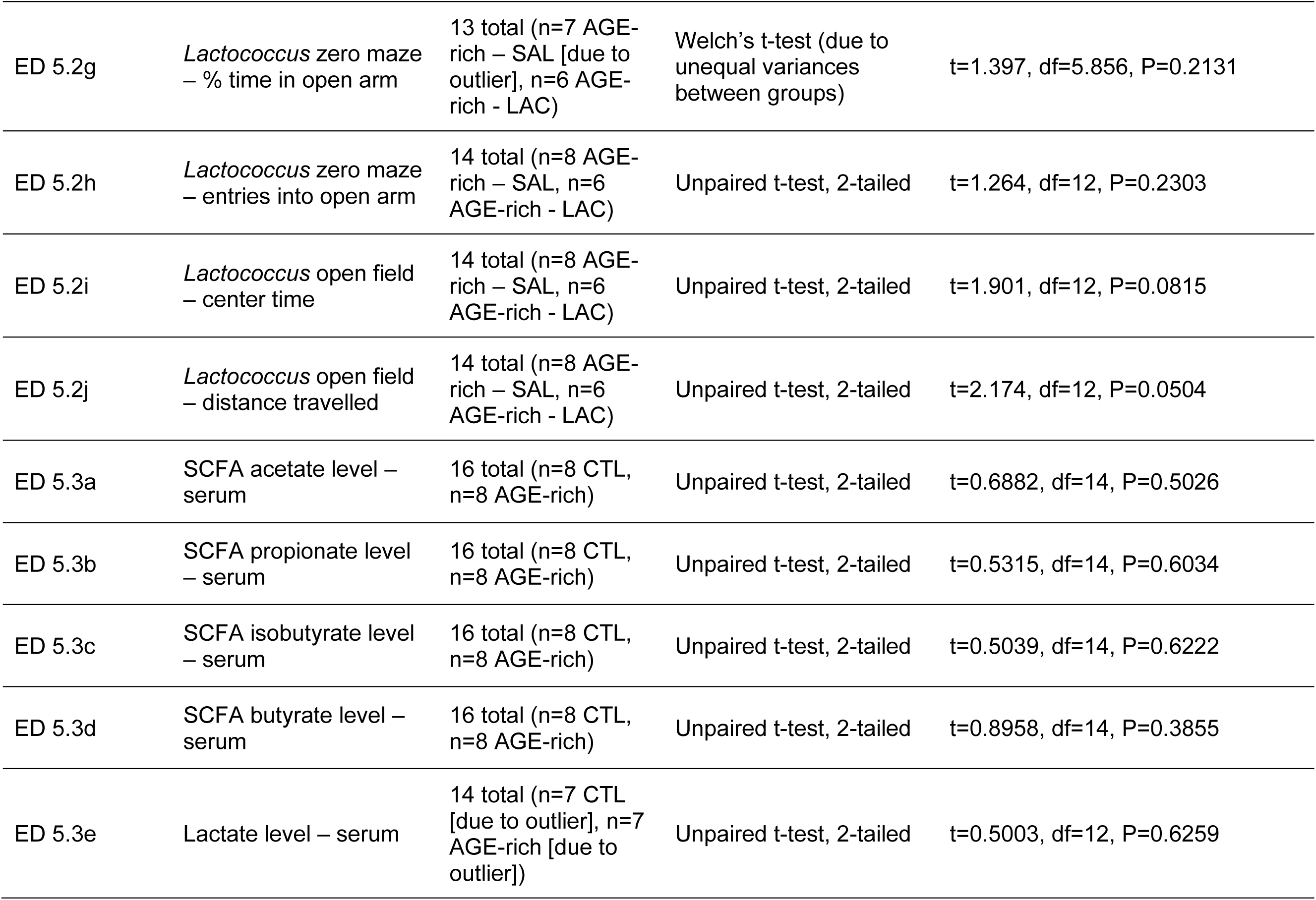
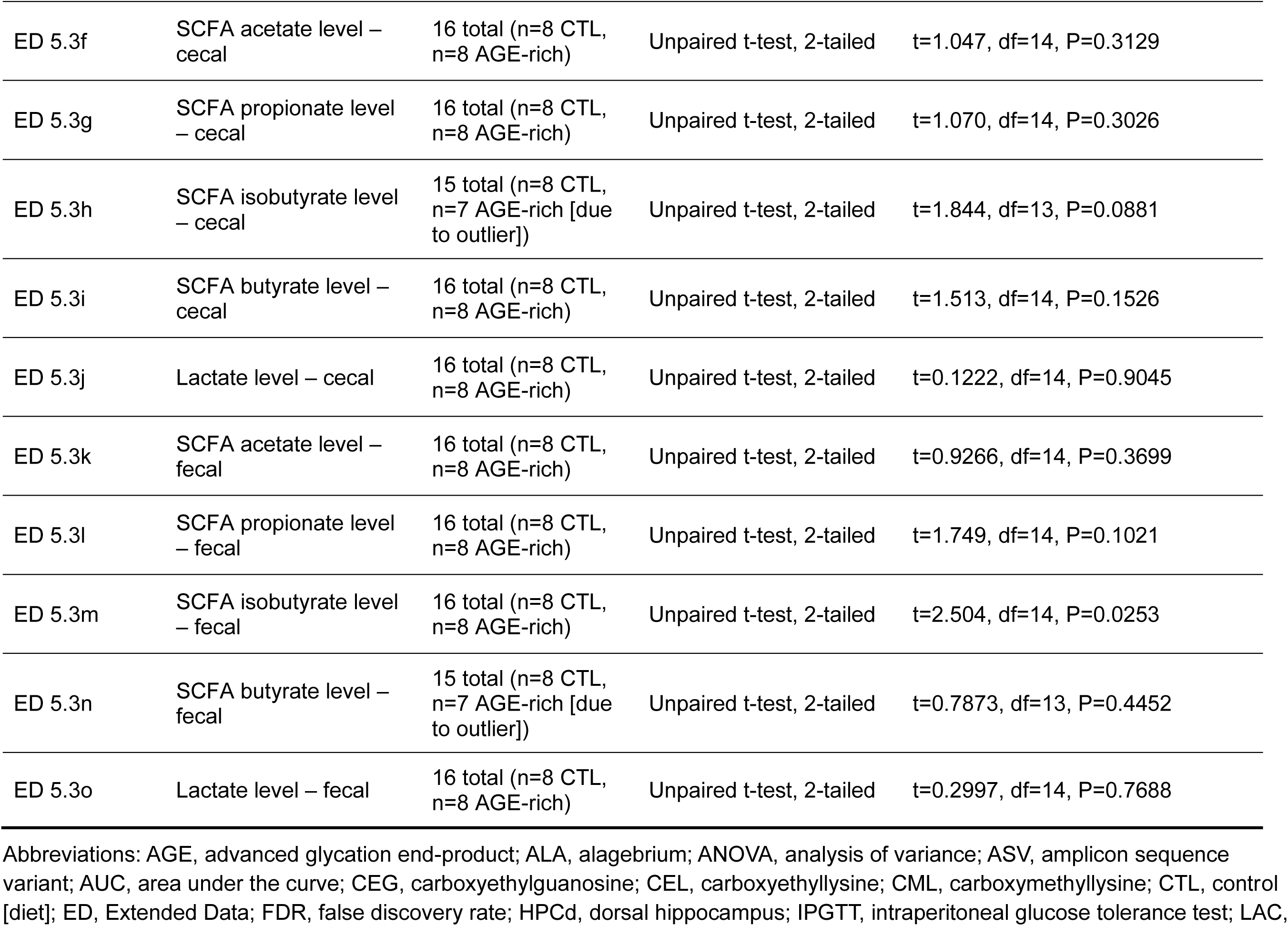
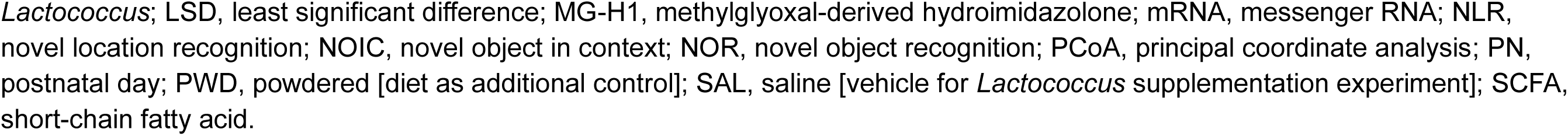
Number of subjects/samples and statistical analyses used per experiment, summarized per figure subpanel.

